# Complex polymorphic genetic architecture underlies *trans*-regulatory variation among strains of *Saccharomyces cerevisiae*

**DOI:** 10.1101/566653

**Authors:** Brian P. H. Metzger, Patricia J. Wittkopp

**Author notes:** Corresponding author: Patricia J. Wittkopp, Department of Ecology and Evolutionary Biology, Department of Molecular, Cellular, and Developmental Biology University of Michigan, Ann Arbor, MI 48109-1048 USA.

## Abstract

Heritable variation in gene expression is common within species. Much of this variation is due to genetic changes at loci other than the affected gene and is thus *trans*-acting. This *trans*-regulatory variation is often polygenic, with individual variants typically having small effects, making the genetic architecture of *trans*-regulatory variation challenging to study. Consequently, key questions about *trans*-regulatory variation remain, including how selection affects this variation and how *trans*-regulatory variants are distributed throughout the genome and within species. Here, we show that *trans*-regulatory variation affecting *TDH3* promoter activity is common among strains of *Saccharomyces cerevisiae*. Comparing this variation to neutral models of *trans*-regulatory evolution based on empirical measures of mutational effects revealed that stabilizing selection has constrained this variation. Using a powerful quantitative trait locus (QTL) mapping method, we identified ∼100 loci altering expression between a reference strain and each of three genetically distinct strains. In all three cases, the non-reference strain alleles increased and decreased *TDH3* promoter activity with similar frequencies, suggesting that stabilizing selection maintained many *trans*-acting variants with opposing effects. Loci altering expression were located throughout the genome, with many loci being strain specific and others being shared among multiple strains. These findings are consistent with theory showing stabilizing selection for quantitative traits can maintain many alleles with opposing effects, and the wide-spread distribution of QTL throughout the genome is consistent with the omnigenic model of complex trait variation. Furthermore, the prevalence of alleles with opposing effects might provide raw material for compensatory evolution and developmental systems drift.

**Significance statement:** Gene expression varies among individuals in a population due to genetic differences in regulatory components. To determine how this variation is distributed within genomes and species, we used a powerful genetic mapping approach to examine multiple strains of *Saccharomyces cerevisiae*. Despite evidence of stabilizing selection maintaining gene expression levels among strains, we find hundreds of loci that affect expression of a single gene. These loci vary among strains and include similar frequencies of alleles that increase and decrease expression. As a result, each strain contains a unique set of compensatory alleles that lead to similar levels of gene expression among strains. This regulatory variation might form the basis for large scale regulatory rewiring observed between distantly related species.

## Introduction

Heritable variation in gene expression results from genetic variation affecting *cis*-regulatory elements (e.g., promoters, enhancers) and *trans*-acting factors (e.g., proteins, RNAs). These *trans*-regulatory changes are located throughout the genome and are the major source of regulatory variation within species (1–10). The number, identity, and effects of individual loci contributing to variation in gene expression has been determined in a variety of species using expression quantitative trait locus (eQTL) mapping (11–18), with the most extensive dissection of eQTL coming from studies of two strains of the baker’s yeast *Saccharomyces cerevisiae* (19–28). These studies have found that (i) expression differences are typically associated with ∼10 or fewer eQTL, (ii) most eQTL have individually small effects on expression, and (iii) most eQTL are located far from the gene whose expression they affect and thus likely contribute to *trans*-regulatory differences.

Traditional eQTL mapping approaches require genotype and expression data for many individuals to detect significant effects. Consequently, studies mapping the genetic basis of regulatory differences have largely been limited to two strains or populations within any given species. In cases where polymorphism of regulatory variation has been studied within a species, experiments have focused on *cis*-regulatory variation for technical reasons (29–34). As a result, key questions about the extent and genetic basis of *trans*-regulatory variation segregating within a species remain unanswered. For example, do multiple *trans*-regulatory variants affecting a gene’s expression often segregate at the same locus within a species? How different are the suites of *trans*-acting eQTL affecting a gene’s expression among individuals or strains? Are the effects of *trans*-regulatory variants at different loci often in the same direction, or do they typically have opposing effects, canceling one another out? Addressing these questions requires identifying *trans*-acting eQTL and their effects on expression among multiple individuals or strains of the same species.

In addition to these questions about the variability in genetic architecture of *trans*-regulatory variation, questions also remain about the impact of selection on this variation. Prior work has shown that gene expression levels are broadly constrained by stabilizing selection (35, 36), and variation in *cis*-regulatory eQTL appears to be limited by purifying selection (30, 37). But the impact of natural selection on the number, identity, or genomic distribution of *trans*-acting eQTL is less clear, and there are reasons to suspect that it might be different than for *cis*-acting eQTL. For example, prior work suggests that *trans*-regulatory mutations arise more frequently than *cis*-regulatory mutations, but tend to have smaller effects on the focal gene’s expression (38). In addition, *trans*-regulatory mutations are more likely to be recessive and have greater pleiotropic effects than *cis*-regulatory mutations (39–43). Any or all of these factors might cause selection for the level of gene expression to have different impacts on *cis*- and *trans*-regulatory variation.

Here, we examine *trans*-regulatory variation segregating among genetically distinct strains of *S. cerevisiae*. We focus on the extent of, genetic basis for, and evolutionary forces acting on, *trans*-regulatory variation affecting expression of the *TDH3* gene, which encodes a glyceraldehyde-3-phosphate dehydrogenase. This gene was chosen because prior work has estimated the effects of new *trans*-regulatory mutations on its expression (38, 40) as well as the fitness consequences of changing its expression (44, 45), allowing us to compare the *trans*-regulatory variation segregating in *S. cerevisiae* to empirically-informed models of neutral evolution. We find that although differences in *trans*-regulation affecting *TDH3* promoter activity are common among strains, they generate less variation in *TDH3* promoter activity than predicted by neutral models, confirming that stabilizing selection has acted on *trans*-regulatory variation affecting *TDH3* promoter activity in the wild. We then use a powerful genetic mapping approach to determine differences in the genetic architecture of this *trans*-regulatory variation by identifying eQTL between each of three strains of *S. cerevisiae* and a common reference strain. In each of these three eQTL mapping experiments, we find ∼100 eQTL affecting activity of the *TDH3* promoter in *trans*. These loci are often different among strains, have opposing effects on expression, and are spread throughout the genome, indicating diverse sources of *trans*-regulatory variation segregating within *S. cerevisiae*. These results agree with theoretical predictions that stabilizing selection can maintain genetic variation for polygenic traits (46–50). They also suggest that natural populations harbor greater regulatory variation than suggested by differences among strains, which can impact the evolution of regulatory systems.

## Results and Discussion

To isolate the effects of *trans*-regulatory variants segregating among *S. cerevisiae* strains on *TDH3* promoter activity, we inserted a Yellow Fluorescent Protein (YFP) coding region under control of a common *TDH3* promoter from the BY lab strain into 56 distinct *S. cerevisiae* strains (Figure 1A). These strains (i) were isolated from a range of environments, (ii) differ at more than 100,000 SNPs and small indels, many larger CNVs and chromosomal rearrangements, and (iii) encompass much of the genetic and phenotypic diversity observed within the species (51, 52). For each strain, we measured YFP fluorescence in twelve biological replicate populations grown in rich media and used the measured YFP fluorescence to estimate changes in *TDH3* mRNA levels due to differences in *trans*-regulation among strains. We observed that *trans*-regulatory variation caused differences in expression that ranged from 71% to 147% of the reference strain (Figure 1B, C). This variance in *trans*-regulation was nearly double the variance of *cis*-regulation described among a similar set of strains (Figure S1A)(53). We detected significant phylogenetic structure for *trans*-regulatory differences among strains, with more closely related strains having on average more similar *TDH3* promoter activity than more distantly related strains (λ= 0.59, p = 0.013; K=0.49, p=0.012; Figure 1D).

**Figure 1.**
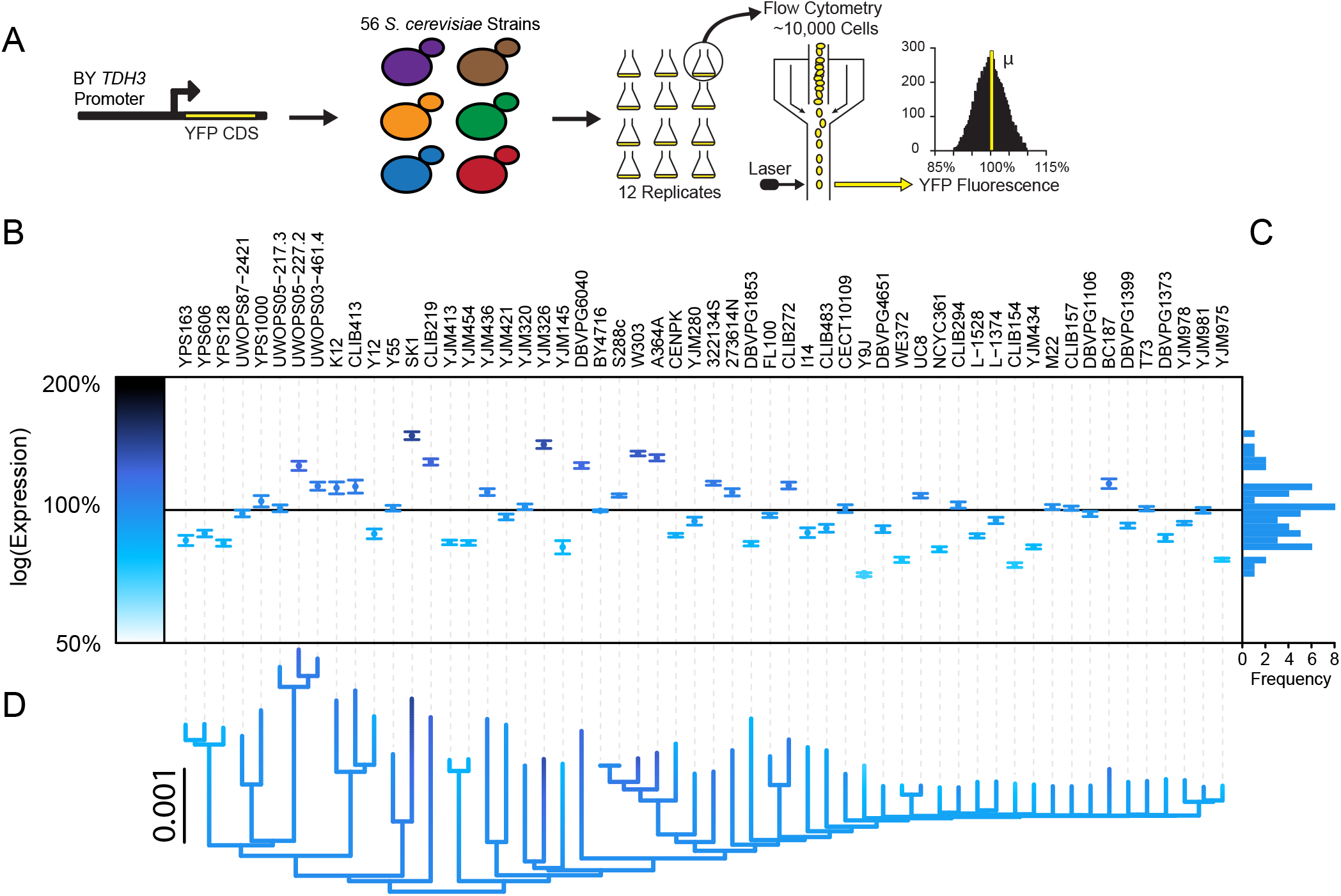
Extensive *trans*-regulatory variation affecting *TDH3* expression is segregating among *S. cerevisiae* strains. A. Variation in *TDH3 trans*-regulatory backgrounds among yeast strains was measured using a reporter gene containing the *TDH3* promoter from the BY strain and a yellow fluorescent protein (YFP). This reporter was integrated into the genome of 56 diverse *S. cerevisiae* strains. Twelve replicate populations were grown in YPD and analyzed by flow cytometry for YFP expression. B. Variation among replicates relative to the BY reference strain was used to calculate the average effect of each strain’s *trans*-regulatory background on *TDH3* promoter activity. Darker colors reflect higher *TDH3* reporter activity. C. Frequency of *trans*-regulatory effects relative to reference strain. D. Phylogenetic relationships among strains as estimated from genome wide polymorphism data (51). Color of branches corresponds to estimated *trans*-regulatory effect from ancestral character estimation.

To determine how natural selection has impacted this *trans*-regulatory variation, we constructed models of neutral evolution and compared these models to the observed differences in *TDH3* promoter activity among strains. We simulated the neutral evolution of *trans*-regulatory variation affecting *TDH3* promoter activity by sampling *trans*-regulatory mutational effects on *TDH3* promoter activity defined in prior work (Figure 2A) (38), and tracking how expression changed with the addition of each new mutation (Figure 2B). We repeated this sampling process 10,000 times and used the observed distributions of expression levels after the addition of each new mutation to define the probability with which we expect to see a given expression level evolve neutrally from the common ancestor after a particular number of genetic changes. We then compared this neutral projection to the observed differences in *TDH3* promoter activity among strains, using the genetic relationships among strains to infer how *TDH3* promoter activity changed along each branch of the phylogeny (Figure 2C). We found that there was significantly less *trans*-regulatory variability among strains than predicted to arise from mutation alone, suggesting that natural selection has constrained *TDH3* promoter activity (p < 0.0001, Figure S1B, C). These simulations do not, however, account for epistatic interactions among new regulatory mutations, for which no current data exists (SI Methods).

**Figure 2.**
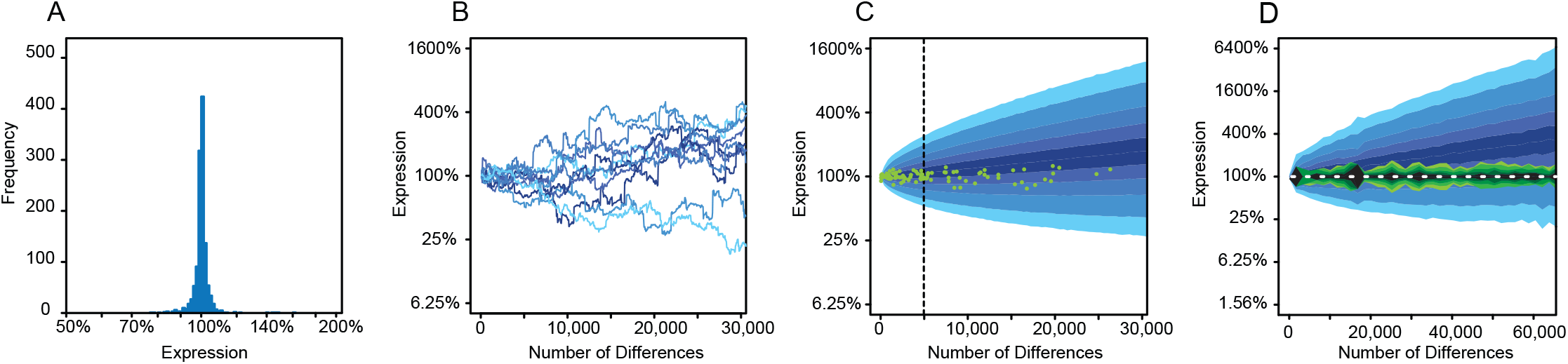
Natural selection has constrained *TDH3 trans*-regulatory variation. A. Effects of *trans*-regulatory mutations on *TDH3* promoter activity. Mutants were collected and analyzed in prior work (38). B. Simulated neutral trajectories for *TDH3* promoter activity based on empirically measured effects of new mutations. Lighter colors reflect more extreme values after 30,000 mutations. C. Comparison of observed differences in *TDH3* promoter activity among *S. cerevisiae* strains with neutral expectation. The blue background represents the 95th, 90th, 80th, 70th, and 60^th^ percentiles, from light to dark, for the simulated neutral trajectories. Green dots are differences in *TDH3* promoter activity and estimated number of mutations based on the *S. cerevisiae* phylogeny. Dashed line indicates the point where the observed data departs significantly from expectation. D. Same as C, but using genetic distance instead of phylogenetic distance between strains. The green areas represent the 95th, 90th, 80th, 70th, and 60^th^ percentiles, from light to dark, for the observed differences from sampling pairs of strains.

We next tested whether our inference of stabilizing selection was robust to uncertainty in the inferred phylogenetic relationships among strains and the inferences of changes in *TDH3* promoter activity on the phylogeny by repeating this analysis using the total genetic distance between pairs of strains instead of the phylogenetic relationships among strains. We again found less *trans*-regulatory variability in *TDH3* promoter activity among strains than predicted by the neutral model, further supporting the hypothesis that *trans*-regulatory variation affecting *TDH3* promoter activity has evolved under stabilizing selection (Figure 2D, Figure S2D). Next, we tested for evidence of natural selection acting on *TDH3 trans*-regulatory variation using an approach that does not rely on empirical estimates of the effects of new *trans*-regulatory mutations. Specifically, we fit the *P_TDH3_-YFP* reporter activity and phylogenetic relationships among strains to two models of quantitative trait evolution: a Brownian Motion model of neutral quantitative trait evolution and an Ornstein-Ulenbeck model that incorporates stabilizing selection (54). We found that the Ornstein-Ulenbeck model fit the data significantly better than the neutral Brownian motion model (p = 0.00007, chi-square test, Figure S2E, F), again suggesting that *trans*-regulatory variation affecting *TDH3* promoter activity in *S. cerevisiae* has been shaped by stabilizing selection. Consistent with these results, the effects of *trans*-regulatory variation on *TDH3* promoter activity are predicted to decrease fitness by less than 0.1% in 80% of strains by prior work describing the relationship between *TDH3* expression level and fitness in rich media (45), with the largest deviation in *TDH3* promoter activity (71% of wild type) expected to decrease fitness by only 0.5% (Figure S2G). Taken together, we conclude that stabilizing selection has constrained *trans*-regulatory variation affecting expression of *TDH3*.

In the presence of stabilizing selection, gene expression can be kept similar among strains by purifying selection purging mutations that alter expression or by maintaining sets of variants with off-setting, or compensatory, effects on expression within the population. To determine which of these mechanisms is more likely to have minimized differences among strains in the *trans*-regulatory effects on *TDH3* promoter activity, we used eQTL mapping to examine the genetic architecture of *trans*-regulatory variation affecting *TDH3* promoter activity in three strains (M22, YPS1000, SK1) relative to a common reference strain (BY). Strain M22 is closely related to BY, with YPS1000 and SK1 more distantly related to BY, M22, and each other (Figure S2A). Each of the three focal strains was individually mated with BY to form an F1 hybrid. Hybrids were then forced through three rounds of sporulation and mating, resulting in haploid individuals that had undergone three rounds of recombination (Figure 3A, Left). The resulting segregants were analyzed for *P_TDH3_-YFP* expression, with the 5% of cells with highest YFP expression collected in one pool and the 5% of cells with lowest YFP expression collected in a second pool using fluorescence assisted cell sorting (FACS). These pools were grown to saturation and sorted two additional times for *P_TDH3_-YFP* expression, for three total rounds of selection (Figure 3A, Middle). Cells from the high and low fluorescence selection pools for each of the three pairs of strains were then sequenced. Genetic variants that do not impact *P_TDH3_-YFP* expression were expected to be found at similar frequencies in the high and low fluorescence pools, whereas genetic variants that affect *P_TDH3_-YFP* expression (plus linked loci) were expected to show significant differences in frequency between the high and low fluorescence pools (Figure 3A, right). This bulk-segregant mapping strategy is similar to the extreme QTL mapping approach described previously (28, 55–60), but with additional rounds of recombination and selection (61). The additional rounds of recombination are expected to better resolve individual eQTL, whereas the additional rounds of selection are expected to enrich the genotypes sampled for extreme phenotypes.

**Figure 3.**
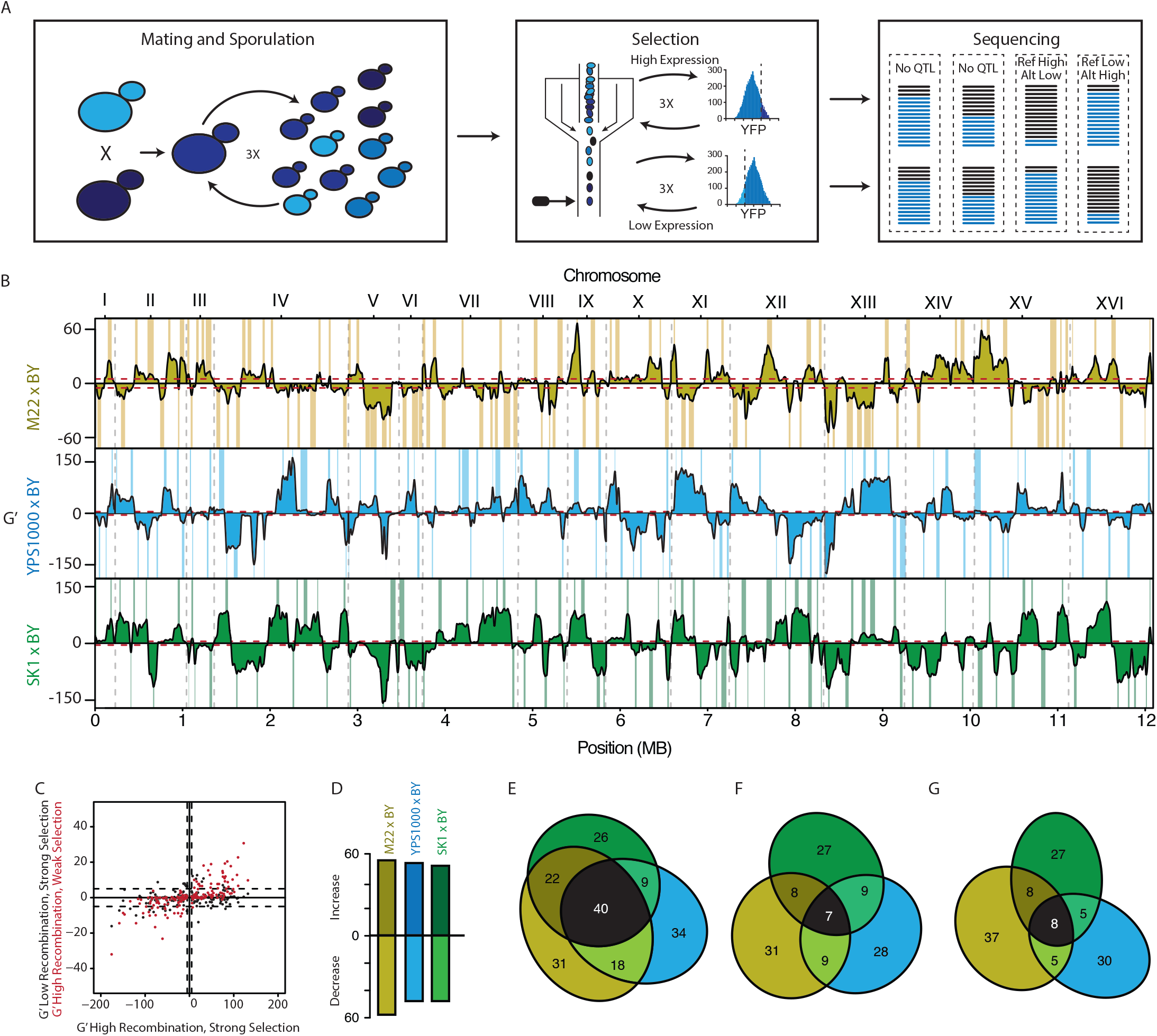
Compensatory alleles underlie the maintenance of *TDH3 trans*-regulatory effects. A. The genomic basis of *TDH3 trans*-regulatory variation was mapped using an xQTL approach. Left: Three rounds of mating and sporulation were used to increase mapping resolution. Middle: Three rounds of FACS based selection were used to enrich for alleles increasing and decreasing *TDH3 trans*-regulatory activity. In each round, the top or bottom 5% of the population was collected. Right: Comparisons of allele frequency from Illumina sequencing of FACS based pools was used to identify eQTL. Each block (dashed lines) represents a different genomic region. Colored lines represent allele frequencies. Black: Reference strain. Blue: Testing strain. For each block, the top bars are after selection for high YFP fluorescence, while the bottom bars are after selection for low YFP fluorescence. eQTL are identified when allele frequencies among the high and low selected pools differ significantly. B. G’ statistic for evidence of eQTL in each comparison. Effects are relative to the non-BY reference allele. Dashed gray lines indicate chromosome boundaries. Dashed red lines gives threshold for statistical significance. Called eQTL with 95% confidence intervals on the location are highlighted for each strain. Brown: M22 x BY. Blue: YPS1000 x BY. Green: SK1 x BY. C. Relationship between G’ statistic for different mapping procedures. X-axis, G’ statistic for high recombination and strong selection (three rounds of crossing and three rounds of selection). Y-axis, (Black) G’ statistic for low recombination and strong selection (one round of crossing and three rounds of selection). (Red) G’ statistic for high recombination and weak selection (three rounds of crossing and one round of selection). Each point is for an eQTL identified with high recombination and strong selection (three rounds of crossing and three rounds of selection) from the M22 x BY cross. D. Number of non-BY eQTL increasing or decreasing *TDH3* promoter activity for each cross. E. eQTL shared among the three crosses irrespective of direction of effect. Areas are proportional to the number of eQTL shared. Brown: eQTL identified only in the M22 x BY cross. Blue: eQTL identified only in the YPS1000 x BY cross. Green: eQTL identified only in the SK1 x BY cross. Black: eQTL identified in all three crosses. F. Same as E, but for non-BY eQTL that increase *TDH3* promoter activity. G. Same as E, but for non-BY eQTL that decrease *TDH3* promoter activity.

Despite a difference of only 1% in *P_TDH3_-YFP* expression between M22 and BY, and of only 4% between YPS1000 and BY, we identified n=113 and n=101 statistically significant eQTL in these crosses, respectively. Between SK1 and BY, which showed a 47% difference in expression, we identified a similar number (n = 99) of statistically significant eQTL (Figure 3B). These numbers of eQTL identified in each experiment are approximately ten-fold greater than the number of eQTL identified for most genes in a prior eQTL mapping study (21) between BY and another strain of *S. cerevisiae,* RM11, which is closely related to M22 (Figure S2A).

To better understand why we found more eQTL than prior studies, we repeated our eQTL mapping experiments with fewer rounds of selection and recombination. We found that reducing the rounds of selection resulted in decreased statistical significance for many eQTL, but did not change the location or direction of effects for most eQTL inferred (red points in Figure 3C, Figure S3). By contrast, repeating these eQTL mapping experiments with crossing limited to a single round resulted in considerably fewer eQTL identified, regardless of the number of rounds of selection (black points in Figure 3C, Figure S3). These data suggest that although additional rounds of selection allowed eQTL with smaller effects to be identified, the high number of eQTL detected results primarily from increased recombination during multiple rounds of meiosis breaking apart physically close eQTL with opposite effects on expression. Indeed, we found that the non-BY alleles were evenly split between those that increased and decreased expression for all three mapping experiments (53 of 101 YPS1000 alleles increase expression, 51 of 99 SK1 alleles increase expression, and 55 of 113 M22 alleles increase expression, p > 0.6 for all, binomial test) (Figure 3D). These observations support the hypothesis that similar *trans*-regulatory effects on *TDH3* promoter activity are observed among strains of *S. cerevisiae* because compensatory alleles are maintained in the population.

To determine the similarity in genetic architecture of *trans*-regulatory variation affecting *TDH3* promoter activity among strains, we compared the genomic locations of eQTL identified in each pair of strains. If *trans*-regulatory variation is caused by the same loci in all strains, the ∼100 eQTLs identified in each comparison should map to similar genomic regions. However, if the sources of *trans*-regulatory variation affecting *TDH3* promoter activity segregating in *S. cerevisiae* are more diverse, eQTL identified in each comparison should map to different genomic regions. We found that the 313 eQTL identified mapped to 180 non-overlapping regions of the genome, with 27% (49 of 180) of these regions containing eQTL in only two of the comparisons and 22% (40 of 180) of these regions containing eQTL in all three comparisons (Figure 3E). Such shared eQTL regions might contain genes that contribute to variation in *trans*-regulation of *TDH3* promoter activity in multiple strains; however, this degree of overlap is not greater than expected by chance given the number and width of eQTL observed (p = 0.079, permutation test). Furthermore, in these shared genomic regions, only 18% of non-BY eQTL alleles had the same direction of effect on *TDH3* promoter activity in two comparisons (26 of 119 for increases, 18 of 120 for decreases), and only 6% of non-BY eQTL alleles had the same direction of effect in all three comparisons (7 of 119 for increases, 8 of 120 for decreases; p > 0.53, permutation test, Figure 3F-G). This lack of consistency in the direction of eQTL effects suggests that even if the same underlying loci contribute to *trans*-regulatory variation in multiple strains, the exact polymorphisms and their effects on *TDH3* promoter activity are likely to differ.

To further assess whether differences in eQTL inferred among strains were more likely to be due to biological differences than reproducibility among independent experiments, we repeated the mapping experiment between M22 and BY and compared the eQTL identified in the two replicate mapping experiments. Of the 74 eQTL found in the second M22/BY eQTL mapping experiment, 73% (54 eQTL) overlapped with eQTL from the initial M22/BY mapping experiment, which is significantly more than expected by chance (p < 0.001, permutation test, Figure S2C,D). This degree of overlap between the two M22/BY mapping experiments is greater than the degree of overlap between the second M22/BY experiment and both the YPS1000/BY (54%, 40 of 74 eQTL, p = 0.03, Fisher’s exact test) and SK1/BY (58%, 43 of 74 eQTL, p = 0.08, Fisher’s exact) test mapping experiments. In addition, the 113 eQTL identified in the initial mapping between M22 and BY strains included all 8 regions of the genome identified as affecting *TDH3* promoter activity in a cross between the BY and RM11 strains in a previous study, 7 of these eQTL alleles had effects in the same direction in both studies (28) (Figure S2E). This overlap is consistent with the close phylogenetic relationship between M22 and RM11 (Figure S2A). These results suggest that the differences in eQTLs mapped among the M22, YPS1000, SK1, and BY strains are unlikely to be primarily explained by low reproducibility of the mapping procedure, but rather reflect real differences in the genetic architecture of *trans*-regulatory variation affecting *TDH3* promoter activity among strains.

Taken together, our data suggest that there are hundreds of genetic variants segregating within *S. cerevisiae* that impact *TDH3* promoter activity in *trans*. These variants (i) differ among strains, (ii) cause increases and decreases in *TDH3* promoter activity with similar frequencies, and (iii) are located in hundreds of distinct regions in the genome. Although it might seem counterintuitive to find such extensive genetic variation affecting a trait whose variance has been limited by stabilizing selection, theoretical work has previously shown that stabilizing selection acting on quantitative traits can maintain abundant cryptic genetic variation with off-setting effects (46–50). These observations are consistent with the recently described “omnigenic model” of complex traits, which predicts that many quantitative trait loci with small, often opposing effects, are located throughout the genome and segregate within a population (62). The pervasiveness of genetic variants with opposing effects on expression might also explain the recurrent observation of compensatory evolution in genomic comparisons of gene expression within and between species (7, 9, 63–66). In addition, this variation might also form the basis for developmental systems drift in which phenotypes stay stable over evolutionary time, but the molecular components responsible for the phenotype change (67–70): with many combinations of alleles available that can produce the same trait value, changes in the regulation of a trait that do not alter the trait value might be common. Additional genetic mapping experiments that have similar power to resolve closely linked loci and detect alleles with very small effects are needed to determine whether the complex genetic architecture observed for the *trans*-regulation of the *TDH3* gene in *S. cerevisiae* is common to other genes, traits, and organisms.

## Methods

### Yeast strains and growth conditions

Strains used in this work are listed in Table S1. To determine variability in *TDH3* expression segregating within *S. cerevisiae*, we used haploid, MATalpha versions of 85 natural *S. cerevisiae* strains created in previous work (51). For each strain, we inserted a P*_TDH3_*-YFP reporter at the *HO* locus using a standard lithium acetate transformation approach with minor alterations (51, 71). The inserted reporter contained a copy of the *TDH3* promoter from the BY strain, a Yellow Fluorescent Protein (YFP) coding sequence, a CYC1 terminator, and a NatMX4 drug resistance marker. For 60 strains, we obtained successful integration and correct sequence of the reporter. Unless noted, all yeast growth was performed at 30°C in YPD (1% Difco yeast extract, 2% peptone, 2% glucose).

### Measurement of YFP expression

The *trans*-regulatory effects on *TDH3* promoter activity for each strain were estimated by measuring YFP expression from the P*_TDH3_*-YFP reporter. Strains were first revived from glycerol stocks on YPG (1% Difco yeast extract, 2% peptone, 2% glycerol) at 30°C. After 24 hours, each strain was inoculated into liquid YPD in a 96 well plate. For each plate, YFP positive (YPW1139) and YFP negative (YPW880) strains were included at specific locations in the 96 well plate as controls. This structure was replicated to solid YPD using a pin tool. To generate replicates, colonies were pin-tool replicated after 24 hours into twelve 96 well plates containing 500 μl of liquid YPD and grown for 24 hours. Cultures were then diluted 1/20 into fresh 500 μl of YPD and grown for an additional four hours. Samples were diluted 1/10 into 500 ul PBS and analyzed on an Accuri C6 flow cytometer connected to an Intellicyt autosampler.

Data was processed using the same procedure as described in Duveau et al. (2018) (45). Briefly, hard gates were used to remove flow cytometry artifacts and instances where multiple cells entered the flow cytometry detector at the same time based on estimates of cell size. For each sample, the most abundant monomorphic population was identified and the effect of cell size on fluorescence removed. For each event in a sample, the YFP levels were converted to estimates of mRNA expression using the formula E(mRNA) = exp(−7.820027*E(YFP)), which was based on a direct comparison of YFP fluorescence and mRNA abundance first reported in Duveau et al. (2018) (45). From these estimates, the population median was calculated. Using the control strains, linear models were used to remove batch effects such as differences among plates and variation due to the position of a sample (row and column) within a plate. Twelve replicate samples from each strain were combined to estimate strain averages. Four strains (NCYC110 (PJW1041), EM93 (PJW1055), YIIc17_E5 (PJW1038), and DBVPG3591 (PJW1053)) were excluded from analysis due to inconsistent measurements between replicates caused by flocculation and cell settling. All scripts used in data processing are included in the supporting files. Raw data is available for download from Flow Repository (FR-FCM-ZYVQ).

The effects of naturally occurring *cis*-regulatory variants on *TDH3* promoter activity within *S. cerevisiae* used data from Metzger et al. 2015 (53) (Flow Repository FR-FCM-ZZBN). The effects of new mutations on *TDH3* promoter activity used data from Metzger et al. 2016 (38) (Flow Repository FR-FCM-ZZNR). The original flow cytometry data from these previous studies was reprocessed with the same procedure as used in the current work.

### Testing for evidence and impacts of selection

To test for the action of natural selection on *trans*-acting factors affecting *TDH3* promoter activity, we developed an empirically based neutral model of gene expression evolution. To inform this model, we used previously collected data on the effects on *TDH3* promoter activity due to new mutations. Briefly, this prior work used ethyl methanesulfonate (EMS) to induce mutations in an isogenic yeast population containing the P*_TDH3_*-YFP reporter and used FACS to isolate ∼1500 individual genotypes irrespective of their YFP expression. Each isolated mutant contained ∼32 mutations relative to the reference strain, the overwhelming majority of which are expected to be *trans*-acting with respect to *TDH3* promoter activity (only two mutations in the *TDH3* promoter are expected among all individuals) (38). To estimate how *TDH3* promoter activity could evolve in the absence of natural selection, we generated a neutral distribution by sampling effects from this mutational distribution, combining the effects of mutations multiplicatively. We repeated this process 10,000 times to create a distribution of effects on *TDH3* promoter activity expected under neutrality for a given number of mutations.

To test for natural selection, we compared the effects of changes in *TDH3* promoter activity due to *trans*-regulatory differences among *S. cerevisiae* strains to our empirically derived neutral model. To account for the phylogenetic relationships among strains, we used a *S. cerevisiae* phylogeny estimated from whole genome polymorphism data(51). We used this phylogeny and the measured effects of each strains *trans*-regulatory background on *TDH3* promoter activity to estimate the ancestral state of *trans*-regulatory effects on *TDH3* promoter activity at each node in the phylogeny. Next, we used the *S. cerevisiae* phylogeny to estimate how many mutations had likely occurred along each branch. We then compared the changes in *trans*-regulatory effects along each branch to the corresponding distribution of effects derived from our neutral model. For each branch, we calculated the likelihood of the observed change in expression along that branch given the number of mutations that had occurred. We combined the likelihoods over all observed branches to determine the likelihood of the complete set of observed expression values and changes in expression on the phylogeny. Observed likelihoods less than expected under neutrality are consistent with positive selection for a new phenotypic value, whereas likelihoods greater than expected under neutrality are consistent with phenotypic constraint due to natural selection. Additional discussion of this approach can be found in SI methods.

To test the robustness of our inference to phylogenetic uncertainty, we repeated the analysis using genetic distance between strains instead of phylogenetic branch lengths to estimate the number of mutations that had occurred between strains. To avoid double counting of individual strains, we used each strain exactly once in the comparison. We then sampled which strains were compared 10,000 times to generate a distribution of observed effects.

As an alternative to tests for selection based on the empirical estimates of the effects of new regulatory mutations, we used the Brownian motion/Ornstein-Uhlenbeck framework to test for the presence of stabilizing selection on *trans*-acting factors affecting *TDH3* promoter activity. We followed the approach of Bedford and Hartl 2009. Briefly, two models of quantitative trait evolution were fit to the data. The Brownian motion model allows for trait values to diverge linearly with time, while the Ornstein-Uhlenbeck model includes an additional parameter that reflects the action of stabilizing selection. We tested whether the Ornstein-Uhlenbeck model fit significantly better than the Brownian motion model using a chi-square distribution with a single degree of freedom.

### eQTL Mapping

Genomic regions responsible for differences in *TDH3* promoter activity were identified by eQTL mapping. We crossed strains YPS1000 (PJW1057), SK1 (PJW1016), and M22 (PJW1072) that were MATalpha, nourseothricin resistant, and contained the P*_TDH3_*-YFP reporter to a version of BY (PJW1240, Figure S4) that was MATa, G418 resistant, and contained the P*_TDH3_*-YFP reporter. This common BY mapping strain also contained a Red Fluorescent Protein (RFP) marker at its mating type locus (72). Detailed methods for the creation of the common mapping strain can be found in SI methods. For each cross, we selected diploids using a combination of nourseothricin and G418 resistance and choose a single colony to ensure homogeneity in the genetic background. Each strain was then sporulated. To increase the amount of recombination in each cross, the resulting spores were mated and sporulated two additional times. YFP expression of the resulting spores was measured using flow cytometry and the 5% highest and lowest expressing cells collected. For each cross, sorted cells were allowed to reach saturation and then resorted based on YFP expression an additional two times. From each population, DNA was extracted and Illumina libraries created. Sequencing was performed on a HiSeq 2000 using 125 bp paired end sequencing at the University of Michigan Sequencing Core. Sequencing barcodes are listed in Table S2. Detailed methods on the mapping procedure can be found in SI methods.

### QTL Identification

After sequencing, samples were processed to identify individual eQTL. First, Sickle was used to remove low quality bases from each read using default setting (73). Next, Cutadapt was used to remove any adapter sequence from read ends (flags -e 0.2 -O 3 -m 15) (74). Samples were aligned to the S228c reference genome using bowtie2 (flags -I 0 - X 1000 --very-sensitive-local) (75) and then sorted and indexed using samtools (76). Overlapping reads were clipped using clipOverlap in bamUtil. SNPs were jointly called within each paired set of samples selected for high and low YFP fluorescence using freebayes (77). Identified SNPs were required to reach at least 20% frequency in at least one of the two paired samples and be observed at least 4 times across both samples.

For each pair of samples, SNPs were filtered based on quality and depth. Each SNP was required to have depth of at least 20 to ensure adequate power, a depth below 500 to reduce the number of SNPs called in low complexity sequences, a mapping quality score of greater than 30, and imbalance scores for left/right, center/end, and forward/reverse for SNP position within reads of less than 30. At each position, only the two highest likelihood SNPs were retained. For each SNP, we calculated a G statistic using a likelihood ratio test of alternative and reference alleles within the high and low selected populations(78). For SNPs where the alternative allele had a higher frequency than the reference allele in the high selected population relative to the low selected population, we maintained the sign of G. For SNPs where the alternative allele had a lower frequency than the reference allele in the high selected population relative to the low selected population, we flipped the sign of G. We then calculated G’ by averaging these estimates over a 40 kb window centered on the SNP (78). Finally, to identify QTL peaks, we located all local maxima and minima in G’. We called significant peaks those with G’ > 5 or G’ < 5. The location of each peak was defined as the distance needed for G’ to drop by 5 from the peak. Local peaks whose locations overlapped were merged into a single peak.

## Acknowledgements

We would like to thank Fabien Duveau, Andrea Hodgins-Davis, Mark Hill, Mo Siddiq, Petra Vande Zande, and members of the Wittkopp lab for helpful comments on the manuscript. Support and access to flow cytometry equipment was provided by the University of Michigan Center for Chemical Genomics and Flow Cytometry Core. This work was supported by the University of Michigan Rackham Graduate School and National Institutes of Health Genome Sciences training grant (T32 HG000040) to B.P.H.M, and the National Science Foundation (MCB-1021398) and National Institutes of Health (1R35GM118073 and 1 R01 GM108826) to P.J.W.

## References

1. Wittkopp PJ, Haerum BK, Clark AG (2004) Evolutionary changes in *cis* and *trans* gene regulation. Nature 430(6995):85–88.

2. Wang D, et al. (2007) Expression evolution in yeast genes of single-input modules is mainly due to changes in *trans*-acting factors. Genome Res 17(8):1161–9.

3. Sung H-M, et al. (2009) Roles of *Trans* and *Cis* Variation in Yeast Intraspecies Evolution of Gene Expression. Mol Biol Evol 26(11):2533–2538.

4. Zhang X, Borevitz JO (2009) Global analysis of allele-specific expression in *Arabidopsis thaliana*. Genetics 182(4):943–54.

5. Emerson JJ, et al. (2010) Natural selection on *cis* and *trans* regulation in yeasts. Genome Res 20:826–836.

6. Bell GDM, Kane NC, Rieseberg LH, Adams KL (2013) RNA-seq analysis of allele-specific expression, hybrid effects, and regulatory divergence in hybrids compared with their parents from natural populations. Genome Biol Evol 5(7):1309–1323.

7. Schaefke B, et al. (2013) Inheritance of Gene Expression Level and Selective Constraints on *Trans*- and *Cis*-Regulatory Changes in Yeast. Mol Biol Evol 30(9):2121–33.

8. Suvorov A, et al. (2013) Intra-specific regulatory variation in *Drosophila pseudoobscura*. PLoS One 8(12):e83547.

9. Coolon J, McManus CJ, Stevenson KR, Graveley BR, Wittkopp PJ (2014) Tempo and mode of regulatory evolution in *Drosophila*. Genome Res 24:797–808.

10. Chen J, Nolte V, Schlötterer C (2015) Temperature Stress Mediates Decanalization and Dominance of Gene Expression in *Drosophila melanogaster*. PLoS Genet 11(2):e1004883.

11. Gilad Y, Rifkin S a., Pritchard JK (2008) Revealing the architecture of gene regulation: the promise of eQTL studies. Trends Genet 24(8):408–415.

12. Hansen BG, Halkier B a., Kliebenstein DJ (2008) Identifying the molecular basis of QTLs: eQTLs add a new dimension. Trends Plant Sci 13(February):72–77.

13. Majewski J, Pastinen T (2011) The study of eQTL variations by RNA-seq: from SNPs to phenotypes. Trends Genet 27(2):72–79.

14. Cubillos FA, Coustham V, Loudet O (2012) Lessons from eQTL mapping studies: Non-coding regions and their role behind natural phenotypic variation in plants. Curr Opin Plant Biol 15(2):192–198.

15. Nica AC, Dermitzakis ET (2013) Expression quantitative trait loci: present and future. Philos Trans R Soc B Biol Sci 368(1620):20120362–20120362.

16. Westra HJ, Franke L (2014) From genome to function by studying eQTLs. Biochim Biophys Acta - Mol Basis Dis 1842(10):1896–1902.

17. Albert FW, Kruglyak L (2015) The role of regulatory variation in complex traits and disease. Nat Rev Genet 16(4):197–212.

18. Pai A a., Pritchard JK, Gilad Y (2015) The Genetic and Mechanistic Basis for Variation in Gene Regulation. PLoS Genet 11(1):e1004857.

19. Yvert G, et al. (2003) *Trans*-acting regulatory variation in *Saccharomyces cerevisiae* and the role of transcription factors. Nat Genet 35(1):57–64.

20. Schadt EE, et al. (2003) Genetics of gene expression surveyed in maize, mouse and man. Nature 205(October 2002):1–6.

21. Albert FW, Bloom JS, Siegel J, Day L, Kruglyak L (2018) Genetics of *trans*-regulatory variation in gene expression. Elife 7:1–39.

22. Ronald J, Brem RB, Whittle J, Kruglyak L (2005) Local regulatory variation in *Saccharomyces cerevisiae*. PLoS Genet 1(2):0213–0222.

23. Brem RB, Yvert G, Clinton R, Kruglyak L (2002) Genetic dissection of transcriptional regulation in budding yeast. Science 296(5568):752–5.

24. Brem RB, Storey JD, Whittle J, Kruglyak L (2005) Genetic interactions between polymorphisms that affect gene expression in yeast. Nature 436(7051):701–703.

25. Brem RB, Kruglyak L (2005) The landscape of genetic complexity across 5,700 gene expression traits in yeast. Proc Natl Acad Sci 102(5):1572–7.

26. Smith EN, Kruglyak L (2008) Gene-environment interaction in yeast gene expression. PLoS Biol 6(4):e83.

27. Parts L, et al. (2014) Heritability and genetic basis of protein level variation in an outbred population. Genome Res 24(8):1363–70.

28. Albert FW, Treusch S, Shockley AH, Bloom JS, Kruglyak L (2014) Genetics of single-cell protein abundance variation in large yeast populations. Nature 506(7489):494–497.

29. Salinas F, et al. (2016) Natural variation in non-coding regions underlying phenotypic diversity in budding yeast. Sci Rep 6:21849.

30. Kita R, Venkataram S, Zhou Y, Fraser HB (2017) High-resolution mapping of *cis*-regulatory variation in budding yeast. Proc Natl Acad Sci:201717421.

31. Gruber JD, Long AD (2009) *Cis*-regulatory variation is typically polyallelic in Drosophila. Genetics 181(2):661–670.

32. de Meaux J (2005) Allele-Specific Assay Reveals Functional Variation in the *Chalcone Synthase* Promoter of *Arabidopsis thaliana* That Is Compatible with Neutral Evolution. Plant Cell Online 17(3):676–690.

33. Moyerbrailean GA, et al. (2016) High-throughput allele-specific expression across 250 environmental conditions. Genome Res:gr.209759.116.

34. Kang EY, et al. (2016) Discovering SNPs Regulating Human Gene Expression Using Allele Specific Expression from RNA-Seq Data. Genetics 204(November):1057–1064.

35. Gilad Y, Oshlack A, Rifkin SA (2006) Natural selection on gene expression. Trends Genet 22(8):456–461.

36. Denver DR, et al. (2005) The transcriptional consequences of mutation and natural selection in *Caenorhabditis elegans*. Nat Genet 37(5):544–548.

37. Josephs EB, Lee YW, Stinchcombe JR, Wright SI (2015) Association mapping reveals the role of purifying selection in the maintenance of genomic variation in gene expression. Proc Natl Acad Sci 112(50):15390–15395.

38. Metzger BPH, et al. (2016) Contrasting Frequencies and Effects of *cis*- and *trans*-Regulatory Mutations Affecting Gene Expression. Mol Biol Evol 33(5):1131–1146.

39. Lemos B, Araripe LO, Fontanillas P, Hartl DL (2008) Dominance and the evolutionary accumulation of *cis*- and *trans*-effects on gene expression. Proc Natl Acad Sci 105(38):14471–6.

40. Gruber JD, Vogel K, Kalay G, Wittkopp PJ (2012) Contrasting Properties of Gene-specific Regulatory, Coding, and Copy Number Mutations in *Saccharomyces cerevisiae*: Frequency, Effects and Dominance. PLoS Genet 8(2):e1002497.

41. Landry CR, Lemos B, Rifkin SA, Dickinson WJ, Hartl DL (2007) Genetic properties influencing the evolvability of gene expression. Science 317(5834):118–121.

42. Fay JC, Wittkopp PJ (2008) Evaluating the role of natural selection in the evolution of gene regulation. Heredity (Edinb) 100(2):191–9.

43. Stern DL (2000) Perspective: Evolutionary Developmental Biology and the Problem of Variation. Evolution 54(4):1079.

44. Duveau F, Toubiana W, Wittkopp PJ (2017) Fitness Effects of *Cis*-Regulatory Variants in the *Saccharomyces cerevisiae* TDH3 Promoter. Mol Biol Evol 8(3):206–216.

45. Duveau F, et al. (2018) Fitness effects of altering gene expression noise in *Saccharomyces cerevisiae*. Elife 7(e37272). doi:10.1101/294603.

46. Lande R (1976) The maintenance of genetic variability by mutation in a polygenic character with linked loci. Genet Res (Camb) 89(5–6):373–387.

47. Turelli M (1984) Heritable genetic variation via mutation-selection balance: Lerch’s zeta meets the abdominal bristle. Theor Popul Biol 25:138–193.

48. Barton NH (1986) The maintenance of polygenic variation through a balance between mutation and stabilizing selection. Genet Res 47(3):209–216.

49. Barton N (1989) The divergence of a polygenic system subject to stabilizing selection, mutation and drift. Genet Res 54(1):59–78.

50. Dover GA, Flavell RB (1984) Molecular coevolution: DNA divergence and the maintenance of function. Cell 38(3):622–623.

51. MacLean CJ, et al. (2017) Deciphering the Genic Basis of Yeast Fitness Variation by Simultaneous Forward and Reverse Genetics. Mol Biol Evol 34(10):1–17.

52. Peter J, et al. (2018) Genome evolution across 1011 Saccharomyces cerevisiae isolates. doi:10.1038/s41586-018-0030-5.

53. Metzger BPH, Yuan DC, Gruber JD, Duveau FD, Wittkopp PJ (2015) Selection on noise constrains variation in a eukaryotic promoter. Nature 521(May 21):344–347.

54. Bedford T, Hartl DL (2009) Optimization of gene expression by natural selection. Proc Natl Acad Sci U S A 106(4):1133–8.

55. Schlötterer C, Tobler R, Kofler R, Nolte V (2014) Sequencing pools of individuals — mining genome-wide polymorphism data without big funding. Nat Rev Genet 15(11):749–763.

56. Parts L, et al. (2011) Revealing the genetic structure of a trait by sequencing a population under selection. Genome Res 21(7):1131–8.

57. Kofler R, Pandey RV, Schlötterer C (2011) PoPoolation2: identifying differentiation between populations using sequencing of pooled DNA samples (Pool-Seq). Bioinformatics 27(24):3435–6.

58. Ehrenreich IM, et al. (2010) Dissection of genetically complex traits with extremely large pools of yeast segregants. Nature 464(7291):1039–42.

59. Ehrenreich IM, et al. (2012) Genetic Architecture of Highly Complex Chemical Resistance Traits across Four Yeast Strains. PLoS Genet 8(3):e1002570.

60. Cubillos FA, et al. (2013) High-resolution mapping of complex traits with a four-parent advanced intercross yeast population. Genetics 195(3):1141–55.

61. Duveau F, et al. (2014) Mapping Small Effect Mutations in *Saccharomyces cerevisiae*: Impacts of Experimental Design and Mutational Properties. G3 Genes|Genomes|Genetics 4(July):1205–1216.

62. Boyle EA, Li YI, Pritchard JK (2017) An Expanded View of Complex Traits: From Polygenic to Omnigenic. Cell 169(7):1177–1186.

63. Mack KL, Campbell P, Nachman MW (2016) Gene regulation and speciation in house mice. Genome Res 26:451–461.

64. Goncalves A, et al. (2012) Extensive compensatory cis-trans regulation in the evolution of mouse gene expression. Genome Res 22(12):2376–2384.

65. Metzger BPH, Wittkopp PJ, Coolon JD (2017) Evolutionary dynamics of regulatory changes underlying gene expression divergence among Saccharomyces species. Genome Biol Evol 177(4):1987–1996.

66. Verta JP, Landry CR, Mackay J (2016) Dissection of expression-quantitative trait locus and allele specificity using a haploid/diploid plant system - insights into compensatory evolution of transcriptional regulation within populations. New Phytol 211(1):159–171.

67. Pavlicev M, Wagner GP (2012) A model of developmental evolution: selection, pleiotropy and compensation. Trends Ecol Evol:1–7.

68. Brachi B, et al. (2010) Linkage and Association Mapping of *Arabidopsis thaliana* Flowering Time in Nature. PLoS Genet 6(5):e1000940.

69. True JR, Haag ES (2001) Developmental system drift and flexibility in evolutionary trajectories. Evol Dev 3(2):109–119.

70. Gordon KL, Ruvinsky I (2012) Tempo and Mode in Evolution of Transcriptional Regulation. PLoS Genet 8(1):e1002432.

71. Cubillos FA, Louis EJ, Liti G (2009) Generation of a large set of genetically tractable haploid and diploid *Saccharomyces* strains. FEMS Yeast Res 9(8):1217–25.

72. Chin BL, Frizzell M a., Timberlake WE, Fink GR (2012) FASTER MT: Isolation of Pure Populations of a and α Ascospores from *Saccharomyces cerevisiae*. G3 Genes|Genomes|Genetics 2(4):449–452.

73. Joshi, N.A., Fass JN (2011) Sickle: A sliding-window, adaptive, quality-based trimming tool for FastQ files.

74. Martin M (2011) Cutadapt removes adapter sequences from high-throughput sequencing reads. EMBnet.journal 17(1):10–12.

75. Langmead B, Salzberg SL (2012) Fast gapped-read alignment with Bowtie 2. Nat Methods 9(4):357–359.

76. Li H, et al. (2009) The Sequence Alignment/Map format and SAMtools. Bioinformatics 25(16):2078–2079.

77. Garrison E, Marth G (2012) Haplotype-based variant detection from short-read sequencing. ArXiv. doi:10.1126/science.1241459.

78. Magwene PM, Willis JH, Kelly JK (2011) The statistics of bulk segregant analysis using next generation sequencing. PLoS Comput Biol 7(11):1–9.

