## Supplementary Information for "Complex polymorphic genetic architecture underlies *trans*-regulatory variation among strains of *Saccharomyces cerevisiae*"

**This PDF file includes:**

Supplementary Methods
Figs. S1 to S4
Tables S1 to S4

**Other supplementary materials for this manuscript include the following:**

Data S1

### Supplementary Information Text
### Methods.

*Test for selection using empirically derived distributions of mutational affects*
The empirically based models of gene expression evolution make several assumptions about the extent and role of epistasis in the evolution of quantitative traits. First, these projections assume that the effects of mutations are additive and do not depend on the effects of prior mutations. This assumption is reasonable if there are many loci in which effects on expression can occur and thus multiple, independent, routes by which *TDH3* promoter activity can be altered. Given the mapping results, this assumption seems reasonable. Second, the neutral projections assume that the mutational distribution itself is stable over time, i.e. that having observed a specific set of changes in expression doesn't alter the set of changes in expression that are possible in the future. This assumption is reasonable for changes in expression that are common, but its validity for effects that are rarely produced by mutation is less clear. For example, if the set of mutations with the largest increases in expression are all due to mutations in the same gene or pathway, then only the first mutation in this pathway is likely to have a large effect. Unfortunately, the stability or fluidity of the distribution of mutational effects is currently unknown. In addition, the stability of the mutational distribution over time requires that mutations have non-additive effects. Thus, while the neutral projections have assumed both additivity of effects and stability of the mutational distribution over time, both assumptions cannot be strictly true. Additional work is needed to know the extent to which not only the effects of *trans*-regulatory mutations are altered by the occurrence of other *trans*-regulatory mutations, but also the extent to which individual mutations alter the entire distribution of mutational effects available to future evolution.

In addition to incorporating the effects of new mutations in the test for selection, this approach explicitly considers the phylogenetic relationship among strains. Standard approaches for identifying the action of selection on quantitative traits assume that the underlying distribution of mutation effects is normal (Gaussian). This assumption implies time irreversibility and allows for explicit calculation of the effects on expression on an unrooted phylogeny without identifying the direction of expression changes. However, empirically derived mutational effects are unlikely to be normally distributed. As a consequence, explicit calculations of likelihoods are not possible with empirical distributions and current approaches. As a consequence, the approach we used makes assumptions about the ancestral states of *TDH3 trans*-regulation to allow for estimating likelihoods by simulation. In particular, it assumes that the expression value for each ancestral node is the average of the descendent node values weighted by the branch lengths. This assumption requires that all ancestral values are intermediate of decedent values, an assumption that is unlikely to be true for a quantitative trait that varies within a population. In addition, it is currently unclear how to incorporate uncertainty in the effects of new mutations within an explicitly phylogenetic framework, nor how to combine empirical estimates of mutational effects with the reality that phylogenetic relationships vary across the genome. While we obtain the same results for the action of selection on *TDH3 trans*-regulation when we individually relax these assumptions, the approaches and methods needed to combine empirical observations about quantitative traits in an explicitly phylogenetic context are currently lacking.

*eQTL mapping approach with multiple rounds of sporulation and selection*
To map eQTL affecting *TDH3* expression, we used a procedure that included multiple rounds of sporulation and selection. To begin, hybrids for each cross (Figure S2B, P.0) were sporulated. To do so, hybrids were grown on GNA (1% Difco yeast extract, 3% Difco nutrient broth, 5% glucose) media for 12-16 hours, transferred to KAc plates, and maintained at room temperature until at least 50% of the cells had sporulated. Cells were then washed twice in 1 ml of water and incubated with 200 ul of 0.3mg/ml 100T zymolyase for one hour with agitation. Next, cells were washed with 1 ml of water and resuspended in 100 ul of water. Cells were vortexed for 2 minutes to stick spores to the tube wall. The supernatant was removed and 1 ml of water was added. Without agitation, this 1 ml was removed and a second 1 ml of $H_2O$ was added. This 1 ml was also removed and 1ml of triton-X (0.02%) was added. Samples were sonicated on ice for 10 seconds at medium power (3.5 on a Sonic Dismembrator Model 100, Fisher). Spores were confirmed to be separated and diploids absent by visual inspection under a microscope.

After spore isolation, the population was split into thirds. One third was added to 1 ml YPD, grown to saturation overnight, and then frozen at -80°C as a glycerol stock. The second third was sorted for the absence of the RFP marker using fluorescence assisted cell sorting (FACS) on a FACS canto II at the University of Michigan Flow Cytometry Core (Figure S2B, P.1). Because all MAT**a** and diploid strains express RFP, this sorting captures only MATalpha cells. For each cross, we collected > $10^6$ individuals lacking RFP fluorescence (Figure S2B, F.1). These were incubated with 1ml YPD and grown for 24-28 hours. The final third was used to initiate additional rounds of crossing by plating onto YPD. After growth overnight, sporulation and spore isolation was repeated. Isolated spores from the second round of sporulation were plated onto YPD and sporulated a final time. After this third round of sporulation, cells lacking RFP fluorescence were again collected (Figure S2B, F.3).

To identify the genetic basis of differences in YFP expression between strains, cells were transferred to 1 ml PBS and cells with the 5% highest (Figure S2B, H.1.1 and H.3.1) or the 5% lowest (Figure S2B, L.1.1 and L.3.1) YFP expression corrected for cell size were sorted from cells within the middle 80% of the cell size distribution based on forward scatter (FSC-H). For each sample, 100,000 individuals were collected from each tail. Each sorted population was grown in liquid YPD for 20 hours after which one half was frozen. To further enrich genotypes with high and low YFP expression within the sorted populations, the second half of each sample was used to initiate two additional rounds of sorting. For each round, the same sorting procedure as above was followed with one exception; populations originally sorted for high YFP expression were only sorted for high YFP expression and populations sorted for low YFP expression were only sorted for low YFP expression.

After all selection steps were completed, samples were revived from glycerol stocks and grown in 1 ml of YPD for 2 hours. DNA was extracted from each sample using the Purgene Yeast Kit from Qiagen. DNA concentration was determined using a Qubit, and Illumina Nextera XT libraries were prepared following the manufacturers guidelines. Barcodes for each sample are listed in Table S2. Library quality was assessed using the bioanalzyer and all samples were pooled equally using concentration estimates from the Qubit.

*Creation of mapping strain*
Determining the genetic and molecular mechanisms underlying complex phenotypes often requires identifying the causative genetic loci and nucleotides contributing to these traits (1). However, in the yeast *Saccharomyces cerevisiae*, the primary laboratory strains, S288c and its descendants, have several phenotypes that limit their usefulness in high throughput mapping approaches.

For example, *S. cerevisiae* isolates from the wild readily undergo meiosis under nutrient starvation and the majority of individual diploids sporulate. By contrast, S288c enters meiosis slowly and only a small proportion of individuals successfully complete meiosis, even under ideal conditions (2, 3). Because genetic mapping requires recombination, and thus, meiosis, the limited meiotic abilities of S288c reduces the number and speed at which mapping populations can be created.

In addition to poor sporulation, S288c and its descendants generate petite cells lacking mitochondria with high frequency. As a consequence, these individuals cannot perform aerobic respiration and often have altered phenotypes compared to wild-type individuals (4). Because linking phenotypes to their genomic location requires high quality phenotyping, additional variation introduced by petite individuals can reduce the accuracy and power of genetic mapping.

Lastly, upon meiosis yeast generate both **a** and $\alpha$ haploids. These haploids will readily reform diploids if not prevented, thus introducing additional variation due to ploidy into a mapping population. Current techniques for limiting the recreation of diploids suffer from a lack of throughput and poor specificity (5). As a consequence, the power to map the genetic basis of recessive traits in yeast is reduced.

To overcome these deficiencies, we modified S288c to increase its sporulation rate and density, reduce the frequency at which it generated petites, and to express a fluorescent marker that allowed easy identification of mating type. To accomplish these goal, we obtained several strains derived from S288c. These strains vary in their mating type and auxotrophies, facilitating crossing. In addition, these strains differ at a set of alleles derived from natural *S. cerevisiae* strains that either improve sporulation rate or lower petite frequency. These include versions of *TAO3* and *RME1* that increase sporulation rate (3) and versions of *SAL1*, *CAT5*, and *MIP1* that decrease petite frequency (6). An allelic variant at *MKT1* has also been identified that affects both sporulation and petite frequency. However, while the wild-type S288c MKT1 allele decreases sporulation rate, it also substantially reduces petite frequency compared to the alternative allele and we kept the S288c version (3, 6).

Through a series of crosses, transformations, and sporulations, we isolated a single individual that contained the desired set of alleles and was free of all auxotrophies except for ura3□0 (**Error! Reference source not found.**). We retained the *URA3* auxotrophy to facilitate future genetic manipulation by the *delitto perfetto* method, which requires *5-FOA* counter-selection and therefore a starting strain that is ura- (7). To facilitate the creation of the correct strain, we tracked the allelic identity of each segregating locus using pyrosequencing (Table S3, Table S4). After identification, the isolated individual was turned into a diploid and sporulated to generate isogenic **a** and $\alpha$ haploids. To the **a** haploid, we introduced a red fluorescent protein into the *MAT* locus (8). This marker allows identification of individuals based on their mating type using fluorescence assisted cell sorting (FACS). Using this marker, populations containing millions of individuals of the same mating type can be collected in minutes. Finally, both **a** and $\alpha$ strains contain a *TDH3* promoter driving Yellow Fluorescent Protein expression located at the *HO* gene to facilitate mapping of mutations and polymorphisms influencing *TDH3* expression.

Growth was performed using YPD. For crosses involving auxotrophies, synthetic complete media was used, minus the appropriate amino acids (1.7g Yeast nitrogen base, 5 g Ammonium Sulfate, 20g glucose per 1 L water; 20g agar for solid plates). Sporulation was induced by growth on YPD plates for 24 hours at room temperature, followed by plating on KAc plates at room temperature (10g Potassium acetate, 0.5g glucose per 1 L water; 20g agar for solid plates). Ascus walls were dissolved prior to tetrad dissections by incubating spores in 200 µl zymolyase (1 mg/ml 20T) for 1 hour without shaking.

To create homozygous diploids from haploids, strains were transformed with plasmid pCM66. pCM66 contains a galactose inducible copy of *HO* and a selective nourseothricin resistance marker. After transformation, nourseothricin resistant cells were grown with galactose as the sole carbon source at 30°C without shaking for 8 hours to induce expression of *HO*. This allowed for mating type switching and subsequent mother-daughter cell mating to produce diploids. Cells were streaked for single colonies on YPD plates and the ploidy of single colonies checked by colony PCR using mating-type specific primers. Diploid colonies were streaked onto fresh, non-selective, YPD plates and assayed for loss of nourseothricin resistance, and thus pCM66.

Pyrosequencing was used to follow sporulation and petite QTN. Methods are as described in (9). PCR primers used are Table S3. Dispensation order for pyrosequencing is in Table S4.

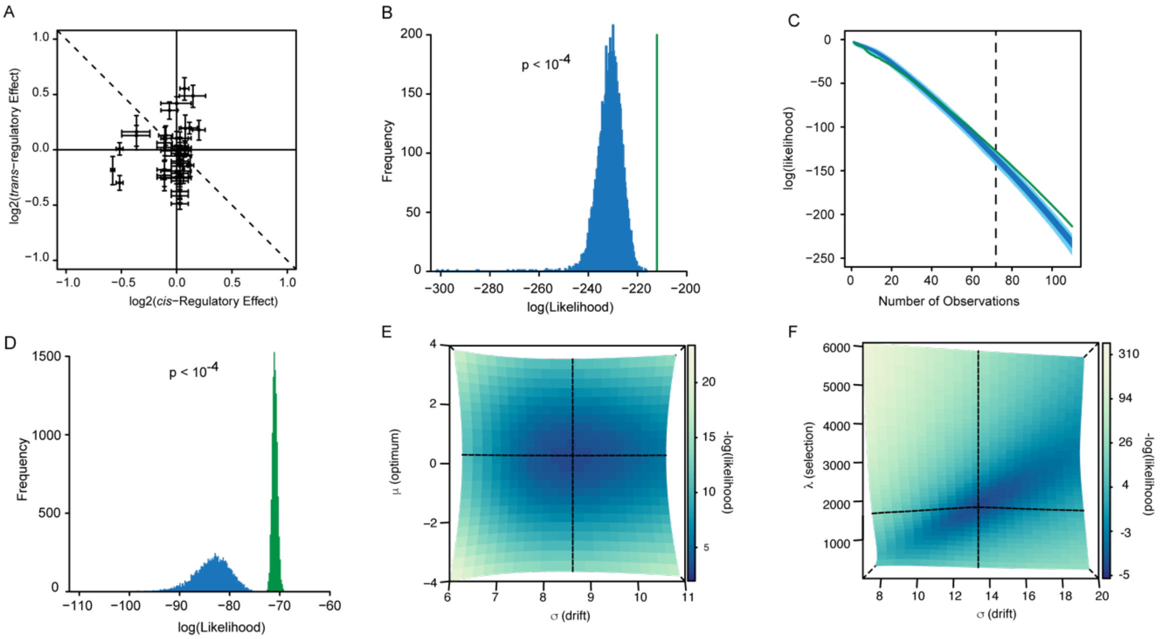


| BM | ML Estimate | 95% CI |
|---|---|---|
| μ | 0.316 | -1.48 to 2.11 |
| σ | 8.62 | 6.93 to 10.3 |

| OU | ML Estimate | 95% CI |
|---|---|---|
| μ | 0.038 | -0.06 to 0.13 |
| σ | 13.4 | 8.0 to 18.8 |
| λ | 1631.8 | 163.9 to 3100 |

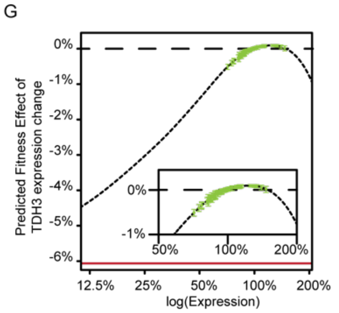


Fig. S1. A. Estimated effects of regulatory variation among natural yeast strains. X-axis, *cis*-regulatory variation due to differences within the *TDH3* promoter. Estimated from prior data that incorporated different *TDH3* promoter sequences into a common reference strain expressing YFP. Y-axis, *trans*-regulatory variation due to differences outside of the *TDH3* promoter. Estimated by inserting a common *TDH3* promoter into different genomic backgrounds. Each point is for a different strain. Error bars are 95% confidence intervals on the mean. Dashed line indicates no net change in TDH3 expression due to compensatory changes in *cis*- and *trans*-regulation. B. Statistical test of natural selection on *trans*-acting variants affecting *TDH3* promoter activity. Blue, distribution of likelihoods expected under neutral model. Green, observed likelihood of *TDH3* expression given the phylogenetic relationships among strains. C. Change in likelihood as additional observations are added. Blue, likelihoods under neutral model. Dark blue gives the middle 50% of the probability space. Light blue gives is the 90% quantiles. Green is observed likelihood. Dashed line is the number of observations needed before the observed data becomes significantly higher than expected under neutrality ($p < 0.01$). D. Statistical test of natural selection on *trans*-acting variants affecting *TDH3* promoter activity. Blue, distribution of likelihoods expected under neutral model. Green, observed likelihood of *TDH3* expression given the genetic distances among strains. E. Likelihood surface for Brownian motion model of trait evolution. Dashed lines give maximum likelihood estimate of parameters. Maximum likelihood estimates and 95% confidence intervals for each parameter of the model are listed below the figure. F. Same as E, but for Ornstein-Uhlenbeck model. G. Estimated fitness effect of each strains *trans*-regulatory background on *TDH3* expression. Small dashed line gives relationship between expression and fitness. Large dashed line is no fitness change. Red line, fitness cost of TDH3 deletion. Green, individuals strains estimated fitness effect. Error bars are 95% confidence intervals on the mean. Inset shows enlarged view of strains near WT expression levels.

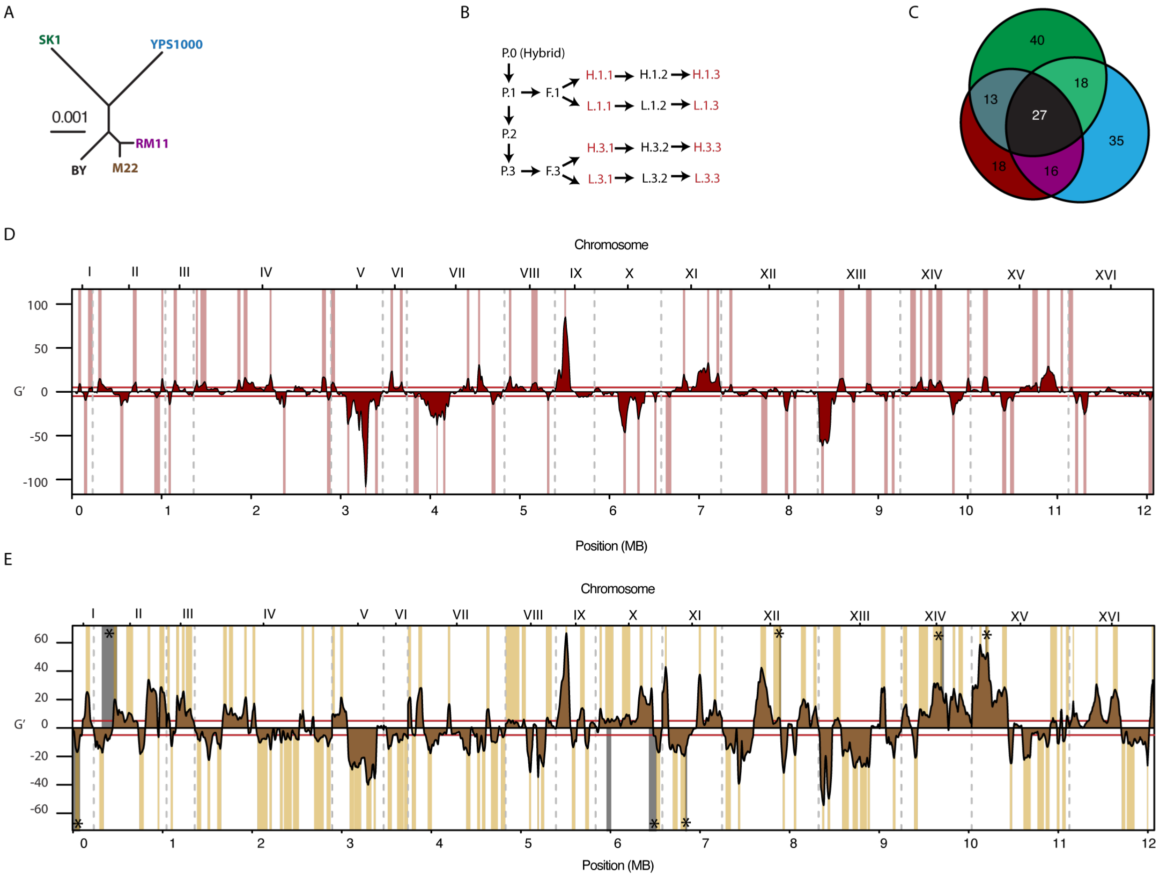


Fig. S2. A. Phylogenetic relationships among strains used in eQTL mapping. Prior work has used the RM11 strain, which is most closely related to the M22 strain used here. B. Mapping procedure. Initial hybrids (P.0) were sporulated to produce population P.1. To increase mapping resolution, spores in P.1 were mated and sporulated two additional times (P.2 and P.3). For the P.1 and P.3 populations, a single mating type was isolated by FACS using a red fluorescent protein incorporated at the mating type locus (F.1. and F.3). Populations were then subjected to FACS based on YFP expression, with the top (H) and bottom (L) 5% of cells sorted in each case (H.1.1, L.1.1, H.3.1, L.3.1). To increase frequency differences between these high and low pools, two additional rounds of FACS based selection were applied. Populations highlighted in red were sequenced. C. Overlap of eQTL between the second M22 x BY eQTL mapping experiment and the YPS1000 x BY and SK1 x BY mapping experiments.  Red: eQTL identified only in the second M22 x BY cross. Blue: eQTL identified only in the YPS1000 x BY cross. Green: eQTL identified only in the SK1 x BY cross. Black: eQTL identified in all three crosses. D. G' statistic for evidence of eQTL in the second M22 x BY eQTL mapping expeirment. Effects are relative to the non-BY reference allele. Dashed gray lines indicate chromosome boundaries. Dashed red lines gives threshold for statistical significance. Called eQTL with 95% confidence intervals on the location are highlighted. E. Same as D, but for the first M22 x BY eQTL mapping experiment. In addition, black regions highlight eQTL identified in previous work using strains RM11 and BY. Asterisk indicate eQTL in the same direction from the two studies.

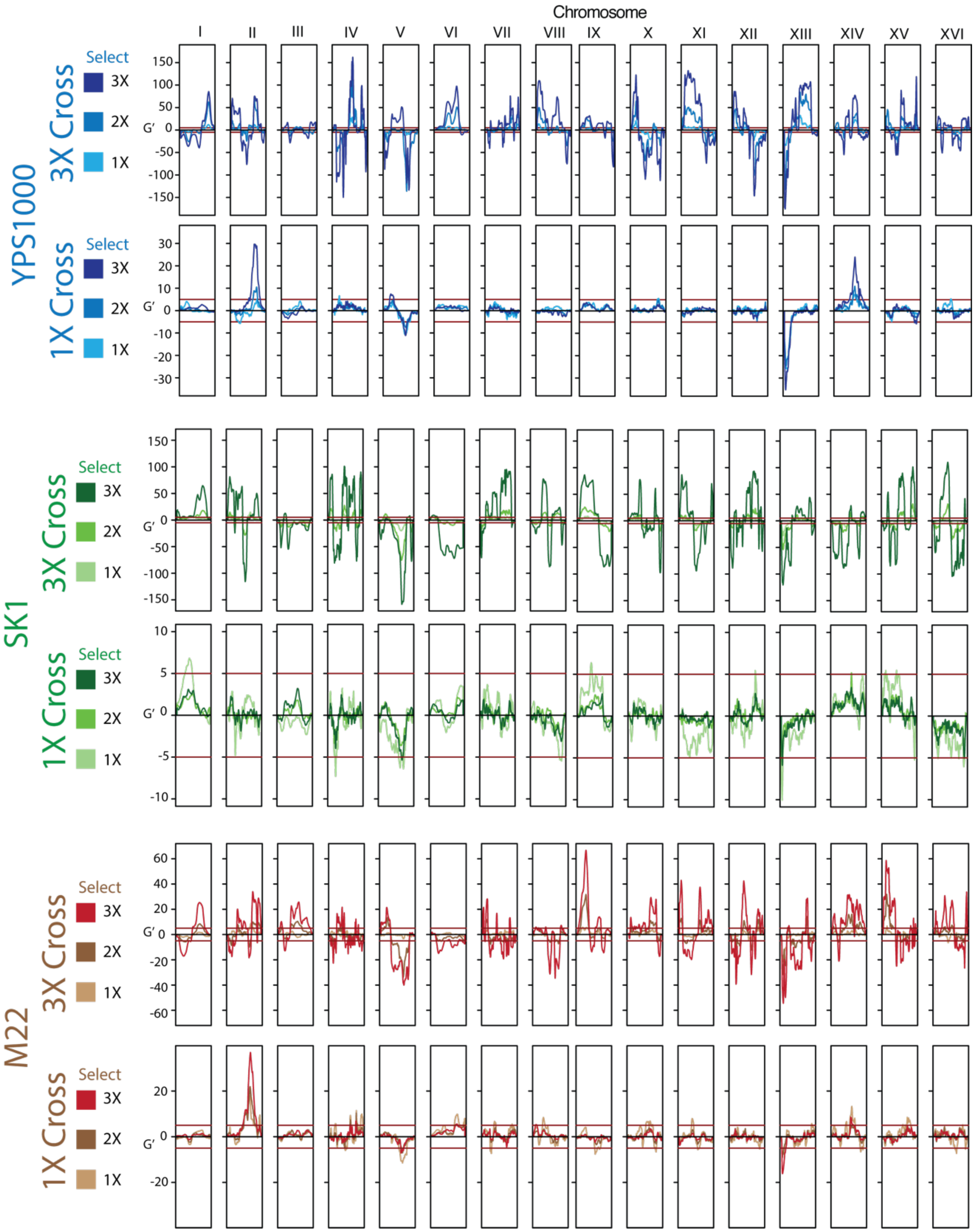


Fig. S3. eQTL mapping data for all three crosses under multiple mapping procedures. Brown, M22 x BY. Blue, YPS1000 x BY. Green, SK1 x BY. For each cross, three rounds of crossing are on top, one round of crossing on the bottom. Darker colors are for more rounds of selection. Red lines give thresholds for statistical significance.

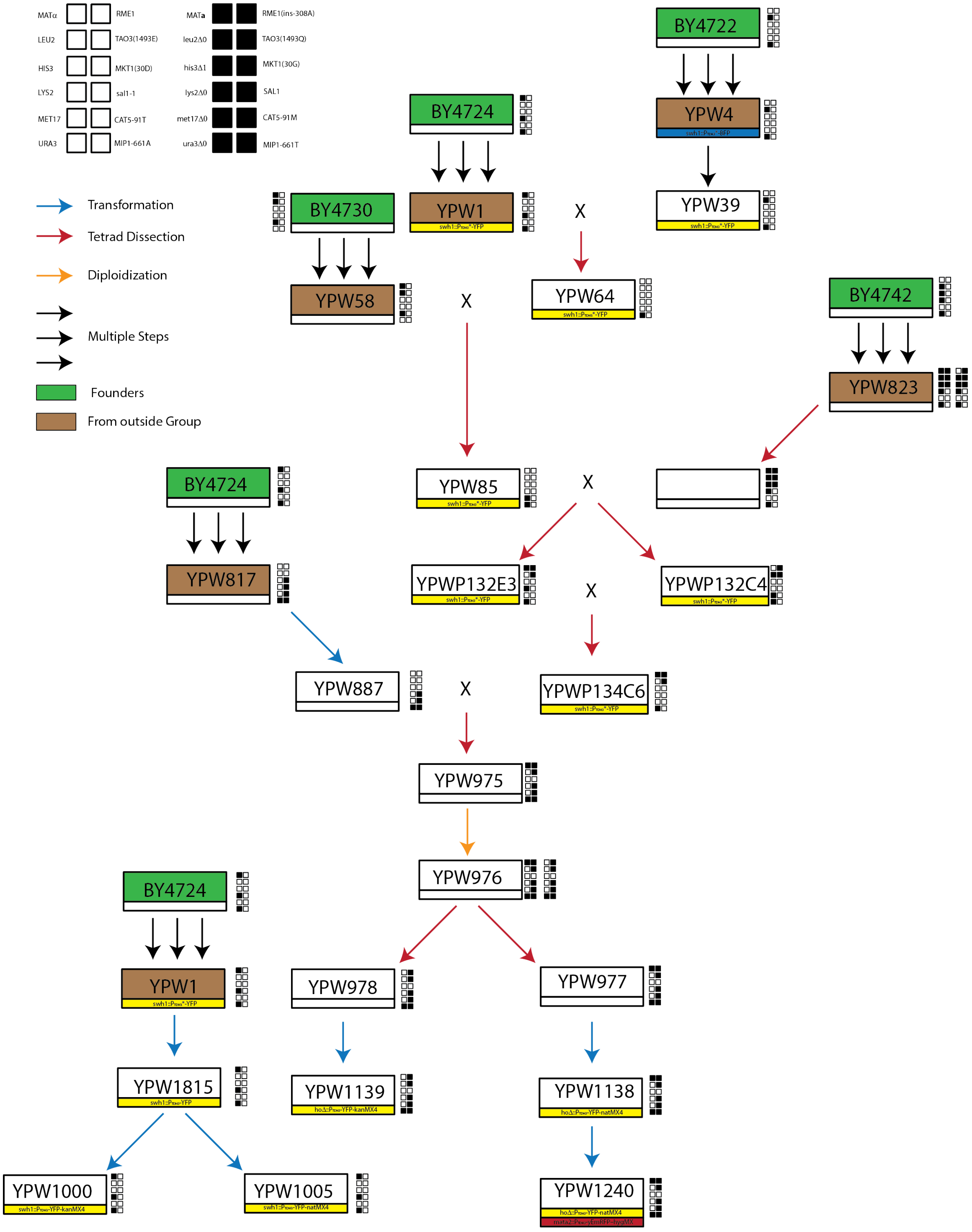


Fig. S4. Creation of common mapping strain YPW1240. Each box represents a unique strain. Strains in green are the original founding strains and strains in brown are those originally obtained by the lab. Genotypes at segregating markers are shown to the right of each strain. Auxotrophies and mating type on the left, sporulation and petite alleles on the right. The shading of the box indicates the allele at a particular locus for each strain as given in the upper left corner.

All crosses are designated by an X, tetrad dissections by a red arrow, transformations by a blue arrow, and diploidizations by an orange arrow. Black arrows represent multiple steps taken prior to receiving a strain. Below each strain, the identity of fluorescent markers is labeled.

**Table S1 Natural strains used for *trans*-regulatory polymorphism effects**
Strain names and collection number for natural strain used to determine the effects of *trans*-regulatory polymorphism on *TDH3* expression.

| Strain Name | Collection Number |
|---|---|
| DBVPG6765 | PJW1015 |
| SK1 | PJW1016 |
| Y55 | PJW1017 |
| YPS128 | PJW1018 |
| DBVPG1373 | PJW1019 |
| DBVPG1853 | PJW1020 |
| YPS606 | PJW1021 |
| L-1374 | PJW1022 |
| L-1528 | PJW1023 |
| Y12 | PJW1024 |
| DBVPG1106 | PJW1025 |
| UWOPS83-787.3 | PJW1026 |
| UWOPS87-2421 | PJW1027 |
| NCYC361 | PJW1028 |
| 322134S | PJW1029 |
| 273614N | PJW1030 |
| YJM978 | PJW1031 |
| UWOPS03-461.4 | PJW1032 |
| UWOPS05-217.3 | PJW1033 |
| S288c | PJW1034 |
| W303 | PJW1035 |
| UWOPS05-227.2 | PJW1036 |
| DBVPG6040 | PJW1037 |
| YIIc17_E5 | PJW1038 |
| YJM981 | PJW1039 |
| YJM975 | PJW1040 |
| NCYC110 | PJW1041 |
| CLIB272 | PJW1042 |
| YJM145 | PJW1043 |

| Strain Name | Collection Number |
|---|---|
| YJM280 | PJW1044 |
| YJM320 | PJW1045 |
| YJM326 | PJW1046 |
| YJM413 | PJW1047 |
| YJM421 | PJW1048 |
| YJM434 | PJW1049 |
| YJM436 | PJW1050 |
| YJM454 | PJW1051 |
| CECT10109 | PJW1052 |
| DBVPG3591 | PJW1053 |
| DBVPG4651 | PJW1054 |
| K12 | PJW1062 |
| YJM269 | PJW1063 |
| BY4716 | PJW1064 |
| A364A | PJW1065 |
| CENPK | PJW1066 |
| CLIB154 | PJW1067 |
| CLIB157 | PJW1068 |
| CLIB219 | PJW1069 |
| DBVPG1399 | PJW1070 |
| I14 | PJW1071 |
| M22 | PJW1072 |
| RM11 | PJW1073 |
| T73 | PJW1074 |
| UC8 | PJW1075 |
| WE372 | PJW1076 |
| Y9J | PJW1077 |
| S288c Copy 2 | PJW1078 |
| BY4716 Copy 2 | PJW1079 |

**Table S2 Barcode sequences used for *TDH3 trans*-regulatory eQTL mapping**
Samples sequenced to determine allele frequency shifts. Selection refers to whether the sample was selected for either high (H) or low (L) expression. Time point indicates the number round of meiosis followed by the number of round of phenotypic selection. Barcodes names refer to those provided by Illumina

| Strain | Selection | Time point | i7 Index | i5 Index |
|---|---|---|---|---|
| PJW1057 | L | 3.1 | N701 | S502 |
| PJW1057 | H | 3.1 | N702 | S502 |
| PJW1016 | L | 3.1 | N703 | S502 |
| PJW1016 | H | 3.1 | N704 | S502 |
| PJW1072 | L | 3.1 | N705 | S502 |
| PJW1072 | H | 3.1 | N706 | S502 |
| PJW1057 | L | 3.2 | N707 | S502 |
| PJW1057 | H | 3.2 | N708 | S502 |
| PJW1016 | L | 3.2 | N709 | S502 |
| PJW1016 | H | 3.2 | N710 | S502 |
| PJW1072 | L | 3.2 | N711 | S502 |
| PJW1072 | H | 3.2 | N712 | S502 |
| PJW1057 | L | 3.3 | N701 | S508 |
| PJW1057 | H | 3.3 | N702 | S508 |
| PJW1016 | L | 3.3 | N703 | S508 |
| PJW1016 | H | 3.3 | N704 | S508 |
| PJW1072 | L | 3.3 | N705 | S508 |
| PJW1072 | H | 3.3 | N706 | S508 |
| PJW1057 | L | 1.1 | N707 | S508 |
| PJW1057 | H | 1.1 | N708 | S508 |
| PJW1016 | L | 1.1 | N709 | S508 |
| PJW1016 | H | 1.1 | N710 | S508 |
| PJW1072 | L | 1.1 | N711 | S508 |
| PJW1072 | H | 1.1 | N712 | S508 |
| PJW1057 | L | 1.2 | N701 | S517 |
| PJW1057 | H | 1.2 | N702 | S517 |
| PJW1016 | L | 1.2 | N703 | S517 |
| PJW1016 | H | 1.2 | N704 | S517 |
| PJW1072 | L | 1.2 | N705 | S517 |
| PJW1072 | H | 1.2 | N706 | S517 |
| PJW1057 | L | 1.3 | N707 | S517 |
| PJW1057 | H | 1.3 | N708 | S517 |
| PJW1016 | L | 1.3 | N709 | S517 |
| PJW1016 | H | 1.3 | N710 | S517 |
| PJW1072 | L | 1.3 | N711 | S517 |
| PJW1072 | H | 1.3 | N712 | S517 |

**Table S3 Primer sequences for tracking sporulation and petite QTN by pyrosequencing**
[1]psq R and F primers used for PCR; pyro primer used for sequencing. [2] /5Biosg/ designates biotinylated primer

| Primer Number | Primer Name[1] | Sequence[2] | Gene |
|---|---|---|---|
| 1530 | RME1_psq_R_bio | /5Biosg/GCACTCTGGCCTTTGTTCTC | RME1 |
| 1529 | RME1_psq_F | TGCTTCGTCACGTAAAATGG | RME1 |
| 1538 | RME1_pyro_F | AAGTGGCCGGGCATGTA | RME1 |
| 1532 | TAO3_psq_F_bio | /5Biosg/TCTCCTTGGGTTTGTATGGTTT | TAO3 |
| 1533 | TAO3_psq_R | GAGAGAGCAGTTCGGCAAAT | TAO3 |
| 1534 | TAO3_pyro_R | AGAATTATGTAATTTCGTTT | TAO3 |
| 1536 | MKT1_psq_R_bio | /5Biosg/TGGCTCTTGGGGTTGAATAG | MKT1 |
| 1535 | MKT1_psq_F | TCCTATGCCATTGAGGCTCT | MKT1 |
| 1537 | MKT1_pyro_F | TCTGAATAATTGTACCCTGG | MKT1 |
| 1588 | SAL1_psq_F_bio | /5Biosg/CATTGCTGGTGGTTTAGCTG | SAL1 |
| 1589 | SAL1_psq_R | TGTGGTAGGTTCAGGGTCTTTG | SAL1 |
| 1590 | SAL1_pyro_R | CACCTCTGTAAAATAATCTG | SAL1 |
| 1591 | CAT5_psq_F_bio | /5Biosg/AGTACTTCGTGTTGGCTCATAGGT | CAT5 |
| 1592 | CAT5_psq_R | AGTGCCCTCCGATTACTGTC | CAT5 |
| 1593 | CAT5_pyro_R | TGATGTATCTCCTGGTCC | CAT5 |
| 1594 | MIP1_psq_F_bio | /5Biosg/CCAATTTGTAGTCCCCAGTTGTAA | MIP1 |
| 1595 | MIP1_psq_R | CCTGGAGGAGCTTTGACTTGAGT | MIP1 |
| 1596 | MIP1_pyro_R | GGATGCGGTTAACCA | MIP1 |

**Table S4 Sequences and dispensation order for pyrosequencing**

[1]Bases on either side of / designates alternative alleles

| Gene | Sequence to analyze[1] | Dispensation order | S228c Allele | Alt Allele |
|---|---|---|---|---|
| RME1 | AT/CAAATATATCACCGTATTTTC | GATCGATATATCA | - | A |
| TAO3 | G/CAACAGCTGAAA | TGCTACAG | C | G |
| MKT1 | A/GTATAGACG | TAGCTATAG | A | G |
| SAL1 | AC/GCCCC/ACCCTCA | GATCGCCACCGTCA | T | C |
| CAT5 | CAC/TATGTGC/TTTT | GCAGTCGATGT | C | G |
| MIP1 | CGC/TATTTTCCA | GCGACTGATTCA | C | T |

**References**


1. Rausher MD, Delph LF (2015) Commentary: When does understanding phenotypic evolution require identification of the underlying genes? *Evolution* 69(7):1655–1664.
2. Gerke JP, Chen CTL, Cohen B a (2006) Natural isolates of *Saccharomyces cerevisiae* display complex genetic variation in sporulation efficiency. *Genetics* 174(2):985–97.
3. Deutschbauer AM, Davis RW (2005) Quantitative trait loci mapped to single-nucleotide resolution in yeast. *Nat Genet* 37(12):1333–40.
4. Chen XJ, Clark-Walker GD (1999) The petite mutation in yeasts: 50 years on. *Int Rev Cytol* 194:197–238.
5. Tong a H, et al. (2001) Systematic genetic analysis with ordered arrays of yeast deletion mutants. *Science* 294(5550):2364–8.
6. Dimitrov LN, Brem RB, Kruglyak L, Gottschling DE (2009) Polymorphisms in multiple genes contribute to the spontaneous mitochondrial genome instability of *Saccharomyces cerevisiae* S288C strains. *Genetics* 183(1):365–83.
7. Storici F, Resnick M a M (2006) The delitto perfetto approach to in vivo site-directed mutagenesis and chromosome rearrangements with synthetic oligonucleotides in yeast. *Methods Enzymol* 409(1976):329–45.
8. Chin BL, Frizzell M a., Timberlake WE, Fink GR (2012) FASTER MT: Isolation of Pure Populations of a and Ascospores from Saccharomyces cerevisiae. *G3 Genes|Genomes|Genetics* 2(4):449–452.
9. Wittkopp PJ (2012) Using Pyrosequencing to Measure Allele-Specific mRNA Abundance and Infer the Effects of *Cis*- and *Trans*-regulatory Differences. *Molecular Methods for Evolutionary Genetics*, Methods in Molecular Biology., eds Orgogozo V, Rockman M V. (Humana Press, Totowa, NJ). doi:10.1007/978-1-61779-228-1.
