## Supplementary material for "Complex polymorphic genetic architecture underlies *trans*-regulatory variation among strains of *Saccharomyces cerevisiae*": Data S1: Analysis.1-Data.Cleaning.R

```
################################################
# R script for cleaning of flow cytometry data #
################################################

# 1 - Necessary packages
# 2 - Functions
# 3A - Cleaning of natural variation trans effects
# 3B - Cleaning of natural variation cis effects
# 3C - Cleaning of mutation trans effects set 1
# 3D - Cleaning of mutation trans effects set 2

################
# 1 - PACKAGES #
################

library(flowCore)
library(flowClust)
library(flowViz)
library(pcaPP)
library(mixtools)
library(plyr)
library(robustlmm)
library(plotrix)
library(moments)
library(MASS)
library(permute)
library(Hmisc)

box <- graphics::box

###############
# 2-FUNCTIONS #
###############
#Set source directory
source.dir <- "/Path.to.functions"
setwd(source.dir)

source("Functions.R")

##################
# 3 - CLEAN DATA #
##################

######################
# 3A - Trans Natural #
######################
#Set working directory
parent.dir <- "/Path.to.data"
setwd(parent.dir)

#Load experiment setup
SETUP <- read.table("TRANS.NATURAL.LAYOUT.txt",header=TRUE,as.is=TRUE)

#Load paths of FCS files
FILENAMES <- list.files(".",pattern=".fcs",recursive=TRUE,include.dirs=TRUE)
SETUP[,"FILENAMES"] <- FILENAMES

#Determine Hard Gates
GATES <- GATE.CALIB(FILENAMES[1],logFSC.H.MAX=6.6)

#Clean Data
TRANS.N.OUTPUT <- adply(apply(SETUP,1,CLEANING,GATES=GATES),1)
TRANS.N.OUTPUT <- cbind(SETUP,TRANS.N.OUTPUT[,-1])
TRANS.N.OUTPUT[,"FILENAMES"] <- NULL
write.table(TRANS.N.OUTPUT,"TRANS.N.CLEAN.txt",quote=FALSE,sep="\t",row.names=FALSE)

#Plate Correction
PLATE.CORRECT(DATA.NAME="TRANS.N",CTRL.STRAIN = "Y1",PLOT=TRUE)

#Strain Estimates
STRAIN.ESTIMATES(DATA.NAME = "TRANS.N", WT.STRAIN = "Y1002", NEG.STRAIN = "Y880", N.PERM= 10000)

############
# 3B - Cis #
############
#Set working directory
parent.dir <- "/Path.to.data"
setwd(parent.dir)

#Load experiment setup
SETUP <- read.table("CIS.NATURAL.LAYOUT.txt",header=TRUE,as.is=TRUE)

#Load paths of FCS files
FILENAMES <- list.files(".",pattern=".fcs",recursive=TRUE,include.dirs=TRUE)
SETUP[,"FILENAMES"] <- FILENAMES

#Determine Hard Gates
GATES <- GATE.CALIB(FILENAMES[1],logFSC.H.MAX=6.8, logFSC.A.MAX = 6.3, logFSC.A.MIN = 5.3)

#Clean Data
CIS.OUTPUT <- adply(apply(SETUP,1,CLEANING,GATES=GATES),1)
CIS.OUTPUT <- cbind(SETUP,CIS.OUTPUT[,-1])
CIS.OUTPUT[,"FILENAMES"] <- NULL
write.table(CIS.OUTPUT,"CIS.CLEAN.txt",quote=FALSE,sep="\t",row.names=FALSE)

#Plate Correction
PLATE.CORRECT(DATA.NAME="CIS",CTRL.STRAIN = "CTRL",PLOT=TRUE)

#Strain Estimates
STRAIN.ESTIMATES(DATA.NAME = "CIS", WT.STRAIN = "WT.Y1.1", NEG.STRAIN = "Y880", N.PERM= 10000)

######################
# 3C - Trans Round 1 #
######################
#Set working directory
parent.dir <- "/Path.to.data"
setwd(parent.dir)

#Load experiment setup
SETUP <- read.table("TRANS.MUTATION.1.LAYOUT.txt",header=TRUE,as.is=TRUE)

#Load paths of FCS files
FILENAMES <- list.files(".",pattern=".fcs",recursive=TRUE,include.dirs=TRUE)
SETUP[,"FILENAMES"] <- FILENAMES

#Determine Hard Gates
GATES <- GATE.CALIB(FILENAMES[1],logFSC.H.MAX=6.5, logFSC.A.MAX = 6.2, logFSC.A.MIN = 5.3)

#Clean Data
TRANS.1.OUTPUT <- adply(apply(SETUP,1,CLEANING,GATES=GATES),1)
TRANS.1.OUTPUT <- cbind(SETUP,TRANS.1.OUTPUT[,-1])
TRANS.1.OUTPUT[,"FILENAMES"] <- NULL
write.table(TRANS.1.OUTPUT,"TRANS.1.CLEAN.txt",quote=FALSE,sep="\t",row.names=FALSE)

#Plate Correction
PLATE.CORRECT(DATA.NAME="TRANS.1",CTRL.STRAIN = "CTRL",PLOT=TRUE)

#Strain Estimates
STRAIN.ESTIMATES(DATA.NAME = "TRANS.1", WT.STRAIN = "SHAM.ALL", NEG.STRAIN = "Y978", N.PERM= 10000)

######################
# 3D - Trans Round 2 #
######################
#Set working directory
parent.dir <- "/Path.to.data"
setwd(parent.dir)

#Load experiment setup
SETUP <- read.table("TRANS.MUTATION.2.LAYOUT.txt",header=TRUE,as.is=TRUE)

#Load paths of FCS files
FILENAMES <- list.files(".",pattern=".fcs",recursive=TRUE,include.dirs=TRUE)
SETUP[,"FILENAMES"] <- FILENAMES

#Determine Hard Gates
GATES <- GATE.CALIB(FILENAMES[1],logFSC.H.MAX=6.5, logFSC.A.MAX = 6.2, logFSC.A.MIN = 5.3)

#Clean Data
TRANS.2.OUTPUT <- adply(apply(SETUP,1,CLEANING,GATES=GATES),1)
TRANS.2.OUTPUT <- cbind(SETUP,TRANS.2.OUTPUT[,-1])
TRANS.2.OUTPUT[,"FILENAMES"] <- NULL
write.table(TRANS.2.OUTPUT,"TRANS.2.CLEAN.txt",quote=FALSE,sep="\t",row.names=FALSE)

#Plate Correction
PLATE.CORRECT(DATA.NAME="TRANS.2",CTRL.STRAIN = "CTRL",PLOT=TRUE)

#Strain Estimates
STRAIN.ESTIMATES(DATA.NAME = "TRANS.2", WT.STRAIN = "SHAM.ALL", NEG.STRAIN = "Y978", N.PERM= 10000)
```
