## Supplementary material for "Complex polymorphic genetic architecture underlies *trans*-regulatory variation among strains of *Saccharomyces cerevisiae*": Data S1: Analysis.2-Figures.R

```
###############################################
#  R script for analyzing flow cytometry data #
###############################################

# 1 - Packages
# 2 - Functions 
# 3 - Data
# 4 - Phylogeny
# 5 - Ancestral State Reconstruction
# 6 - Mean-Noise Correlation
# 7 - Mutational Distributions
# 8 - Test for selection using mutational effects
# 9 - Phylogenetic Constraint
# 10 - Fitness Effects
# 11 - Natural variation in TDH3 promoter
# 12 - Brownian Motion and Ornstein Uhlenbeck Modeling

################
# 1 - Packages #
################
library(plotrix)
library(ape)
library(phytools)
library(plyr)
library(Hmisc)
library(numDeriv)
library(mvtnorm)
library(quantmod)

#################
# 2 - FUNCTIONS #
#################
#Set source directory
source.dir <- "/Path.to.functions"
setwd(source.dir)

source("Functions.R")

############
# 3 - DATA #
############
source.dir <- "/Path.to.data"
setwd(source.dir)

#Read in data for natural strains
DATA <- read.table("FILTER.TRANS.N.DATA.txt",header=TRUE)
NV.STRAINS  <- read.table("NV.Strains.txt",header=TRUE)

#Drop NCYC110, EM93, YIIc17_E5, and DBVPG3591 due to poor data quality
DATA <- droplevels(subset(DATA,!DATA$STRAIN %in% c("Y1041","Y1055","Y1038","Y1053")))

##Average expression and noise for each strain with SE
MEAN.STRAIN     <- tapply(DATA$YFP.MEDIAN.RELATIVE,DATA$STRAIN,FUN=mean)
NOISE.STRAIN    <- tapply(DATA$YFP.CV.RELATIVE,DATA$STRAIN,FUN=mean)
MEAN.STRAIN.SE  <- tapply(DATA$YFP.MEDIAN.RELATIVE,DATA$STRAIN,FUN=std.error)
NOISE.STRAIN.SE <- tapply(DATA$YFP.CV.RELATIVE,DATA$STRAIN,FUN=std.error)

#Combine with names of natural isolates
NV.MEAN     <- droplevels(merge(NV.STRAINS,MEAN.STRAIN,by.x=1,by.y=0))
NV.NOISE    <- droplevels(merge(NV.STRAINS,NOISE.STRAIN,by.x=1,by.y=0))
NV.MEAN.SE  <- droplevels(merge(NV.STRAINS,MEAN.STRAIN.SE,by.x=1,by.y=0))
NV.NOISE.SE <- droplevels(merge(NV.STRAINS,NOISE.STRAIN.SE,by.x=1,by.y=0))

#Clean up names
NV.MEAN$y     <- unname(NV.MEAN$y)
NV.NOISE$y    <- unname(NV.NOISE$y)
NV.MEAN.SE$y  <- unname(NV.MEAN.SE$y)
NV.NOISE.SE$y <- unname(NV.NOISE.SE$y)
colnames(NV.MEAN)     <- c("STRAIN","NAME","EFFECT")
colnames(NV.NOISE)    <- c("STRAIN","NAME","EFFECT")
colnames(NV.MEAN.SE)  <- c("STRAIN","NAME","SE")
colnames(NV.NOISE.SE) <- c("STRAIN","NAME","SE")

#################
# 4 - PHYLOGENY #
#################

##Code to clean up phylogeny
##Phylogeny from Maclean et al. 2017 MBE
#PHYLOGENY <- read.tree("RC6.hom.boot.nwk")
##Only keep strains with expression data
#DROP <- !(PHYLOGENY$tip.label %in% NV.MEAN$NAME)
#PHYLOGENY.PRUNE <- drop.tip(PHYLOGENY, PHYLOGENY$tip.label[DROP])
#NV.MEAN <- NV.MEAN[NV.MEAN$NAME %in% PHYLOGENY.PRUNE$tip.label,]
#NV.NOISE <- NV.NOISE[NV.NOISE$NAME %in% PHYLOGENY.PRUNE$tip.label,]

##Reroot phylogeny
#PHYLOGENY.PRUNE <- reroot(PHYLOGENY.PRUNE,60,0.0041)
#PHYLOGENY.PRUNE <- rotate(PHYLOGENY.PRUNE,57)
#PHYLOGENY.PRUNE <- rotate(PHYLOGENY.PRUNE,71)
#PHYLOGENY.PRUNE <- rotate(PHYLOGENY.PRUNE,61)
#PHYLOGENY.PRUNE <- rotate(PHYLOGENY.PRUNE,62)
#PHYLOGENY.PRUNE <- rotate(PHYLOGENY.PRUNE,65)
#PHYLOGENY.PRUNE <- rotate(PHYLOGENY.PRUNE,67)
#PHYLOGENY.PRUNE <- rotate(PHYLOGENY.PRUNE,69)
##12071326 sites in S228c reference genome
##421773 segregating sites among strains used in Maclean et al. 2017 MBE
#PHYLOGENY.PRUNE$edge.length <- PHYLOGENY.PRUNE$edge.length/(12071326/421773)
#plot(PHYLOGENY.PRUNE,cex=0.5,main="Mean Expression",x.lim=0.004)
#add.scale.bar(x=0,y=0,length=0.001)
#write.tree(PHYLOGENY.PRUNE,file = "Sc.nwk")

#Read in phylogeny
PHYLOGENY.PRUNE <- read.tree("Sc.nwk")
PHYLOGENY.PRUNE$node.label <- 57:111
NV.MEAN <- NV.MEAN[NV.MEAN$NAME %in% PHYLOGENY.PRUNE$tip.label,]
NV.NOISE <- NV.NOISE[NV.NOISE$NAME %in% PHYLOGENY.PRUNE$tip.label,]

##Show expression values on phylogeny
plot(PHYLOGENY.PRUNE,cex=0.5,main="Mean Expression",x.lim=0.006)
abline(v =  0.005)
abline(v =  0.0045)
abline(v =  0.0055)

I <- 1:length(PHYLOGENY.PRUNE$tip.label)
NAME <- numeric(length(I))
for(i in I) {
	NAME[i]   <- PHYLOGENY.PRUNE$tip.label[i]
	DATA.STRAIN <- DATA[which(DATA$STRAIN %in% as.character(NV.MEAN[which(NV.MEAN$NAME == NAME[i]),]$STRAIN)),]
	plotCI(log2(mean(DATA.STRAIN$YFP.MEDIAN.RELATIVE))*.0005 + .005,i,ui = log2(mean(DATA.STRAIN$YFP.MEDIAN.RELATIVE) + 1.96*sd(DATA.STRAIN$YFP.MEDIAN.RELATIVE)/nrow(DATA.STRAIN))*0.0005 + .005, li = log2(mean(DATA.STRAIN$YFP.MEDIAN.RELATIVE) - 1.96*sd(DATA.STRAIN$YFP.MEDIAN.RELATIVE)/nrow(DATA.STRAIN))*0.0005 + .005,pch=19,cex=0.5,add=TRUE,err="x")
}

#####################################
# 5 - Ancstral State Reconstruction #
#####################################
#Combine replicates of S288c and combine replicates of BY
BY.STRAINS <- as.character(NV.MEAN.SE[which(NV.MEAN.SE$NAME == "BY4716"),]$STRAIN)
S288c.STRAINS <- as.character(NV.MEAN.SE[which(NV.MEAN.SE$NAME == "S288c"),]$STRAIN)

NV.MEAN[which(NV.MEAN$NAME == "S288c")[1],]$EFFECT <- mean(subset(DATA,DATA$STRAIN %in% S288c.STRAINS)$YFP.MEDIAN.RELATIVE)
NV.MEAN <- NV.MEAN[-which(NV.MEAN$NAME == "S288c")[2],]
NV.MEAN[which(NV.MEAN$NAME == "BY4716")[1],]$EFFECT <- mean(subset(DATA,DATA$STRAIN %in% BY.STRAINS)$YFP.MEDIAN.RELATIVE)
NV.MEAN <- NV.MEAN[-which(NV.MEAN$NAME == "BY4716")[2],]

NV.MEAN.SE[which(NV.MEAN.SE$NAME == "S288c")[1],]$SE <- std.error(subset(DATA,DATA$STRAIN %in% S288c.STRAINS)$YFP.MEDIAN.RELATIVE)
NV.MEAN.SE <- NV.MEAN.SE[-which(NV.MEAN.SE$NAME == "S288c")[2],]
NV.MEAN.SE[which(NV.MEAN.SE$NAME == "BY4716")[1],]$SE <- std.error(subset(DATA,DATA$STRAIN %in% BY.STRAINS)$YFP.MEDIAN.RELATIVE)
NV.MEAN.SE <- NV.MEAN.SE[-which(NV.MEAN.SE$NAME == "BY4716")[2],]

NV.NOISE[which(NV.NOISE$NAME == "S288c")[1],]$EFFECT <- mean(subset(DATA,DATA$STRAIN %in% S288c.STRAINS)$YFP.CV.RELATIVE)
NV.NOISE <- NV.NOISE[-which(NV.NOISE$NAME == "S288c")[2],]
NV.NOISE[which(NV.NOISE$NAME == "BY4716")[1],]$EFFECT <- mean(subset(DATA,DATA$STRAIN %in% BY.STRAINS)$YFP.CV.RELATIVE)
NV.NOISE <- NV.NOISE[-which(NV.NOISE$NAME == "BY4716")[2],]

NV.NOISE.SE[which(NV.NOISE.SE$NAME == "S288c")[1],]$SE <- std.error(subset(DATA,DATA$STRAIN %in% S288c.STRAINS)$YFP.CV.RELATIVE)
NV.NOISE.SE <- NV.NOISE.SE[-which(NV.NOISE.SE$NAME == "S288c")[2],]
NV.NOISE.SE[which(NV.NOISE.SE$NAME == "BY4716")[1],]$SE <- std.error(subset(DATA,DATA$STRAIN %in% BY.STRAINS)$YFP.CV.RELATIVE)
NV.NOISE.SE <- NV.NOISE.SE[-which(NV.NOISE.SE$NAME == "BY4716")[2],]

row.names(NV.MEAN) <- NV.MEAN$NAME
row.names(NV.NOISE) <- NV.NOISE$NAME

#Ancestral state estimates
ASR.MEAN  <- ace(NV.MEAN[PHYLOGENY.PRUNE$tip.label,]$EFFECT, PHYLOGENY.PRUNE,type="continuous",method="pic")
ASR.NOISE <- ace(NV.NOISE[PHYLOGENY.PRUNE$tip.label,]$EFFECT,PHYLOGENY.PRUNE,type="continuous",method="pic")

#Phylogeny labels
plot(PHYLOGENY.PRUNE,cex=0.5,main="")
edgelabels(1:110,1:110,cex=0.6)
nodelabels(1:111,1:111,cex=0.6)

#Mean
NODES.MEAN <- c(NV.MEAN[PHYLOGENY.PRUNE$tip.label,]$EFFECT,ASR.MEAN$ace)
names(NODES.MEAN) <- 1:length(NODES.MEAN)
N <- 0
P <- 0

I <- 1:(length(NODES.MEAN)-1)
DIFF.MEAN <- numeric(length(I))
for(i in I) {
	A <- NODES.MEAN[PHYLOGENY.PRUNE$edge[i,2]]
	D <- NODES.MEAN[PHYLOGENY.PRUNE$edge[i,1]]
	E <- PHYLOGENY.PRUNE$edge.length[i]*(D-A)
	if(E < 0) {
		N <- N + E
	}else if (E > 0) {
		P <- P + E
	}
	DIFF.MEAN[i] <- D-A
}
hist(DIFF.MEAN,breaks=40,xlab="Expression",main="")

###Plot phylogeny with ancestral expression estimates
MEAN.EFFECT <- log2(NV.MEAN$EFFECT)
names(MEAN.EFFECT) <- NV.MEAN$NAME
MEAN.PLOT <- contMap(PHYLOGENY.PRUNE,100*MEAN.EFFECT,plot=FALSE,lims=c(-100,100))

n <- length(MEAN.PLOT$cols)
COLOR <- colorRampPalette(c("white","deepskyblue","royalblue","black"), space="rgb")(n)

MEAN.PLOT$cols[1:n] <- COLOR
plot(MEAN.PLOT,fsize=0.6,outline=FALSE,lwd=5,sig=0)
add.scale.bar(x=0,y=0,length=0.001)

#Add points at tips with final character state
lastPP <- get("last_plot.phylo", envir = .PlotPhyloEnv)
tip <- 1:lastPP$Ntip
XX <- lastPP$xx[tip]
YY <- lastPP$yy[tip]

C <- COLOR[1000*round((MEAN.EFFECT - -1)/(1 - -1),4)+1]
ORDER <- match(names(MEAN.EFFECT), PHYLOGENY.PRUNE$tip.label)
points(XX[ORDER],YY[ORDER],pch=19,col=C,cex=1)

###Plot phylogeny with ancestral expression noise estimates
COLOR <- colorRampPalette(c("white","lightgreen","forestgreen","black"), space="rgb")(n)
NOISE.EFFECT <- log2(NV.NOISE$EFFECT)
names(NOISE.EFFECT) <- NV.NOISE$NAME
NOISE.PLOT <- contMap(PHYLOGENY.PRUNE,100*NOISE.EFFECT,plot=FALSE,lims=c(-200,200))
n <- length(NOISE.PLOT$cols)
NOISE.PLOT$cols[1:n] <- COLOR
plot(NOISE.PLOT,fsize=0.6,outline=FALSE,lwd=5,sig=0)
add.scale.bar(x=0,y=0,length=0.001)

#Add points at tips with final character state
lastPP <- get("last_plot.phylo", envir = .PlotPhyloEnv)
tip <- 1:lastPP$Ntip
XX <- lastPP$xx[tip]
YY <- lastPP$yy[tip]

C <- COLOR[1000*round((NOISE.EFFECT- - 2)/(2 - -2),4)+1]
ORDER <- match(names(NOISE.EFFECT), PHYLOGENY.PRUNE$tip.label)
points(XX[ORDER],YY[ORDER],pch=19,col=C,cex=1)

###################
# 6 - Correlation #
###################
#Measured Effects
cor.test(NV.MEAN$EFFECT,NV.NOISE$EFFECT)
par(pty="s")
par(mar=c(5,5,5,5))
plot(x=NULL,y=NULL,xlim=c(0,2),ylim=c(0,2),xlab="Mean",ylab="Noise")
I <- 1:length(PHYLOGENY.PRUNE$tip.label)
for(i in I) {
  NAME[i]   <- PHYLOGENY.PRUNE$tip.label[i]
  DATA.STRAIN <- DATA[which(DATA$STRAIN %in% as.character(NV.NOISE[which(NV.NOISE$NAME == NAME[i]),]$STRAIN)),]
  plotCI((mean(DATA.STRAIN$YFP.MEDIAN.RELATIVE)),(mean(DATA.STRAIN$YFP.CV.RELATIVE)),uiw=1.96*sd(DATA.STRAIN$YFP.MEDIAN.RELATIVE)/nrow(DATA.STRAIN), liw =1.96*sd(DATA.STRAIN$YFP.MEDIAN.RELATIVE)/nrow(DATA.STRAIN),pch=19,cex=0.5,add=TRUE,err="x")
  plotCI((mean(DATA.STRAIN$YFP.MEDIAN.RELATIVE)),(mean(DATA.STRAIN$YFP.CV.RELATIVE)),uiw=1.96*sd(DATA.STRAIN$YFP.CV.RELATIVE)/nrow(DATA.STRAIN),     liw =1.96*sd(DATA.STRAIN$YFP.CV.RELATIVE)/nrow(DATA.STRAIN),    pch=19,cex=0.5,add=TRUE,err="y")
}
abline(h=1)
abline(v=1)

################################
# 7 - Mutational distributions #
################################
###Trans set 1
TRANS.DATA.1 <- read.table("FILTER.TRANS.1.DATA.txt",header=TRUE)

##Low tail biased effects on mean and noise
TRANS.MUTANTS.LOW <- droplevels(subset(TRANS.DATA.1,TRANS.DATA.1$TREATMENT == "EMS" & TRANS.DATA.1$TAIL == "LOW"))
TRANS.STRAIN.LOW <- data.frame(matrix(0,nrow=length(levels(TRANS.MUTANTS.LOW$STRAIN)),ncol=2))
colnames(TRANS.STRAIN.LOW) <- c("MEAN","NOISE")
rownames(TRANS.STRAIN.LOW) <- levels(TRANS.MUTANTS.LOW$STRAIN)

TRANS.STRAIN.LOW$MEAN  <- tapply(TRANS.MUTANTS.LOW$YFP.MEDIAN.RELATIVE,TRANS.MUTANTS.LOW$STRAIN,FUN=median)
TRANS.STRAIN.LOW$NOISE <- tapply(TRANS.MUTANTS.LOW$YFP.CV.RELATIVE,TRANS.MUTANTS.LOW$STRAIN,FUN=median)

##High tail biased effects on mean and noise
TRANS.MUTANTS.HIGH <- droplevels(subset(TRANS.DATA.1,TRANS.DATA.1$TAIL == "HIGH"))
TRANS.STRAIN.HIGH <- data.frame(matrix(0,nrow=length(levels(TRANS.MUTANTS.HIGH$STRAIN)),ncol=2))
colnames(TRANS.STRAIN.HIGH) <- c("MEAN","NOISE")
rownames(TRANS.STRAIN.HIGH) <- levels(TRANS.MUTANTS.HIGH$STRAIN)

TRANS.STRAIN.HIGH$MEAN  <- tapply(TRANS.MUTANTS.HIGH$YFP.MEDIAN.RELATIVE,TRANS.MUTANTS.HIGH$STRAIN,FUN=median)
TRANS.STRAIN.HIGH$NOISE <- tapply(TRANS.MUTANTS.HIGH$YFP.CV.RELATIVE,TRANS.MUTANTS.HIGH$STRAIN,FUN=median)

###Trans set 2
TRANS.DATA.2 <- read.table("FILTER.TRANS.2.DATA.txt",header=TRUE)

#Unbiased sample effects on mean and noise 
TRANS.DATA.2 <- droplevels(subset(TRANS.DATA.2,TRANS.DATA.2$TREATMENT == "EMS" & TRANS.DATA.2$TAIL == "ALL" ))
TRANS.STRAIN.2 <- data.frame(matrix(0,nrow=length(levels(TRANS.DATA.2$STRAIN)),ncol=2))
colnames(TRANS.STRAIN.2) <- c("MEAN","NOISE")
rownames(TRANS.STRAIN.2) <- levels(TRANS.DATA.2$STRAIN)

TRANS.STRAIN.2$MEAN      <- tapply(TRANS.DATA.2$YFP.MEDIAN.RELATIVE,TRANS.DATA.2$STRAIN,FUN=median)
TRANS.STRAIN.2$MEAN.MAD  <- tapply(TRANS.DATA.2$YFP.MEDIAN.RELATIVE,TRANS.DATA.2$STRAIN,FUN=mad)
TRANS.STRAIN.2$NOISE     <- tapply(TRANS.DATA.2$YFP.CV.RELATIVE,TRANS.DATA.2$STRAIN,FUN=median)
TRANS.STRAIN.2$NOISE.MAD <- tapply(TRANS.DATA.2$YFP.CV.RELATIVE,TRANS.DATA.2$STRAIN,FUN=mad)

#Combine samples from tails
TRANS.STRAIN.TAIL <- droplevels(rbind(TRANS.STRAIN.LOW,TRANS.STRAIN.HIGH))

##############################################
# 8 - Compare mutation and natural variation #
##############################################
#Final phylogeny based on 311287 sites
SITES <- 311287

#Genome is 12071326 bp
GENOME <- 12071326

#Correct phylogeny distances
DISTANCE <- PHYLOGENY.PRUNE$edge.length
DIST <- dist.nodes(PHYLOGENY.PRUNE)

####Test for selection####
#Get observed changes in expression and distance along each branch
I <- 1:110
START.DIST <- numeric(length(I))
STOP.DIST <- numeric(length(I))
START.VALUE <- numeric(length(I))
STOP.VALUE <- numeric(length(I))
for(i in I){
  START.DIST[i]  <- DIST[57,PHYLOGENY.PRUNE$edge[i,1]]*GENOME/43
  STOP.DIST[i]   <- DIST[57,PHYLOGENY.PRUNE$edge[i,2]]*GENOME/43
  START.VALUE[i] <- log(NODES.MEAN[PHYLOGENY.PRUNE$edge[i,1]],2)
  STOP.VALUE[i]  <- log(NODES.MEAN[PHYLOGENY.PRUNE$edge[i,2]],2)
}

#Get mutation effects over time
MUTATION.NUMBER.EQUAL <- c(1,seq(10,1600,10))
I <- 1:length(MUTATION.NUMBER.EQUAL )
J <- 1:10000
MEAN.PRED.TIME <- matrix(0,nrow=length(I),ncol=length(J))
for(j in J) {
  for(i in I) {
    x <- sample(TRANS.STRAIN.2$MEAN,MUTATION.NUMBER.EQUAL[i],replace=TRUE)
    MEAN.PRED.TIME[i,j] <- prod(x)
  }
}

I <- 1:110
#Calculate likelihood of observed data
OBSERVED.LOG.LIKELIHOOD <- numeric(length(I))
for(i in I) {
  x1 <- DMUTATION.MEAN(seq(1+round_any(STOP.VALUE[i]-START.VALUE[i],.001,floor),1+round_any(STOP.VALUE[i]-START.VALUE[i],.001,ceiling),0.001), MEAN.PRED.TIME[round_any((STOP.DIST[i]-START.DIST[i]), 10, floor)/10 + 1,])
  x2 <- DMUTATION.MEAN(seq(1+round_any(STOP.VALUE[i]-START.VALUE[i],.001,floor),1+round_any(STOP.VALUE[i]-START.VALUE[i],.001,ceiling),0.001), MEAN.PRED.TIME[round_any((STOP.DIST[i]-START.DIST[i]), 10, ceiling)/10 + 1,])
  
  OBSERVED.LOG.LIKELIHOOD[i] <- log10(0.01*((1 - (STOP.DIST[i]-START.DIST[i])/round_any(STOP.DIST[i]-START.DIST[i], 10, ceiling))*
                                                (x1[1]*1000*(round_any(STOP.VALUE[i]-START.VALUE[i],.001,ceiling)-(STOP.VALUE[i]-START.VALUE[i])) +
                                                 x1[2]*1000*((STOP.VALUE[i]-START.VALUE[i])-round_any(STOP.VALUE[i]-START.VALUE[i],.001,floor))) +
                                              (((STOP.DIST[i]-START.DIST[i])/round_any(STOP.DIST[i]-START.DIST[i], 10, ceiling))*
                                                 (x2[1]*1000*(round_any(STOP.VALUE[i]-START.VALUE[i],.001,ceiling)-(STOP.VALUE[i]-START.VALUE[i])) +
                                                  x2[2]*1000*((STOP.VALUE[i]-START.VALUE[i])-round_any(STOP.VALUE[i]-START.VALUE[i],.001,floor))))))
}

#Calculate likelihood of mutation
I <- 1:110
J <- 1:100000
NEUTRAL.EXPECTATION <- matrix(0,nrow=length(I),ncol=length(J))
LOG.LIKELIHOOD <- matrix(0,nrow=length(I),ncol=length(J))
for(j in J) {
  for(i in I) {
    x <- sample(TRANS.STRAIN.2$MEAN,max(1,STOP.DIST[i]-START.DIST[i]),replace=TRUE)
    NEUTRAL.EXPECTATION[i,j] <- prod(x)

    #Get densities bracketing distance and change in expression
    x1 <- DMUTATION.MEAN(seq(round_any(NEUTRAL.EXPECTATION[i,j],.001,floor),round_any(NEUTRAL.EXPECTATION[i,j],.001,ceiling),0.001), MEAN.PRED.TIME[round_any(STOP.DIST[i]-START.DIST[i], 10, floor)/10 + 1,])
    x2 <- DMUTATION.MEAN(seq(round_any(NEUTRAL.EXPECTATION[i,j],.001,floor),round_any(NEUTRAL.EXPECTATION[i,j],.001,ceiling),0.001), MEAN.PRED.TIME[round_any(STOP.DIST[i]-START.DIST[i], 10, ceiling)/10 + 1,])
    
    #Weight densities to get likelihood  
    LOG.LIKELIHOOD[i,j] <- log10(0.01*((1 - (STOP.DIST[i]-START.DIST[i])/round_any(STOP.DIST[i]-START.DIST[i], 10, ceiling))*
                      (x1[1]*1000*(round_any(NEUTRAL.EXPECTATION[i,j],.001,ceiling)-NEUTRAL.EXPECTATION[i,j]) +
                       x1[2]*1000*(NEUTRAL.EXPECTATION[i,j]-round_any(NEUTRAL.EXPECTATION[i,j],.001,floor))) +
                      (((STOP.DIST[i]-START.DIST[i])/round_any(STOP.DIST[i]-START.DIST[i], 10, ceiling))*
                      (x2[1]*1000*(round_any(NEUTRAL.EXPECTATION[i,j],.001,ceiling)-NEUTRAL.EXPECTATION[i,j]) +
                       x2[2]*1000*(NEUTRAL.EXPECTATION[i,j]-round_any(NEUTRAL.EXPECTATION[i,j],.001,floor))))))
    
  }
}

#Compare observed likelihood vs. mutation likelihoods with same distances
hist(colSums(LOG.LIKELIHOOD,na.rm=TRUE),breaks=1000,xlim=c(-300,-200),main=NULL,xlab="log(Likelihood)")
abline(v=sum(OBSERVED.LOG.LIKELIHOOD),col="red")

#Calculate likelihood of observed data and mutational effects for ever increasing distances
K <- order(STOP.DIST-START.DIST)
EXP <- matrix(0,nrow=110,ncol=100000)
OBS <- numeric(length(I))
SET <- K[1]
for(i in I) {
  SET <- c(SET,K[i])
  EXP[i,] <- colSums(LOG.LIKELIHOOD[SET,],na.rm=TRUE)
  OBS[i] <- sum(OBSERVED.LOG.LIKELIHOOD[SET])
}

plot(1:110,OBS,type="l",ylim=c(-250,0))
points(1:110,apply(EXP,1,quantile,probs=0.5),type="l",col="blue",lwd=2)
points(1:110,apply(EXP,1,quantile,probs=0.1),type="l",col="royalblue",lwd=1)
points(1:110,apply(EXP,1,quantile,probs=0.9),type="l",col="royalblue",lwd=1)
points(1:110,apply(EXP,1,quantile,probs=0.01),type="l",col="lightblue",lwd=1)
points(1:110,apply(EXP,1,quantile,probs=0.99),type="l",col="lightblue",lwd=1)

#Find break point where observation because more likely than expected
BREAK <- min(which(OBS > apply(EXP,1,quantile,probs=0.99)))
BREAK.DIST <- STOP.DIST[K[BREAK]] - START.DIST[K[BREAK]]
abline(v=BREAK)  

#Plot mutational trajectory 
plot(1,type="n",ylim=c(-4,4),xlim=c(0,700),ylab="log(Expression)",xlab="Distance",xaxt='n')
axis(side=1,at=c(0,5000/42,10000/42,15000/42,20000/42,25000/42,30000/42),labels=c(0,5000,10000,15000,20000,25000,30000),cex.axis=0.6)
QUANTILES <- c(0.05,0.1,0.2,0.3,0.4,0.5,0.6,0.7,0.8,0.9,0.95)
Q <- 1:length(QUANTILES)
for(q in Q) {
  if(QUANTILES[q] == 0.5) {
    points(MUTATION.NUMBER.EQUAL,log(apply(MEAN.PRED.TIME,1,quantile,probs=QUANTILES[q]),2),type="l",col="blue", lwd=2)
  } else {
    points(MUTATION.NUMBER.EQUAL,log(apply(MEAN.PRED.TIME,1,quantile,probs=QUANTILES[q]),2),type="l",col="blue")
  }
}
abline(v = BREAK.DIST)

#Add observed distances, correcting for phylogeny
I <- 1:110
for(i in I){
  if(STOP.DIST[i]-START.DIST[i] < 1) {
  } else {
    points(STOP.DIST[i]-START.DIST[i],STOP.VALUE[i]-START.VALUE[i],pch=1,cex=0.4,col="white")
    points(STOP.DIST[i]-START.DIST[i],STOP.VALUE[i]-START.VALUE[i],pch=19,cex=0.4)
  }
}

#Plot small number of trajectories for figure
J <- 1:10
plot(1,type="n",xlim=c(0,700),ylim=c(-4,4),xaxt="n",xlab="Number of mutations",ylab="log2(Expression)")
axis(side=1,at=c(0,5000/42,10000/42,15000/42,20000/42,25000/42,30000/42),labels=c(0,5000,10000,15000,20000,25000,30000),cex.axis=0.6)
for(j in J) {
  x <- sample(TRANS.STRAIN.2$MEAN,1600,replace=TRUE)
  y <- cumprod(x)
  points(log2(y),type="l")
}

###############################
# 9 - Phylogenetic Constraint #
###############################
MEAN.SIGNAL <- NV.MEAN[,3]
names(MEAN.SIGNAL) <- NV.MEAN[,2]
MEAN.SIGNAL.SE <- NV.MEAN.SE[,3]
names(MEAN.SIGNAL.SE) <- NV.MEAN.SE[,2]
phylosig(PHYLOGENY.PRUNE,MEAN.SIGNAL,method="lambda",test=TRUE,se=MEAN.SIGNAL.SE)
phylosig(PHYLOGENY.PRUNE,MEAN.SIGNAL,method="K",test=TRUE,nsim=10000,se=MEAN.SIGNAL.SE)

NOISE.SIGNAL <- NV.NOISE[,3]
names(NOISE.SIGNAL) <- NV.NOISE[,2]
NOISE.SIGNAL.SE <- NV.NOISE.SE[,3]
names(NOISE.SIGNAL.SE) <- NV.NOISE.SE[,2]
phylosig(PHYLOGENY.PRUNE,NOISE.SIGNAL,method="lambda",test=TRUE,se=NOISE.SIGNAL.SE)
phylosig(PHYLOGENY.PRUNE,NOISE.SIGNAL,method="K",test=TRUE,nsim=10000,se=NOISE.SIGNAL.SE)

#CI estimates for ASR values
plotCI(ASR.MEAN$ace,ui=ASR.MEAN$CI95[,2],li=ASR.MEAN$CI95[,1],ylab="",xlab="",pch=19,cex=0.6)
plotCI(ASR.NOISE$ace,ui=ASR.NOISE$CI95[,2],li=ASR.NOISE$CI95[,1],ylab="",xlab="",pch=19,cex=0.6)

##################################################
# 10 - Fitness Effects based on TDH3 cis mutants #
##################################################
#Borrows approach from Duveau et al. 2018 PNAS
#Load files generated in sections 4 and 5.
Fit <- read.table("SUMMARY.DATA.FITNESS.txt",header=TRUE,as.is=TRUE)
Expression <- read.table("SUMMARY.DATA.EXPRESSION.V2.txt",header=TRUE,as.is=TRUE)
All.Fit <- read.table("Experiment_s.estimates_filtered.txt",header=TRUE,as.is=TRUE)

Expression <- subset(Expression, STRAIN != "Y2675")
Expression[which(Expression$ID == 3),"YFP.CONSTRUCT"] <- "SINGLE" 

All.Fit[which(All.Fit$MUTATION == "URA3" & All.Fit$YFP.CONSTRUCT == "DOUBLE"),"MUTATION"] <- "WT_WT"

for (i in 1:nrow(Expression))
{
  if (Expression[i,"MUTATION"] == "URA3" & Expression[i,"YFP.CONSTRUCT"] == "DOUBLE")
  {
    Expression[i,"MUTATION"] <- "WT_WT"
  }
  
  if (Expression[i,"MUTATION"] == "NEGATIVE")
  {
    Expression[i,"MUTATION"] <- "TDH3.Deletion"
  }
  
}

for (i in 1:nrow(Fit))
{
  if (Fit[i,"MUTATION"] == "URA3" & Fit[i,"YFP.CONSTRUCT"] == "DOUBLE")
  {
    Fit[i,"MUTATION"] <- "WT_WT"
  }
}

Expression <- Expression[order(Expression$ENVIRONMENT,Expression$MUTATION),]
Fit <- Fit[order(Fit$ENVIRONMENT, Fit$MUTATION),]

#Filter out strains with more than two copies of YFP or TDH3 (as measured by pyrosequencing).
GLU.EXPR <- subset(Expression, ENVIRONMENT == "GLUCOSE" & MUTATION %nin% c("TATA.81_TATA.81","m75_TATA.81","m63.m90_m63.m90","TATA.129_TATA.129","TATA.42_TATA.42","WT"))
GLU.FIT <- subset(Fit, ENVIRONMENT == "GLUCOSE" & MUTATION %nin% c("TATA.81_TATA.81","m75_TATA.81","m63.m90_m63.m90","TATA.129_TATA.129","TATA.42_TATA.42","WT"))
GLU.ALL <- subset(All.Fit, ENVIRONMENT == "GLUCOSE" & MUTATION %in% GLU.FIT[,"MUTATION"])

####1. Loess fit#########
WT.1 <- subset(GLU.ALL, ID == 2)
WT.2 <- subset(GLU.ALL, ID == 51)

for (i in 1:nrow(GLU.ALL))
{
  EXPR <- subset(GLU.EXPR, MUTATION == GLU.ALL[i,"MUTATION"] & YFP.CONSTRUCT == GLU.ALL[i,"YFP.CONSTRUCT"])
  GLU.ALL[i,"YFP.MEDIAN"] <- EXPR[1,"YFP.MEDIAN.RELATIVE.MEAN"]
}

WEIGHT <- rep(1,nrow(GLU.ALL))
WEIGHT[which(GLU.ALL$ID ==43)] <- 50
WEIGHT[which(GLU.ALL$ID ==2)] <- 5
GLU.MODEL <- loess(Fitness ~ YFP.MEDIAN, data = GLU.ALL, span=2/3, degree=2, weights = WEIGHT)

#Predict from loess regression
FITNESS.PREDICT <- matrix(0,nrow=nrow(NV.MEAN),ncol=9)
FITNESS.PREDICT[,1] <- predict(GLU.MODEL,NV.MEAN$EFFECT,se=TRUE)$fit
FITNESS.PREDICT[,2] <- predict(GLU.MODEL,NV.MEAN$EFFECT,se=TRUE)$fit + predict(GLU.MODEL,NV.MEAN$EFFECT,se=TRUE)$se
FITNESS.PREDICT[,3] <- predict(GLU.MODEL,NV.MEAN$EFFECT,se=TRUE)$fit - predict(GLU.MODEL,NV.MEAN$EFFECT,se=TRUE)$se

FITNESS.PREDICT[,4] <- predict(GLU.MODEL,NV.MEAN$EFFECT+NV.MEAN.SE$SE,se=TRUE)$fit
FITNESS.PREDICT[,5] <- predict(GLU.MODEL,NV.MEAN$EFFECT+NV.MEAN.SE$SE,se=TRUE)$fit + predict(GLU.MODEL,NV.MEAN$EFFECT+NV.MEAN.SE$SE,se=TRUE)$se
FITNESS.PREDICT[,6] <- predict(GLU.MODEL,NV.MEAN$EFFECT+NV.MEAN.SE$SE,se=TRUE)$fit - predict(GLU.MODEL,NV.MEAN$EFFECT+NV.MEAN.SE$SE,se=TRUE)$se

FITNESS.PREDICT[,7] <- predict(GLU.MODEL,NV.MEAN$EFFECT-NV.MEAN.SE$SE,se=TRUE)$fit
FITNESS.PREDICT[,8] <- predict(GLU.MODEL,NV.MEAN$EFFECT-NV.MEAN.SE$SE,se=TRUE)$fit + predict(GLU.MODEL,NV.MEAN$EFFECT-NV.MEAN.SE$SE,se=TRUE)$se
FITNESS.PREDICT[,9] <- predict(GLU.MODEL,NV.MEAN$EFFECT-NV.MEAN.SE$SE,se=TRUE)$fit - predict(GLU.MODEL,NV.MEAN$EFFECT-NV.MEAN.SE$SE,se=TRUE)$se

par(mar=c(5,5,5,5))
plotCI(log2(NV.MEAN$EFFECT),FITNESS.PREDICT[,1]-1,ui=apply(FITNESS.PREDICT,1,max)-1,li=apply(FITNESS.PREDICT,1,min)-1,pch=19,cex=0.6,xlim=c(-3,1),ylim=c(-.06,0.002),xlab="Expression",ylab="Predicted Fitness Effect of TDH3 expression change")
points(log2(seq(0.1,4,0.001)), predict(GLU.MODEL,seq(0.1,4,0.001))-1,type="l")

abline(h=0)
abline(h=-0.001)
abline(h=0.001)
abline(h=predict(GLU.MODEL,0)- 1,col="red")

sum(apply(FITNESS.PREDICT,1,max) > 1 & apply(FITNESS.PREDICT,1,min) < 1)
sum(apply(FITNESS.PREDICT,1,min) > 1)
sum(apply(FITNESS.PREDICT,1,max) < .999)

###################################
# 11 - TDH3 promoter polymorphism #
###################################
HAPLOTYPES <- read.table("Strain.pTDH3.Haplotypes.txt", header=TRUE)
CIS.DATA <- read.table("FILTER.CIS.DATA.txt",header=TRUE)

CIS.MUTATION <- droplevels(subset(CIS.DATA,CIS.DATA$CLASS == "CIS" & CIS.DATA$TAIL == "ALL" & CIS.DATA$TYPE == "RANDOM"))
CIS.MUTATION.STRAIN <- data.frame(matrix(0,nrow=length(levels(CIS.MUTATION$STRAIN)),ncol=2))
colnames(CIS.MUTATION.STRAIN) <- c("MEAN","NOISE")
rownames(CIS.MUTATION.STRAIN) <- levels(CIS.MUTATION$STRAIN)
CIS.MUTATION.STRAIN$MEAN      <- tapply(CIS.MUTATION$YFP.MEDIAN.RELATIVE,CIS.MUTATION$STRAIN,FUN=mean)
CIS.MUTATION.STRAIN$MEAN.MAD  <- tapply(CIS.MUTATION$YFP.MEDIAN.RELATIVE,CIS.MUTATION$STRAIN,FUN=std.error)
CIS.MUTATION.STRAIN$NOISE     <- tapply(CIS.MUTATION$YFP.CV.RELATIVE,CIS.MUTATION$STRAIN,FUN=mean)
CIS.MUTATION.STRAIN$NOISE.MAD <- tapply(CIS.MUTATION$YFP.CV.RELATIVE,CIS.MUTATION$STRAIN,FUN=std.error)

CIS.NATURAL  <- droplevels(subset(CIS.DATA, CIS.DATA$STRAIN %in% c("A","B","C","E","F","G","H","I","J","K","L","M","N","O","P","Q","R","S","T","U","V","W","Y","Z","AA","DD","EE","FF","GG","HH","II","JJ")))
CIS.NATURAL.STRAIN <- data.frame(matrix(0,nrow=length(levels(CIS.NATURAL$STRAIN)),ncol=2))
colnames(CIS.NATURAL.STRAIN) <- c("MEAN","NOISE")
rownames(CIS.NATURAL.STRAIN) <- levels(CIS.NATURAL$STRAIN)
CIS.NATURAL.STRAIN$MEAN      <- tapply(CIS.NATURAL$YFP.MEDIAN.RELATIVE,CIS.NATURAL$STRAIN,FUN=median)
CIS.NATURAL.STRAIN$MEAN.MAD  <- tapply(CIS.NATURAL$YFP.MEDIAN.RELATIVE,CIS.NATURAL$STRAIN,FUN=mad)
CIS.NATURAL.STRAIN$NOISE     <- tapply(CIS.NATURAL$YFP.CV.RELATIVE,CIS.NATURAL$STRAIN,FUN=median)
CIS.NATURAL.STRAIN$NOISE.MAD <- tapply(CIS.NATURAL$YFP.CV.RELATIVE,CIS.NATURAL$STRAIN,FUN=mad)

var(CIS.NATURAL.STRAIN$MEAN)
var(CIS.MUTATION.STRAIN$MEAN)
var(NV.MEAN$EFFECT)/var(CIS.NATURAL.STRAIN$MEAN)

###########################
# 12 - BM and OU Modeling #
###########################
###Fit models following Bedford and Hartl 2009 PNAS
#Remove S288c as zero branch length from BY reference
DROP.STRAIN <- "S288c"
DROP.NUMBER <- which(PHYLOGENY.PRUNE$tip.label == DROP.STRAIN)
PHYLOGENY.PRUNE <- drop.tip(PHYLOGENY.PRUNE,DROP.STRAIN)

#Get phenotype data in same order as tip labels
EXPR.MEAN <- as.vector(NV.MEAN[PHYLOGENY.PRUNE$tip.label,]$EFFECT)
names(EXPR.MEAN) <- as.vector(NV.MEAN[PHYLOGENY.PRUNE$tip.label,]$NAME)
EXPR.MEAN <- log2(EXPR.MEAN)

EXPR.NOISE <- as.vector(NV.NOISE[PHYLOGENY.PRUNE$tip.label,]$EFFECT)
names(EXPR.NOISE) <- as.vector(NV.NOISE[PHYLOGENY.PRUNE$tip.label,]$NAME)
EXPR.NOISE <- log2(EXPR.NOISE)

#Plot unrooted phylogeny
plot(PHYLOGENY.PRUNE,type="unrooted",cex=0.6)

#Define Reference strain
REF.STRAIN <- "BY4716"
REF.NUMBER <- which(PHYLOGENY.PRUNE$tip.label == REF.STRAIN)

##Calculate distance from reference strain to each pair of strains
I <- 1:length(PHYLOGENY.PRUNE$tip.label)
J <- 1:length(PHYLOGENY.PRUNE$tip.label)
tdmatrix <- matrix(0,nrow=length(I),ncol=length(J))
rownames(tdmatrix) <- PHYLOGENY.PRUNE$tip.label
colnames(tdmatrix) <- PHYLOGENY.PRUNE$tip.label
for(i in I){
  for(j in J){
    STRAIN.KEEP <- c(PHYLOGENY.PRUNE$tip.label[i],PHYLOGENY.PRUNE$tip.label[j],REF.STRAIN)
    TREE <-  drop.tip(PHYLOGENY.PRUNE,PHYLOGENY.PRUNE$tip.label[-match(STRAIN.KEEP,PHYLOGENY.PRUNE$tip.label)])
    tdmatrix[i,j] <- sum(TREE$edge.length)
  }
}

#Remove Reference strain
tdmatrix <- tdmatrix[-REF.NUMBER,-REF.NUMBER]

#Calculate total distance between strains relative to reference strain
meanslist <- diag(tdmatrix)
I <- 1:length(meanslist)
J <- 1:length(meanslist)
cmatrix <- matrix(0,nrow=length(I),ncol=length(J))
rownames(cmatrix) <- rownames(tdmatrix)
colnames(cmatrix) <- colnames(tdmatrix)
for(i in I){
  for(j in J){
    cmatrix[i,j] <- meanslist[i] + meanslist[j]
  }
}

#Calculate shared distance between strains relative to reference strain
sdmatrix <- cmatrix - tdmatrix

#Set up functions for estimating probability densities
#OU Model
OU.pdf <- function(param,data) {
  opt <- param[1]
  drift <- param[2] 
  sel <- param[3]
  if ( sel>0 & drift>0 ) { 
    -sum(dnorm(data[REF.NUMBER],mean=opt,sd=drift/sqrt(2*sel),log=TRUE), dmvnorm(data[-REF.NUMBER],opt*(1-exp(-meanslist*sel)) + data[REF.NUMBER]*exp(-meanslist*sel),(drift^2/(2*sel)) * exp(-sel*cmatrix) * (exp(2*sel*sdmatrix) - 1),log=TRUE))
  } else { 
    10000 
  } 
}

#BM model
BM.pdf <- function(param,data) {
  opt <- param[1]
  drift <- param[2]
  sel <- (drift^2)/2 
  if ( sel>0 & drift>0 ) {
    -sum(dnorm(data[REF.NUMBER],mean=opt,sd=drift/sqrt(2*sel),log=TRUE), dmvnorm(data[-REF.NUMBER],opt*(1-exp(-meanslist*sel)) + data[REF.NUMBER]*exp(-meanslist*sel),(drift^2/(2*sel)) * exp(-sel*cmatrix) * (exp(2*sel*sdmatrix) - 1),log=TRUE))
  } else { 
    10000 
  } 
}

#Fit models by ML for mean expression
OU.MEAN <- optim(c(0,1,1),OU.pdf,gr=NULL,EXPR.MEAN,control=list(maxit=1000))
names(OU.MEAN$par) <- c("opt","drift","sel")
BM.MEAN <- optim(c(0,1),BM.pdf,gr=NULL,EXPR.MEAN,control=list(maxit=1000))
names(BM.MEAN$par) <- c("opt","drift")

#Calculate p-value
#May want to change to a sampling approach to get a more realistic value
1-pchisq(2*(BM.MEAN$value-OU.MEAN$value),1)

#Estimate 95% CIs on parameter estimates
OU.MEAN.HES <- hessian(func=OU.pdf,x=OU.MEAN$par,data=EXPR.MEAN)
OU.MEAN.FISHER <- solve(OU.MEAN.HES)
OU.MEAN.SIGMA <- sqrt(diag(OU.MEAN.FISHER))
OU.MEAN.UPPER <- OU.MEAN$par+1.96*OU.MEAN.SIGMA
OU.MEAN.LOWER <- OU.MEAN$par-1.96*OU.MEAN.SIGMA
OU.MEAN.INTERVAL <-data.frame(value=OU.MEAN$par, upper=OU.MEAN.UPPER, lower=OU.MEAN.LOWER)

BM.MEAN.HES <- hessian(func=BM.pdf,x=BM.MEAN$par,data=EXPR.MEAN)
BM.MEAN.FISHER <- solve(BM.MEAN.HES)
BM.MEAN.SIGMA <- sqrt(diag(BM.MEAN.FISHER))
BM.MEAN.UPPER <- BM.MEAN$par+1.96*BM.MEAN.SIGMA
BM.MEAN.LOWER <- BM.MEAN$par-1.96*BM.MEAN.SIGMA
BM.MEAN.INTERVAL <-data.frame(value=BM.MEAN$par, upper=BM.MEAN.UPPER, lower=BM.MEAN.LOWER)

pal <- colorRampPalette(c("#2D3184","#75CEBE","#F3F1E4"))

OU.MEAN.GRID <- expand.grid(OU.MEAN$par[1],seq(7,20,length=25),seq(.001,6100,length=50))
OU.MEAN.OUT <- unname(cbind(OU.MEAN.GRID,apply(OU.MEAN.GRID ,1,OU.pdf,data=EXPR.MEAN)))
colnames(OU.MEAN.OUT) <- c("opt","drift","sel","pd")
wireframe(log10(pd+6) ~ drift*sel,data=OU.MEAN.OUT,group=opt,drape = TRUE, colorkey = TRUE,scales=list(arrows=FALSE),screen=list(z=0,x=0,y=0), col.regions=pal(100))

BM.MEAN.GRID <- expand.grid(seq(-4,4,length=25),seq(6,11,length=25))
BM.MEAN.OUT <- unname(cbind(BM.MEAN.GRID,apply(BM.MEAN.GRID ,1,BM.pdf,data=EXPR.MEAN)))
colnames(BM.MEAN.OUT) <- c("opt","drift","pd")
wireframe(pd ~ drift*opt,data=BM.MEAN.OUT,drape = TRUE, colorkey = TRUE,scales=list(arrows=FALSE),screen=list(z=0,x=0,y=0),col.regions=pal(100))
```
