## Supplementary material for "Complex polymorphic genetic architecture underlies *trans*-regulatory variation among strains of *Saccharomyces cerevisiae*": Data S1: Analysis.3-Mapping.R

```
########################################
#  R script for analyzing mapping data #
########################################

# 1 - Packages
# 2 - Functions 
# 3 - File Setup
# 4 - SNP Filtering and G' Calculation
# 5 - QTL Identification 
# 6 - Peak Calls
# 7 - Merge Peaks
# 8 - Peak overlaps between strains 
# 9 - Effect Size Estimation 


################
# 1 - Packages #
################
library(VariantAnnotation)
library(Deducer)
library(gtools)
library(zoo)

#################
# 2 - FUNCTIONS #
#################
#Set source directory
source.dir <- "/Path.to.functions"
setwd(source.dir)

source("Functions.R")

##################
# 3 - FILE SETUP #
##################
setwd("/Users/Chimera/Desktop/Trans Natural Variant Paper/Mapping Analysis/April 2018/")

#Set up files and names
SAMPLES <- read.table("SAMPLES.txt",header=TRUE)
VCF.FILES <- list.files(".",pattern=".vcf")
VCF.FILES <- VCF.FILES[c(3:20,1:2)]
  
N <- 1:(length(VCF.FILES))
UB <- numeric(length(N))
LB <- numeric(length(N))
CHROMOSOMES <- read.table("S288c.length",header=TRUE)
GENES <- read.table("GeneListAnalysis.txt",header=TRUE)
CANDIDATE <- read.table("Candidate.Genes.txt",header=TRUE)
OUTPUT <- list()
DATA.G <- list()
G.PRIME <- list()
DATA.G.PRIME <- list()
SEG <- list()
FIXED <- list()
FRAGMENT.LOW <- list()
FRAGMENT.HIGH <- list()
STATS <- list()

CHROMOSOME.LABELS <- c("I","II","III","IV","V","VI","VII","VIII","IX","X","XI","XII","XIII","XIV","XV","XVI")

########################################
# 4 - SNP Filtering and G' Calculation #
########################################

for(n in N) {
  paste(SAMPLES[n,1],sep="")
  
  ###SNP Filtering###
  #Get SNP Filtering function
  source("SNP.FILTER.R")

  #Set SNP Filtering options
  MAP.FILTER <- 50
  QUAL.FILTER <- 30
  PAIR.FILTER <- 0
  DEPTH.MIN <- 20
  DEPTH.MAX <- 500
  FREQ.CUTOFF <- 0.2
  
  #Apply SNP Filtering  
  OUTPUT[[n]] <- SNP.FILTER(readVcf(VCF.FILES[n],"S288c"), MAP.FILTER,QUAL.FILTER,PAIR.FILTER,DEPTH.MIN,DEPTH.MAX)

  #Remove low quality SNPs
  DATA <- OUTPUT[[n]][[1]]
  GOOD <- which(DATA$PASS == 1)
  GOOD.DATA <- DATA[GOOD,]

  #Calculate allele frequencies
  FREQ.LOW  <- GOOD.DATA$AO.L/(GOOD.DATA$AO.L + GOOD.DATA$RO.L)
  FREQ.HIGH <- GOOD.DATA$AO.H/(GOOD.DATA$AO.H + GOOD.DATA$RO.H)

  #Calculate difference in allele frequncy between neighboring SNPs
  FREQ.DIFF.LOW.INCREASE <- c(diff(FREQ.LOW,1),0)
  FREQ.DIFF.LOW.DECREASE <- c(0,rev(diff(rev(FREQ.LOW),1)))
  FREQ.DIFF.HIGH.INCREASE <- c(diff(FREQ.HIGH,1),0)
  FREQ.DIFF.HIGH.DECREASE <- c(0,rev(diff(rev(FREQ.HIGH),1)))

  #Remove SNPs with large frequency differences from both neighboring SNPs
  KEEP <- which((abs(FREQ.DIFF.LOW.INCREASE)  < FREQ.CUTOFF | abs(FREQ.DIFF.LOW.DECREASE)  < FREQ.CUTOFF | sign(FREQ.DIFF.LOW.DECREASE*FREQ.DIFF.LOW.INCREASE)   != 1) & 
	              (abs(FREQ.DIFF.HIGH.INCREASE) < FREQ.CUTOFF | abs(FREQ.DIFF.HIGH.DECREASE) < FREQ.CUTOFF | sign(FREQ.DIFF.HIGH.DECREASE*FREQ.DIFF.HIGH.INCREASE) != 1) &
		             nchar(GOOD.DATA$REF) == 1)

  OUTPUT[[n]][[1]][GOOD,][-KEEP,]$PASS <- 0

  DATA.G[[n]] <- subset(OUTPUT[[n]][[1]],OUTPUT[[n]][[1]]$PASS == 1)
  SEG[[n]]    <- subset(DATA.G[[n]], DATA.G[[n]]$RO.L + DATA.G[[n]]$RO.H > 5)
  FIXED[[n]]  <- subset(DATA.G[[n]], DATA.G[[n]]$RO.L + DATA.G[[n]]$RO.H < 5)

  ###G' calculation
  #Ignore SNPs on mitochondria or in drug resistance markers
  I <- 1:nrow(subset(DATA.G[[n]],!DATA.G[[n]]$CHR %in% c("chrMito","chrNat","chrKan")))
  G.PRIME[[n]] <- data.frame(matrix(0,nrow=length(I),ncol=2))
  colnames(G.PRIME[[n]]) <- c("ABS","DIR")
  
  #Set window for G' calculation
  W <- 20000

  for(i in I) {
	  #Get SNPs within window
	  WINDOW <- which(DATA.G[[n]]$CHR == DATA.G[[n]]$CHR[i] & DATA.G[[n]]$POS < DATA.G[[n]]$POS[i] + W & DATA.G[[n]]$POS > DATA.G[[n]]$POS[i] - W)
	  SUBSET <- DATA.G[[n]][WINDOW,]

	  #Calculate weights
	  if(nrow(SUBSET) == 1) {
		  D <- 0
	  } else {
	  	D <- abs(DATA.G[[n]]$POS[i] - SUBSET$POS)/(max(abs(DATA.G[[n]]$POS[i] - SUBSET$POS)))
	  }
	  S <- sum((1-D^3)^3)
	  K <- ((1-D^3)^3)/S

	  G.PRIME[[n]][i,1] <- sum(abs(DATA.G[[n]]$G[WINDOW])*K)
	  G.PRIME[[n]][i,2] <- sum(DATA.G[[n]]$G[WINDOW]*K)
  }
  DATA.G.PRIME[[n]] <- cbind(subset(DATA.G[[n]],!DATA.G[[n]]$CHR %in% c("chrMito","chrNat","chrKan")),G.PRIME[[n]])
  colnames(DATA.G.PRIME[[n]]) <- c(colnames(DATA.G[[n]]),colnames(G.PRIME[[n]]))

  #Set chromosome boundries to zero
  I <- 2:33
  BOUNDARY <- data.frame(matrix(0,nrow=length(I),ncol=4))
  names(BOUNDARY) <- c("CHR","POS",colnames(G.PRIME[[n]]))
  BOUNDARY[,1] <- 1
  for(i in I) {
	  BOUNDARY[i-1,1] <- paste("chr",sprintf("%02d",i%/%2),sep="")
	  if(i%%2 == 0) {
		  BOUNDARY[i-1,2] <- 1
	  }else {
		  BOUNDARY[i-1,2] <- CHROMOSOMES$LENGTH[i%/%2]
	  }
	  BOUNDARY[i-1,3] <- 0
	  BOUNDARY[i-1,4] <- 0
  }
  DATA.G.PRIME[[n]] <- smartbind(DATA.G.PRIME[[n]],BOUNDARY)
  DATA.G.PRIME[[n]] <- DATA.G.PRIME[[n]][order(DATA.G.PRIME[[n]]$CHR,DATA.G.PRIME[[n]]$POS),]
  rownames(DATA.G.PRIME[[n]]) <- 1:nrow(DATA.G.PRIME[[n]])
}

#Write G' to file
for(n in N) {
	write.table(DATA.G.PRIME[[n]],paste(SAMPLES[n,1],".data.txt",sep=""),quote=FALSE,sep="\t")
}

#Read G' 
for(n in N) {
  DATA.G.PRIME[[n]] <- read.table(paste(SAMPLES[n,1],".data.txt",sep=""),header=TRUE)
}


##########################
# 5 - QTL Identification #
##########################

#G' and SNP Frequency Plots
for(n in N) {
  pdf(paste(SAMPLES[n,1],".qtl.pdf",sep=""))

  I <- 1:16
  
  #G' across the genome
  par(mfrow=c(1,1))
  par(mar=c(5.1,4.1,4.1,2.1))
  plot(0,xlim=c(0,12080000),ylim=c(-1*max(abs(DATA.G.PRIME[[n]]$DIR)),max(abs(DATA.G.PRIME[[n]]$DIR))),type="n",xaxt="n",xlab="Position (Mb)",ylab="G'",main="")
  for(i in I) {
	  x <- subset(DATA.G.PRIME[[n]],DATA.G.PRIME[[n]]$CHR==paste("chr",sprintf("%02d", i),sep=""))
	  points(x$POS+sum(CHROMOSOMES[0:(i-1),]$LENGTH), x$DIR, type="l",col="black")
	  abline(v=sum(CHROMOSOMES[0:(i-1),]$LENGTH),lty=2,col="gray")
  }
  abline(v = sum(CHROMOSOMES$LENGTH),lty=2,col="gray")
  abline(h = 0)
  abline(h = 5, col="red")
  abline(h = -5, col="red")
  axis(1,at=c(seq(0,12080000,1000000)),labels=seq(0,12,1))
  axis(3,at=(2*cumsum(CHROMOSOMES$LENGTH)-CHROMOSOMES$LENGTH)/2,CHROMOSOME.LABELS,cex.axis=0.6)

  #Individual chromosome G'
  par(mfrow=c(4,4))
  par(mar=c(2,2,2,2))
  for(i in I) {
	  x <- subset(DATA.G.PRIME[[n]],DATA.G.PRIME[[n]]$CHR==paste("chr",sprintf("%02d",i),sep=""))
	  plot(x$POS/1000, x$DIR, type="l",col="black",ylab="",xlab="",main=CHROMOSOME.LABELS[i],lwd=2,ylim=c(-1*max(x$ABS),max(x$ABS)),cex.axis=0.8)
  	abline(h = 0)
	  abline(h = 5, col="red")
	  abline(h = -5, col="red")
  }

  #Individual chromosome SNP Frequency
  COLOR <- numeric(2)
  if(n %in% c(1,4,7,10,13,16)) {
	  COLOR <- colors()[c(24,79)]
  } else if(n %in% c(2,5,8,11,14,17)) {
	  COLOR <- colors()[c(30,122)]
  } else if(n %in% c(3,6,9,12,15,18)) {
	  COLOR <- colors()[c(81,514)]
  } else if (n %in% c(19,20)) {
    COLOR <- colors()[c(34,556)]
  }
  par(mfrow=c(4,4))
  par(mar=c(2,2,2,2))
  for(i in I) {
  	x <- subset(DATA.G.PRIME[[n]],DATA.G.PRIME[[n]]$CHR==paste("chr",sprintf("%02d",i),sep=""))
	  plot(x$POS/1000, x$AO.L/(x$AO.L + x$RO.L), pch=19,cex=0.1,ylim=c(0,1),col=COLOR[2],xlab="Position",ylab="Frequency",main=CHROMOSOME.LABELS[i],cex.axis=0.8)
	  points(x$POS/1000, x$AO.H/(x$AO.H + x$RO.H), pch=19,cex=0.1,col=COLOR[1])
	  abline(h=0.5)
  }

  #Individual chromosome SNP Frequency Differences
  par(mfrow=c(4,4))
  par(mar=c(2,2,2,2))
  for(i in I) {
	  x <- subset(DATA.G.PRIME[[n]],DATA.G.PRIME[[n]]$CHR==paste("chr",sprintf("%02d",i),sep=""))
	  plot(x$POS/1000,x$AO.H/(x$AO.H + x$RO.H) -  x$AO.L/(x$AO.L + x$RO.L), pch=19,cex=0.1,ylim=c(-1.2,1),col="black",xlab="Position",ylab="Frequency",main=CHROMOSOME.LABELS[i],cex.axis=0.8)
	  abline(h=0)
  }
dev.off()
}


##################
# 6 - Peak calls #
##################

for(n in N) {
  pdf(paste(SAMPLES[n,1],".qtl2.pdf",sep=""))

  W <- 50000
  CUTOFF.1 <- 5
  CUTOFF.2 <- 5

  MAX.OUT <- list()
  MIN.OUT <- list()
  MAX.INT <- list()
  length(MAX.INT) <- 16
  MIN.INT <- list()
  length(MIN.INT) <- 16

  J <- 1:16
  for(j in J) {
	  CHR.DATA <- subset(DATA.G.PRIME[[n]], DATA.G.PRIME[[n]]$CHR == paste("chr",sprintf("%02d", j),sep="")) 
	  ADD <- min(as.numeric(row.names(CHR.DATA)))

	  I <- 1:nrow(CHR.DATA)
	  MAX.OUT[[j]] <- numeric(length(I))
	  MIN.OUT[[j]] <- numeric(length(I))
	  for(i in I) {
		  SITE <- CHR.DATA[i,]
		  SET <- subset(CHR.DATA,abs(CHR.DATA$POS - SITE$POS) < W)

		  MAX <- as.numeric(rownames(SET[which(SET$DIR == max(SET$DIR)),]))
		  MIN <- as.numeric(rownames(SET[which(SET$DIR == min(SET$DIR)),]))

		  while(as.numeric(rownames(SITE)) != MAX) {
			  SITE <- DATA.G.PRIME[[n]][MAX,]
			  SET <- subset(CHR.DATA,abs(CHR.DATA$POS - SITE$POS) < W)
			  MAX <- as.numeric(rownames(SET[which(SET$DIR == max(SET$DIR)),]))
		  }
		  MAX.OUT[[j]][i] <- MAX

		  while(as.numeric(rownames(SITE)) != MIN) {
			  SITE <- DATA.G.PRIME[[n]][MIN,]
			  SET <- subset(CHR.DATA,abs(CHR.DATA$POS - SITE$POS) < W)
			  MIN <- as.numeric(rownames(SET[which(SET$DIR == min(SET$DIR)),]))
		  }
		  MIN.OUT[[j]][i] <- MIN
  	}
	  FILTER.MAX.OUT <- unique(subset(MAX.OUT[[j]],CHR.DATA$DIR > CUTOFF.1))
	  FILTER.MIN.OUT <- unique(subset(MIN.OUT[[j]],CHR.DATA$DIR < -CUTOFF.1))

	  #Plot G'
	  par(mfrow=c(1,1))
	  par(mar=c(5.1,4.1,4.1,2.1))
	  plot(0,xlim=c(0,max(CHR.DATA$POS)),ylim=c(-1*max(abs(DATA.G.PRIME[[n]]$DIR)),max(abs(DATA.G.PRIME[[n]]$DIR))),type="n",xaxt="n",xlab="Position",ylab="G",main="")
	  abline(v = seq(0,max(CHR.DATA$POS),10000),col="lightgray",lty=2)	
	  points(CHR.DATA$POS, CHR.DATA$DIR, pch=19,cex=0.2,col="black")
	  abline(h = 0)
	  abline(h = 5, col="red")
	  abline(h = -5, col="red")
	
	  #Add peaks and CIs for increases
	  if(length(FILTER.MAX.OUT) > 0) {
	  	T <- 1:length(FILTER.MAX.OUT)
		  MAX.INT[[j]] <- matrix(0,nrow=length(T),ncol=3)
		  for(t in T) {
			  MAX.INT[[j]][t,1] <- FILTER.MAX.OUT[t]
			  TEST <- CHR.DATA$DIR < DATA.G.PRIME[[n]][FILTER.MAX.OUT[t],]$DIR - CUTOFF.2 | CHR.DATA$DIR > DATA.G.PRIME[[n]][FILTER.MAX.OUT[t],]$DIR
		  	M <- (FILTER.MAX.OUT - ADD)[t]+1
			  TEST.L <- rev(TEST[1:(M-1)])
			  TEST.H <- TEST[(M+1):length(TEST)]
			  MAX.INT[[j]][t,2] <-  M - min(which(TEST.L == TRUE)) + ADD - 1
			  MAX.INT[[j]][t,3] <-  M + min(which(TEST.H == TRUE)) + ADD - 1

		  	L.BOUND <- DATA.G.PRIME[[n]][MAX.INT[[j]][t,2]:MAX.INT[[j]][t,1],]$DIR
			  H.BOUND <- DATA.G.PRIME[[n]][MAX.INT[[j]][t,1]:MAX.INT[[j]][t,3],]$DIR
		  	MAX.INT[[j]][t,2] <- which(L.BOUND == min(L.BOUND)) - 1 + MAX.INT[[j]][t,2]
			  MAX.INT[[j]][t,3] <- which(H.BOUND == min(H.BOUND)) - 1 + MAX.INT[[j]][t,1]
		  }
		  abline(v= DATA.G.PRIME[[n]][MAX.INT[[j]][,1],"POS"],col="red")
		  abline(v= DATA.G.PRIME[[n]][MAX.INT[[j]][,2],"POS"],col="red",lty=2)
		  abline(v= DATA.G.PRIME[[n]][MAX.INT[[j]][,3],"POS"],col="red",lty=2)
	  }

	  #Add peaks and CIs for decreases
  	if(length(FILTER.MIN.OUT) > 0) {
	  	T <- 1:length(FILTER.MIN.OUT)
		  MIN.INT[[j]] <- matrix(0,nrow=length(T),ncol=3)
		  for(t in T) {
			  MIN.INT[[j]][t,1] <- FILTER.MIN.OUT[t]
			  TEST <- CHR.DATA$DIR > DATA.G.PRIME[[n]][FILTER.MIN.OUT[t],]$DIR + CUTOFF.2 | CHR.DATA$DIR < DATA.G.PRIME[[n]][FILTER.MIN.OUT[t],]$DIR
			  M <- (FILTER.MIN.OUT - ADD)[t]+1
			  TEST.L <- rev(TEST[1:(M-1)])
			  TEST.H <- TEST[(M+1):length(TEST)]
			  MIN.INT[[j]][t,2] <-  M - min(which(TEST.L == TRUE)) + ADD - 1
			  MIN.INT[[j]][t,3] <-  M + min(which(TEST.H == TRUE)) + ADD - 1

			  L.BOUND <- DATA.G.PRIME[[n]][MIN.INT[[j]][t,2]:MIN.INT[[j]][t,1],]$DIR
			  H.BOUND <- DATA.G.PRIME[[n]][MIN.INT[[j]][t,1]:MIN.INT[[j]][t,3],]$DIR
			  MIN.INT[[j]][t,2] <- which(L.BOUND == max(L.BOUND)) - 1 + MIN.INT[[j]][t,2]
			  MIN.INT[[j]][t,3] <- which(H.BOUND == max(H.BOUND)) - 1 + MIN.INT[[j]][t,1]
		  }
		  abline(v= DATA.G.PRIME[[n]][MIN.INT[[j]][,1],"POS"],col="blue")
		  abline(v= DATA.G.PRIME[[n]][MIN.INT[[j]][,2],"POS"],col="blue",lty=2)
		  abline(v= DATA.G.PRIME[[n]][MIN.INT[[j]][,3],"POS"],col="blue",lty=2)
	  }
  }

  #Output info for increase peaks
  if(length(do.call(rbind,MAX.INT)) > 0) {
    PEAK.MAX.CALLS <- data.frame(cbind(rep(1:16,sapply(MAX.INT,FUN=length)/3), 
                                       do.call(rbind,MAX.INT),
                                       do.call(rbind,sapply(MAX.INT,FUN=function(x) {matrix(DATA.G.PRIME[[n]][x,"POS"],ncol=3,byrow=F)})),
                                       rep(1,sum(sapply(MAX.INT,FUN=length)/3)),
                                       do.call(rbind,sapply(MAX.INT,FUN=function(x) {matrix(DATA.G.PRIME[[n]][x[,1],"DIR"])}))))
  }
  #Output info for decrease peaks
  if(length(do.call(rbind,MIN.INT)) > 0) {
	  PEAK.MIN.CALLS <- data.frame(cbind(rep(1:16,sapply(MIN.INT,FUN=length)/3), 
	                                  do.call(rbind,MIN.INT),
	                                  do.call(rbind,sapply(MIN.INT,FUN=function(x) {matrix(DATA.G.PRIME[[n]][x,"POS"],ncol=3,byrow=F)})),
	                                  rep(-1,sum(sapply(MIN.INT,FUN=length)/3)),
	                                  do.call(rbind,sapply(MIN.INT,FUN=function(x) {matrix(DATA.G.PRIME[[n]][x[,1],"DIR"])}))))
  }
  #Merge and write to file
  PEAK.CALLS <- rbind(PEAK.MAX.CALLS,PEAK.MIN.CALLS)
  colnames(PEAK.CALLS) <- c("CHR","SNP.PEAK","SNP.MIN","SNP.MAX","POS.PEAK","POS.MIN","POS.MAX","DIR","EFFECT")
  write.table(PEAK.CALLS,paste(SAMPLES[n,1],".peak.calls.txt",sep=""),quote=FALSE,sep="\t",row.names=FALSE)
  dev.off()
}

###################
# 7 - Merge Peaks #
###################
#Strain, cross, and selection IDs
LAYOUT.MAPPING <- read.table("Layout.Mapping.txt",header=TRUE)

#Read in peak locations
PEAK.CALL.FILES <- list.files(".",pattern="peak.calls.5.txt")
PEAK.CALL.FILES <- PEAK.CALL.FILES[c(3:20,1:2)]

I <- 1:length(PEAK.CALL.FILES)
PEAK.CALLS <- list()
for(i in I) {
  PEAK.CALLS[[i]] <- read.table(PEAK.CALL.FILES[i],header=TRUE)
  SAMPLE.STRAIN <- rep(LAYOUT.MAPPING$SAMPLE.STRAIN[i],nrow(PEAK.CALLS[[i]]))
  SAMPLE.CROSS  <- rep(LAYOUT.MAPPING$CROSS[i],        nrow(PEAK.CALLS[[i]]))
  SAMPLE.SELECT <- rep(LAYOUT.MAPPING$SELECT[i],       nrow(PEAK.CALLS[[i]]))
  PEAK.CALLS[[i]] <- cbind(PEAK.CALLS[[i]],SAMPLE.STRAIN,SAMPLE.CROSS,SAMPLE.SELECT)
}

##Merge Peaks within each sample
PEAK.MERGE <- list()
for(n in N) {
  #Collect overlaps
  I <- 1:nrow(PEAK.CALLS[[n]])
  PEAK.MATRIX <- matrix(0,nrow=length(I),ncol=length(I))
  for(i in I) {
    #For merging, a peak overlaps a second peak if:
    #1) It is on the same chromosome 
    #2) In the same direction
    #3) The second peak, or one of its CIs, are within the CI of the first peak
    OUT <- which(   PEAK.CALLS[[n]]$CHR == PEAK.CALLS[[n]]$CHR[i] &
                    PEAK.CALLS[[n]]$DIR == PEAK.CALLS[[n]]$DIR[i] &   
                   (PEAK.CALLS[[n]]$POS.MIN  > PEAK.CALLS[[n]]$POS.MIN[i] & PEAK.CALLS[[n]]$POS.MIN  < PEAK.CALLS[[n]]$POS.MAX[i] |
                    PEAK.CALLS[[n]]$POS.PEAK > PEAK.CALLS[[n]]$POS.MIN[i] & PEAK.CALLS[[n]]$POS.PEAK < PEAK.CALLS[[n]]$POS.MAX[i] |
                    PEAK.CALLS[[n]]$POS.MAX  > PEAK.CALLS[[n]]$POS.MIN[i] & PEAK.CALLS[[n]]$POS.MAX  < PEAK.CALLS[[n]]$POS.MAX[i] ))
    PEAK.MATRIX[i,OUT] <- 1
  }

  #Merge overlaps
  PEAK.MERGE[[n]] <- PEAK.CALLS[[n]]
  SITE <- 1
  X <- 1
  while(SITE <= nrow(PEAK.MERGE[[n]])) {
    if(SITE < nrow(PEAK.MERGE[[n]])) {
      while(sum(PEAK.MATRIX[SITE:(SITE+X),SITE:(SITE+X)]) >= 0.7*(X+1)^2) {
        if(SITE + X + 1 <= nrow(PEAK.MERGE[[n]])) {
          X <- X + 1
        } else {
          X <- X + 1
         break
        }
      }
    }
    X <- X-1
  
  
    if(X == 0) {
      #If peak doesn't have any overlaps
      PEAK.MERGE[[n]][SITE,] <- PEAK.CALLS[[n]][SITE,]
    } else {
      #If peak has overlaps, weight merged peaks by width and height
      PEAK.MERGE[[n]][SITE:(SITE+X),] <- PEAK.CALLS[[n]][SITE:(SITE+X),]
      PEAK.STARTS <- PEAK.CALLS[[n]]$POS.MIN[SITE:(SITE+X)]
      PEAK.MIDS   <- PEAK.CALLS[[n]]$POS.PEAK[SITE:(SITE+X)]
      PEAK.STOPS  <- PEAK.CALLS[[n]]$POS.MAX[SITE:(SITE+X)]
      PEAK.WIDTH <- PEAK.STOPS - PEAK.STARTS + 1
      WIDTH.WEIGHTS <- 1 - (PEAK.WIDTH^2)/sum(PEAK.WIDTH^2)
      PEAK.HEIGHTS <- PEAK.CALLS[[n]]$EFFECT[SITE:(SITE+X)]
      HEIGHT.WEIGHTS <- (PEAK.HEIGHTS^2)/sum(PEAK.HEIGHTS^2)
    
      PEAK.MERGE[[n]]$POS.PEAK[SITE:(SITE+X)] <- round(sum(0.5*(WIDTH.WEIGHTS*PEAK.MIDS)   + 0.5*(HEIGHT.WEIGHTS*PEAK.MIDS)),0)
      PEAK.MERGE[[n]]$POS.MIN[SITE:(SITE+X)]  <- round(sum(0.5*(WIDTH.WEIGHTS*PEAK.STARTS) + 0.5*(HEIGHT.WEIGHTS*PEAK.STARTS)),0)
      PEAK.MERGE[[n]]$POS.MAX[SITE:(SITE+X)]  <- round(sum(0.5*(WIDTH.WEIGHTS*PEAK.STOPS)  + 0.5*(HEIGHT.WEIGHTS*PEAK.STOPS)),0)
      PEAK.MERGE[[n]]$EFFECT[SITE:(SITE+X)]   <- PEAK.CALLS[[n]]$DIR[SITE]*max(abs(PEAK.HEIGHTS))
    }
    SITE <- SITE + X + 1
    X <- 1
  }
  PEAK.MERGE[[n]] <- PEAK.MERGE[[n]][,c(1,5:10)]
  PEAK.MERGE[[n]] <- unique(PEAK.MERGE[[n]])
}

#####################################
# 8 - Peak overlaps between strains #
#####################################
#Use only 3 cross, 3 selection samples
STRAIN.OVERLAP <- rbind(PEAK.MERGE[[7]],PEAK.MERGE[[8]],PEAK.MERGE[[9]])
STRAIN.OVERLAP <- STRAIN.OVERLAP[order(STRAIN.OVERLAP$CHR,STRAIN.OVERLAP$POS.PEAK),]
STRAIN.OVERLAP$SAMPLE.STRAIN <- factor(STRAIN.OVERLAP$SAMPLE.STRAIN)

STRAIN.MERGE <- OVERLAP(STRAIN.OVERLAP)
STRAIN.UP.MERGE <- OVERLAP(subset(STRAIN.OVERLAP,STRAIN.OVERLAP$DIR == 1))
STRAIN.DOWN.MERGE <- OVERLAP(subset(STRAIN.OVERLAP,STRAIN.OVERLAP$DIR == -1))

#Calculate p-values for overlaps
REPS <- 10000
TEST.OVERLAP <- OVERLAP.TEST(STRAIN.OVERLAP,REPS)
hist(TEST.OVERLAP,breaks=20)
abline(v = (sum(STRAIN.MERGE$DOUBLES) + STRAIN.MERGE$TRIPLES)/(sum(STRAIN.MERGE$SINGLES) + sum(STRAIN.MERGE$DOUBLES) + STRAIN.MERGE$TRIPLES), col= "red")
sum(TEST.OVERLAP > (sum(STRAIN.MERGE$DOUBLES) + STRAIN.MERGE$TRIPLES)/(sum(STRAIN.MERGE$SINGLES) + sum(STRAIN.MERGE$DOUBLES) + STRAIN.MERGE$TRIPLES))/REPS

TEST.OVERLAP.UP <- OVERLAP.TEST(subset(STRAIN.OVERLAP,STRAIN.OVERLAP$DIR == 1),REPS)
hist(TEST.OVERLAP.UP,breaks=20)
abline(v = (sum(STRAIN.UP.MERGE$DOUBLES) + STRAIN.UP.MERGE$TRIPLES)/(sum(STRAIN.UP.MERGE$SINGLES) + sum(STRAIN.UP.MERGE$DOUBLES) + STRAIN.UP.MERGE$TRIPLES), col= "red")
sum(TEST.OVERLAP.UP > (sum(STRAIN.UP.MERGE$DOUBLES) + STRAIN.UP.MERGE$TRIPLES)/(sum(STRAIN.UP.MERGE$SINGLES) + sum(STRAIN.UP.MERGE$DOUBLES) + STRAIN.UP.MERGE$TRIPLES))/REPS

TEST.OVERLAP.DOWN <- OVERLAP.TEST(subset(STRAIN.OVERLAP,STRAIN.OVERLAP$DIR == -1),REPS)
hist(TEST.OVERLAP.DOWN,breaks=20)
abline(v = (sum(STRAIN.DOWN.MERGE$DOUBLES) + STRAIN.DOWN.MERGE$TRIPLES)/(sum(STRAIN.DOWN.MERGE$SINGLES) + sum(STRAIN.DOWN.MERGE$DOUBLES) + STRAIN.DOWN.MERGE$TRIPLES), col= "red")
sum(TEST.OVERLAP.DOWN > (sum(STRAIN.DOWN.MERGE$DOUBLES) + STRAIN.DOWN.MERGE$TRIPLES)/(sum(STRAIN.DOWN.MERGE$SINGLES) + sum(STRAIN.DOWN.MERGE$DOUBLES) + STRAIN.DOWN.MERGE$TRIPLES))/REPS

#Test overlap between M22 and M22 repeat
M22a.M22b.MERGE <- rbind(PEAK.MERGE[[9]],PEAK.MERGE[[20]])
M22a.M22b.MERGE <- M22a.M22b.MERGE[order(M22a.M22b.MERGE$CHR,M22a.M22b.MERGE$POS.PEAK),]
M22a.M22b.MERGE$SAMPLE.STRAIN <- factor(M22a.M22b.MERGE$SAMPLE.STRAIN)
M22a.M22b.OVERLAP <- OVERLAP(M22a.M22b.MERGE)

TEST.M22a.M22b.OVERLAP <- OVERLAP.TEST(M22a.M22b.MERGE, REPS)
hist(TEST.M22a.M22b.OVERLAP,breaks=10,xlim=c(0,1))
abline(v = sum(M22a.M22b.OVERLAP$DOUBLES)/(min(M22a.M22b.OVERLAP$SINGLES[M22a.M22b.OVERLAP$SINGLES > 0]) + sum(M22a.M22b.OVERLAP$DOUBLES)), col= "red")
sum(TEST.M22a.M22b.OVERLAP > sum(M22a.M22b.OVERLAP$DOUBLES)/(min(M22a.M22b.OVERLAP$SINGLES[M22a.M22b.OVERLAP$SINGLES > 0]) + sum(M22a.M22b.OVERLAP$DOUBLES)))/REPS

#Get overlap between M22 replicate and SK1/YPS1000 experiments
SK1.YPS1000.M22b.MERGE <- rbind(PEAK.MERGE[[7]],PEAK.MERGE[[8]],PEAK.MERGE[[20]])
SK1.YPS1000.M22b.MERGE <- SK1.YPS1000.M22b.MERGE[order(SK1.YPS1000.M22b.MERGE$CHR,SK1.YPS1000.M22b.MERGE$POS.PEAK),]
SK1.YPS1000.M22b.MERGE$SAMPLE.STRAIN <- factor(SK1.YPS1000.M22b.MERGE$SAMPLE.STRAIN)
SK1.YPS1000.M22b.OVERLAP <- OVERLAP(SK1.YPS1000.M22b.MERGE)

#Fisher exact test for comparing degree of overlap between experiments
#M22a.M22b vs. YPS1000.M22b and SK1.M22b
m <- matrix(c(54,20,43,31),nrow=2); fisher.test(m)
m <- matrix(c(54,20,40,34),nrow=2); fisher.test(m)

#M22a.M22b vs. YPS1000.M22a and SK1.M22a
m <- matrix(c(54,20,58,53),nrow=2); fisher.test(m)
m <- matrix(c(54,20,62,49),nrow=2); fisher.test(m)

#Frequency of increases and decreases per strain
barplot(c(table(subset(STRAIN.OVERLAP,STRAIN.OVERLAP$DIR == 1)$SAMPLE.STRAIN),-1*table(subset(STRAIN.OVERLAP,STRAIN.OVERLAP$DIR == -1)$SAMPLE.STRAIN)),ylim=c(-60,60))

#Plot G' and merged peaks 
N <- c(7:9,20)
J <- 1:16
pdf("Final.Peak.Calls.pdf")
for(n in N) {
  D <- DATA.G.PRIME[[n]]
  PEAKS <- PEAK.MERGE[[n]]
  COLOR <- list("steelblue3","darkgreen","goldenrod3","darkred")[[which(N == n)]]
  
  #G' across the genome
  par(mfrow=c(1,1))
  par(mar=c(5.1,4.1,4.1,2.1))
  plot(0,xlim=c(0,12080000),ylim=c(-1*max(abs(D$DIR)),max(abs(D$DIR))),type="n",xaxt="n",xlab="Position (Mb)",ylab="G'",main="")
  for(j in J) {
    #Plot G.Prime
    G.DATA <- subset(D,D$CHR==paste("chr",sprintf("%02d", j),sep=""))
    points(G.DATA$POS+sum(CHROMOSOMES[0:(j-1),]$LENGTH), G.DATA$DIR, type="l",col="black")
    abline(v=sum(CHROMOSOMES[0:(j-1),]$LENGTH),lty=2,col="gray")
    
    #Add called eQTL
    PEAK.DATA <- subset(PEAKS,PEAKS$CHR == j)
    PEAK.DATA.UP   <- subset(PEAK.DATA, PEAK.DATA$DIR == 1)
    PEAK.DATA.DOWN <- subset(PEAK.DATA, PEAK.DATA$DIR == -1)
    
    if(nrow(PEAK.DATA.UP) > 0) {
    segments(x0 = PEAK.DATA.UP$POS.PEAK+sum(CHROMOSOMES[0:(j-1),]$LENGTH), y0=max(abs(D$DIR)), x1 = PEAK.DATA.UP$POS.PEAK+sum(CHROMOSOMES[0:(j-1),]$LENGTH), y1 = 0, col=COLOR)
    segments(x0 = PEAK.DATA.UP$POS.MAX +sum(CHROMOSOMES[0:(j-1),]$LENGTH), y0=max(abs(D$DIR)), x1 = PEAK.DATA.UP$POS.MAX+sum(CHROMOSOMES[0:(j-1),]$LENGTH), y1 = 0, col=COLOR, lty=2)
    segments(x0 = PEAK.DATA.UP$POS.MIN +sum(CHROMOSOMES[0:(j-1),]$LENGTH), y0=max(abs(D$DIR)), x1 = PEAK.DATA.UP$POS.MIN+sum(CHROMOSOMES[0:(j-1),]$LENGTH), y1 = 0, col=COLOR, lty=2)
    } 
    if(nrow(PEAK.DATA.DOWN) > 0) {
    segments(x0 = PEAK.DATA.DOWN$POS.PEAK+sum(CHROMOSOMES[0:(j-1),]$LENGTH), y0=-1*max(abs(D$DIR)), x1 = PEAK.DATA.DOWN$POS.PEAK+sum(CHROMOSOMES[0:(j-1),]$LENGTH), y1 = 0, col=COLOR)
    segments(x0 = PEAK.DATA.DOWN$POS.MAX +sum(CHROMOSOMES[0:(j-1),]$LENGTH), y0=-1*max(abs(D$DIR)), x1 = PEAK.DATA.DOWN$POS.MAX+sum(CHROMOSOMES[0:(j-1),]$LENGTH), y1 = 0, col=COLOR, lty=2)
    segments(x0 = PEAK.DATA.DOWN$POS.MIN +sum(CHROMOSOMES[0:(j-1),]$LENGTH), y0=-1*max(abs(D$DIR)), x1 = PEAK.DATA.DOWN$POS.MIN+sum(CHROMOSOMES[0:(j-1),]$LENGTH), y1 = 0, col=COLOR, lty=2)
    }
  }
  abline(v = sum(CHROMOSOMES$LENGTH),lty=2,col="gray")
  abline(h = 0)
  abline(h = 5, col="red")
  abline(h = -5, col="red")
  axis(1,at=c(seq(0,12080000,1000000)),labels=seq(0,12,1))
  axis(3,at=(2*cumsum(CHROMOSOMES$LENGTH)-CHROMOSOMES$LENGTH)/2,CHROMOSOME.LABELS,cex.axis=0.6)

  #Individual Chromosomes
  for(j in J) {
    CHR.DATA <- subset(D, D$CHR == paste("chr",sprintf("%02d", j),sep="")) 
    PEAK.DATA <- subset(PEAKS,PEAKS$CHR == j)
    
    par(mfrow=c(1,1))
    par(mar=c(5.1,4.1,4.1,2.1))
    plot(0,xlim=c(0,max(CHR.DATA$POS)),ylim=c(-1*max(abs(DATA.G.PRIME[[n]]$DIR)),max(abs(DATA.G.PRIME[[n]]$DIR))),type="n",xaxt="n",xlab="Position",ylab="G",main=paste(c("YPS1000","SK1","M22","M22b")[[which(N == n)]]," chr",sprintf("%02d", j),sep=""))
    abline(v = seq(0,max(CHR.DATA$POS),10000),col="lightgray",lty=2)	
    points(CHR.DATA$POS, CHR.DATA$DIR, pch=19,cex=0.2,col="black")
    abline(h = 0)
    abline(h = 5, col="red")
    abline(h = -5, col="red")
    abline(v = PEAK.DATA$POS.PEAK,col=COLOR)
    abline(v = PEAK.DATA$POS.MAX,col=COLOR, lty=2)
    abline(v = PEAK.DATA$POS.MIN,col=COLOR, lty=2)
  } 
}
dev.off()

##############################
# 9 - Effect Size Estimation #
##############################

#Get G' for each peak
YPS1000.PEAK.G.PRIME <- numeric(nrow(PEAK.MERGE[[7]]))
I <- 1:nrow(PEAK.MERGE[[7]])
for(i in I) {
  CHR.G.PRIME <- DATA.G.PRIME[[7]][as.numeric(substr(DATA.G.PRIME[[7]]$CHR,4,5)) == PEAK.MERGE[[7]]$CHR[i],]
  YPS1000.PEAK.G.PRIME[i] <- mean(CHR.G.PRIME[which(abs(CHR.G.PRIME$POS - PEAK.MERGE[[7]]$POS.PEAK[i]) == min(abs(CHR.G.PRIME$POS - PEAK.MERGE[[7]]$POS.PEAK[i]))),]$DIR)
}

SK1.PEAK.G.PRIME <- numeric(nrow(PEAK.MERGE[[8]]))
I <- 1:nrow(PEAK.MERGE[[8]])
for(i in I) {
  CHR.G.PRIME <- DATA.G.PRIME[[8]][as.numeric(substr(DATA.G.PRIME[[8]]$CHR,4,5)) == PEAK.MERGE[[8]]$CHR[i],]
  SK1.PEAK.G.PRIME[i] <- mean(CHR.G.PRIME[which(abs(CHR.G.PRIME$POS - PEAK.MERGE[[8]]$POS.PEAK[i]) == min(abs(CHR.G.PRIME$POS - PEAK.MERGE[[8]]$POS.PEAK[i]))),]$DIR)
}

M22.PEAK.G.PRIME <- numeric(nrow(PEAK.MERGE[[9]]))
I <- 1:nrow(PEAK.MERGE[[9]])
for(i in I) {
  CHR.G.PRIME <- DATA.G.PRIME[[9]][as.numeric(substr(DATA.G.PRIME[[9]]$CHR,4,5)) == PEAK.MERGE[[9]]$CHR[i],]
  M22.PEAK.G.PRIME[i] <- mean(CHR.G.PRIME[which(abs(CHR.G.PRIME$POS - PEAK.MERGE[[9]]$POS.PEAK[i]) == min(abs(CHR.G.PRIME$POS - PEAK.MERGE[[9]]$POS.PEAK[i]))),]$DIR)
}

YPS1000.RELATIVE <- 1.044
SK1.RELATIVE <- 1.467
M22.RELATIVE <- 1.013

YPS1000.SUM <- sum(YPS1000.PEAK.G.PRIME)
SK1.SUM <- sum(SK1.PEAK.G.PRIME)
M22.SUM <- sum(M22.PEAK.G.PRIME)

YPS1000.EFFECT <- YPS1000.RELATIVE/YPS1000.SUM
SK1.EFFECT <- SK1.RELATIVE/SK1.SUM
M22.EFFECT <- M22.RELATIVE/M22.SUM

par(mfrow=c(3,1))
hist(YPS1000.EFFECT*YPS1000.PEAK.G.PRIME,breaks=seq(-.8,.8,0.025),xlim=c(-0.8,0.8),main="YPS1000",xlab="EFFECT", col="deepskyblue1")
hist(SK1.EFFECT*SK1.PEAK.G.PRIME,breaks=seq(-.8,.8,0.025),xlim=c(-0.8,0.8),main="SK1",xlab="EFFECT",col="darkgreen")
hist(M22.EFFECT*M22.PEAK.G.PRIME,breaks=seq(-.8,.8,0.025),xlim=c(-0.8,0.8),main="M22",xlab="EFFECT",col="darkgoldenrod4")

###Comparison to Albert et al.
ALBERT <- read.table("Albert.txt", header=TRUE)

#Plot G' and merged peaks 
N <- 9
J <- 1:16
D <- DATA.G.PRIME[[n]]
PEAKS <- PEAK.MERGE[[n]]
COLOR <- "goldenrod3"
  
#G' across the genome
par(mfrow=c(1,1))
par(mar=c(5.1,4.1,4.1,2.1))
plot(0,xlim=c(0,12080000),ylim=c(-1*max(abs(D$DIR)),max(abs(D$DIR))),type="n",xaxt="n",xlab="Position (Mb)",ylab="G'",main="")
for(j in J) {
  #Plot G.Prime
  G.DATA <- subset(D,D$CHR==paste("chr",sprintf("%02d", j),sep=""))
  points(G.DATA$POS+sum(CHROMOSOMES[0:(j-1),]$LENGTH), G.DATA$DIR, type="l",col="black")
  abline(v=sum(CHROMOSOMES[0:(j-1),]$LENGTH),lty=2,col="gray")
    
  #Add called eQTL
  PEAK.DATA <- subset(PEAKS,PEAKS$CHR == j)
  PEAK.DATA.UP   <- subset(PEAK.DATA, PEAK.DATA$DIR == 1)
  PEAK.DATA.DOWN <- subset(PEAK.DATA, PEAK.DATA$DIR == -1)
    
  if(nrow(PEAK.DATA.UP) > 0) {
    segments(x0 = PEAK.DATA.UP$POS.PEAK+sum(CHROMOSOMES[0:(j-1),]$LENGTH), y0=max(abs(D$DIR)), x1 = PEAK.DATA.UP$POS.PEAK+sum(CHROMOSOMES[0:(j-1),]$LENGTH), y1 = 0, col=COLOR)
    segments(x0 = PEAK.DATA.UP$POS.MAX +sum(CHROMOSOMES[0:(j-1),]$LENGTH), y0=max(abs(D$DIR)), x1 = PEAK.DATA.UP$POS.MAX+sum(CHROMOSOMES[0:(j-1),]$LENGTH), y1 = 0, col=COLOR, lty=2)
    segments(x0 = PEAK.DATA.UP$POS.MIN +sum(CHROMOSOMES[0:(j-1),]$LENGTH), y0=max(abs(D$DIR)), x1 = PEAK.DATA.UP$POS.MIN+sum(CHROMOSOMES[0:(j-1),]$LENGTH), y1 = 0, col=COLOR, lty=2)
  } 
  if(nrow(PEAK.DATA.DOWN) > 0) {
    segments(x0 = PEAK.DATA.DOWN$POS.PEAK+sum(CHROMOSOMES[0:(j-1),]$LENGTH), y0=-1*max(abs(D$DIR)), x1 = PEAK.DATA.DOWN$POS.PEAK+sum(CHROMOSOMES[0:(j-1),]$LENGTH), y1 = 0, col=COLOR)
    segments(x0 = PEAK.DATA.DOWN$POS.MAX +sum(CHROMOSOMES[0:(j-1),]$LENGTH), y0=-1*max(abs(D$DIR)), x1 = PEAK.DATA.DOWN$POS.MAX+sum(CHROMOSOMES[0:(j-1),]$LENGTH), y1 = 0, col=COLOR, lty=2)
    segments(x0 = PEAK.DATA.DOWN$POS.MIN +sum(CHROMOSOMES[0:(j-1),]$LENGTH), y0=-1*max(abs(D$DIR)), x1 = PEAK.DATA.DOWN$POS.MIN+sum(CHROMOSOMES[0:(j-1),]$LENGTH), y1 = 0, col=COLOR, lty=2)
  }
    
    K <- 1:nrow(ALBERT)
    for(k in K) {
      if(ALBERT$CHR[k] == j) {
        if(ALBERT$DIFF[k] < 0) {
          segments(x0 = ALBERT$PEAK[k] + sum(CHROMOSOMES[0:(j-1),]$LENGTH),y0=-1*max(abs(D$DIR)), x1 = ALBERT$PEAK[k] + sum(CHROMOSOMES[0:(j-1),]$LENGTH), y1 = 0, col="black",lwd=2)
          segments(x0 = ALBERT$MIN[k] + sum(CHROMOSOMES[0:(j-1),]$LENGTH), y0=-1*max(abs(D$DIR)), x1 = ALBERT$MIN[k] + sum(CHROMOSOMES[0:(j-1),]$LENGTH), y1 = 0, col="black",lty=2)
          segments(x0 = ALBERT$MAX[k] + sum(CHROMOSOMES[0:(j-1),]$LENGTH), y0=-1*max(abs(D$DIR)), x1 = ALBERT$MAX[k] + sum(CHROMOSOMES[0:(j-1),]$LENGTH), y1 = 0, col="black",lty=2)
        }
        if(ALBERT$DIFF[k] > 0) {
          segments(x0 = ALBERT$PEAK[k] + sum(CHROMOSOMES[0:(j-1),]$LENGTH),y0=1*max(abs(D$DIR)), x1 = ALBERT$PEAK[k] + sum(CHROMOSOMES[0:(j-1),]$LENGTH), y1 = 0, col="black",lwd=2)
          segments(x0 = ALBERT$MIN[k] + sum(CHROMOSOMES[0:(j-1),]$LENGTH), y0=1*max(abs(D$DIR)), x1 = ALBERT$MIN[k] + sum(CHROMOSOMES[0:(j-1),]$LENGTH), y1 = 0, col="black",lty=2)
          segments(x0 = ALBERT$MAX[k] + sum(CHROMOSOMES[0:(j-1),]$LENGTH), y0=1*max(abs(D$DIR)), x1 = ALBERT$MAX[k] + sum(CHROMOSOMES[0:(j-1),]$LENGTH), y1 = 0, col="black",lty=2)
        }
      }
    }
}
abline(v = sum(CHROMOSOMES$LENGTH),lty=2,col="gray")
abline(h = 0)
abline(h = 5, col="red")
abline(h = -5, col="red")
axis(1,at=c(seq(0,12080000,1000000)),labels=seq(0,12,1))
axis(3,at=(2*cumsum(CHROMOSOMES$LENGTH)-CHROMOSOMES$LENGTH)/2,CHROMOSOME.LABELS,cex.axis=0.6)


#Plot G' for variable rounds of selection and crosses
N <- c(1:3)
J <- 1:16
pdf("Supp.Figure.3.pdf")
for(n in N) {
  par(mfrow=c(4,8))
  par(mar=c(1,1,1,1))
  
  if(n == 1) {
    T1 <- c(1,4,7)
    T2 <- c(16,13,10)
    COLOR <- c("lightblue","blue","darkblue")
  } else if (n == 2) {
    T1 <- c(2,5,8)
    T2 <- c(17,14,11)
    COLOR <- c("lightgreen","green","darkgreen")
  } else if (n == 3) {
    T1 <- c(3,6,9)
    T2 <- c(18,15,12)
    COLOR <- c("goldenrod3","tan","brown")
  }
  ABS.MAX <- max(c(abs(DATA.G.PRIME[[T1[1]]]$DIR), abs(DATA.G.PRIME[[T1[2]]]$DIR),abs(DATA.G.PRIME[[T1[3]]]$DIR)))
  for(j in J) {
    POS.MAX <- max(subset(DATA.G.PRIME[[T1[1]]],DATA.G.PRIME[[T1[1]]]$CHR == paste("chr",sprintf("%02d", j),sep=""))$POS)
    plot(0,xlim=c(0,POS.MAX),ylim=c(-1*ABS.MAX,ABS.MAX),type="n",xaxt="n",xlab="Position",ylab="G'",main=paste(c("YPS1000","SK1","M22")[[which(N == n)]]," chr",sprintf("%02d", j),sep=""))
    for(t in T1) {
      D <- subset(DATA.G.PRIME[[t]],DATA.G.PRIME[[t]]$CHR == paste("chr",sprintf("%02d", j),sep=""))
      points(D$POS, D$DIR, pch=19,cex=0.2,col= COLOR[which(t == T1)], type = "l")
    }
    abline(h = 0)
    abline(h = 5, col="darkred")
    abline(h =-5, col="darkred")
  }
  ABS.MAX <- max(c(abs(DATA.G.PRIME[[T2[1]]]$DIR), abs(DATA.G.PRIME[[T2[2]]]$DIR),abs(DATA.G.PRIME[[T2[3]]]$DIR)))
  for(j in J) {
    POS.MAX <- max(subset(DATA.G.PRIME[[T2[1]]],DATA.G.PRIME[[T2[1]]]$CHR == paste("chr",sprintf("%02d", j),sep=""))$POS)
    plot(0,xlim=c(0,POS.MAX),ylim=c(-1*ABS.MAX,ABS.MAX),type="n",xaxt="n",xlab="Position",ylab="G'",main=paste(c("YPS1000","SK1","M22")[[which(N == n)]]," chr",sprintf("%02d", j),sep=""))
    for(t in T2) {
      D <- subset(DATA.G.PRIME[[t]],DATA.G.PRIME[[t]]$CHR == paste("chr",sprintf("%02d", j),sep=""))
      points(D$POS, D$DIR, pch=19,cex=0.2,col= COLOR[which(t == T2)], type = "l")
    }
    abline(h = 0)
    abline(h = 5, col="darkred")
    abline(h =-5, col="darkred")
  }
}
dev.off()


######Correlation of G' Stat Test######

X3.S3 <- subset(PEAK.MERGE[[9]])
colnames(X3.S3)[2] <- "POS"
X1.S1 <- subset(DATA.G.PRIME[[12]])
X1.S2 <- subset(DATA.G.PRIME[[15]])
X1.S3 <- subset(DATA.G.PRIME[[18]])
X3.S1 <- subset(DATA.G.PRIME[[3]])
X3.S2 <- subset(DATA.G.PRIME[[6]])

X1.S1$CHR <- as.integer(substr(X1.S1$CHR,4,5))
X1.S2$CHR <- as.integer(substr(X1.S2$CHR,4,5))
X1.S3$CHR <- as.integer(substr(X1.S3$CHR,4,5))
X3.S1$CHR <- as.integer(substr(X3.S1$CHR,4,5))
X3.S2$CHR <- as.integer(substr(X3.S2$CHR,4,5))

X1.S1 <- X1.S1[,c(1,2,14)]
X1.S2 <- X1.S2[,c(1,2,14)]
X1.S3 <- X1.S3[,c(1,2,14)]
X3.S1 <- X3.S1[,c(1,2,14)]
X3.S2 <- X3.S2[,c(1,2,14)]
X3.S3 <- X3.S3[,c(1,2,6)]

M.test <- merge(X3.S3,X1.S1,by.x=c("CHR","POS"), by.y =c("CHR","POS"))
M.test <- merge(M.test,X1.S2,by.x=c("CHR","POS"), by.y =c("CHR","POS"))
M.test <- merge(M.test,X1.S3,by.x=c("CHR","POS"), by.y =c("CHR","POS"))
M.test <- merge(M.test,X3.S1,by.x=c("CHR","POS"), by.y =c("CHR","POS"))
M.test <- merge(M.test,X3.S2,by.x=c("CHR","POS"), by.y =c("CHR","POS"))
colnames(M.test) <- c("CHR","POS","X3.S3","X1.S1","X1.S2","X1.S3","X3.S1","X3.S2")


M.test1 <- M.test
M.test2 <- M.test
M.test3 <- M.test


M.testing <- rbind(M.test1,M.test2,M.test3)

cor.test(M.testing$X3.S3, M.testing$X3.S1, method="spearman")
cor.test(M.testing$X3.S3, M.testing$X1.S3, method="spearman")


plot(M.testing$X3.S3,M.testing$X1.S3, pch = 19, cex=0.6,xlab="3 cross, 3 select",ylab="",main="M22",col="red",xlim=c(-200,200),ylim=c(-40,40))
points(M.testing$X3.S3,M.testing$X3.S1, pch = 19, cex=0.6)

abline(h=0)
abline(h=5,lty=2)
abline(h=-5,lty=2)
abline(v=0)
abline(v=5,lty=2)
abline(v=-5,lty=2)
```
