## Supplementary material for "Complex polymorphic genetic architecture underlies *trans*-regulatory variation among strains of *Saccharomyces cerevisiae*": Data S1: Functions.R

```
###########################################
# Functions for Metzger and Wittkopp 2019 #
###########################################

# 1 - Rotation of FCS data - ROT(x,Rotation)
# 2 - Hard gate calibration - GATE.CALIB(x,logFSC.A.MIN,logFSC.A.MAX,logFSC.H.MIN,logFSC.H.MAX,FSC.A_FSC.H.MIN,FSC.A_FSC.H.MAX,Width.MIN,Width.MAX)
# 3 - Cleaning FCS data - CLEANING(x,GATES)
# 4 - Quality control plots - FCS.PLOT(x)
# 5 - Perform plate correction - PLATE.CORRECT(DATA.NAME, CTRL.STRAIN, PLOT)
# 6 - Get strain estimates - STRAIN.ESTIMATES(DATA.NAME, WT.STRAIN, NEG.STRAIN, N.PERM) {
# 7 - Geometric average - GEOMEAN(x, na.rm = FALSE, trim = 0, ...)
# 8 - Geometric standard deviation - GEOSD(x, na.rm = FALSE, ...)
# 9 - Probability density for arbitrary distribution DMUTATION.MEAN(x, base.dist = TRANS.STRAIN.2$MEAN)
# 10- Calculating overlap of eQTL OVERLAP(x)
# 11- Calculating significance of eQTL overlap OVERLAP.TEST(x, N)

#--------------------------------------------------------------------------------------------------------------------------------#

  ##########################
### 1-ROTATION OF FCS DATA ###
  ##########################
   
ROT <- function(x,Rotation){
	Result <- Rotation%*%x
	return(Result)
}

#--------------------------------------------------------------------------------------------------------------------------------#

  ###########################
### 2-HARD GATE CALIBRATION ###
  ###########################

GATE.CALIB <- function(x,logFSC.A.MIN,logFSC.A.MAX,logFSC.H.MIN,logFSC.H.MAX,FSC.A_FSC.H.MIN,FSC.A_FSC.H.MAX,Width.MIN,Width.MAX) {

	if (missing(logFSC.A.MIN)) {logFSC.A.MIN <- 5.1}
	if (missing(logFSC.A.MAX)) {logFSC.A.MAX <- 6.1}
	if (missing(logFSC.H.MIN)) {logFSC.H.MIN <- 5.2}
	if (missing(logFSC.H.MAX)) {logFSC.H.MAX <- 6.8}
	if (missing(FSC.A_FSC.H.MIN)) {FSC.A_FSC.H.MIN <- 0.88}
	if (missing(FSC.A_FSC.H.MAX)) {FSC.A_FSC.H.MAX <- 0.94}
	if (missing(Width.MIN)) {Width.MIN <- 30}
	if (missing(Width.MAX)) {Width.MAX <- 80}
	
	Merge.Frame <- read.FCS(x,transformation=FALSE,alter.names=TRUE)
		
	##############################	
	##Log transformation of data##
	##############################
	
	logTrans <- logTransform(transformationId="log10-transformation",logbase=10,r=1,d=1)
	Merge.Frame <- transform(Merge.Frame,`logFSC.A`=logTrans(`FSC.A`))
	Merge.Frame <- transform(Merge.Frame,`logFSC.H`=logTrans(`FSC.H`))
	Merge.Frame <- transform(Merge.Frame,`logFL1.A`=logTrans(`FL1.A`))
	Merge.Frame <- transform(Merge.Frame,`logFL1.H`=logTrans(`FL1.H`))
	
	####################################
	##Calculate phenotypes of interest##
	####################################
	
	Data.Fluo <- as.data.frame(exprs(Merge.Frame))
	Data.Fluo[Data.Fluo == 0] <- 1
	
	Phenotype3 <- Data.Fluo[,"logFL1.A"]/Data.Fluo[,"logFSC.A"]
	Phenotype3 <- as.matrix(Phenotype3)
	colnames(Phenotype3) <- "FL1/FSC"
	Merge.Frame <- cbind2(Merge.Frame, Phenotype3)
	
	Phenotype4 <- (Data.Fluo[,"logFSC.A"])/(Data.Fluo[,"logFSC.H"])
	Phenotype4 <- as.matrix(Phenotype4)
	colnames(Phenotype4) <- "FSC.A/FSC.H"
	Merge.Frame <- cbind2(Merge.Frame, Phenotype4)		
	
	PlotAll <- as.data.frame(exprs(Merge.Frame))
	
	
	#############################
	###Quick plot to set gates###
	#############################
	
	# #Quick plot to set gates
	# quartz(height=14,width=14)
	par(mfrow=c(2,2))
	
	plot(PlotAll[,"Width"],PlotAll[,"logFSC.A"],pch=20,cex=0.5,col="#00000022",xlim=c(20,100),ylim=c(4,7))
	abline(v=Width.MIN)
	abline(v=Width.MAX)
	abline(h=logFSC.A.MIN)
	abline(h=logFSC.A.MAX)
	
	plot(PlotAll[,"logFSC.A"],PlotAll[,"logFL1.A"],pch=20,cex=0.5,col="#00000022",xlim=c(4,7),ylim=c(1.5,6.5))
	abline(v=logFSC.A.MIN)
	abline(v=logFSC.A.MAX)
	
	plot(PlotAll[,"logFSC.H"],PlotAll[,"logFSC.A"],pch=20,cex=0.4,col="#00000022",xlim=c(3,8),ylim=c(4,7))
	abline(v=logFSC.H.MIN)
	abline(v=logFSC.H.MAX)
	abline(h=logFSC.A.MIN)
	abline(h=logFSC.A.MAX)
	
	plot(PlotAll[,"logFSC.A"],PlotAll[,"FSC.A/FSC.H"],pch=20,cex=0.4,col="#00000022",xlim=c(3,8))
	abline(v=logFSC.A.MIN)
	abline(v=logFSC.A.MAX)
	abline(h=FSC.A_FSC.H.MIN)
	abline(h=FSC.A_FSC.H.MAX)
	
	OUTPUT <- c(logFSC.A.MIN,logFSC.A.MAX,logFSC.H.MIN,logFSC.H.MAX,FSC.A_FSC.H.MIN,FSC.A_FSC.H.MAX,Width.MIN,Width.MAX)
	names(OUTPUT) <- c("logFSC.A.MIN","logFSC.A.MAX","logFSC.H.MIN","logFSC.H.MAX","FSC.A_FSC.H.MIN","FSC.A_FSC.H.MAX","Width.MIN","Width.MAX")

	return(OUTPUT)
}


#--------------------------------------------------------------------------------------------------------------------------------#

  #######################
### 3-CLEANING FCS DATA ###
  #######################


CLEANING <- function(x,GATES) {

	Merge.Frame <- read.FCS(x["FILENAMES"],transformation=FALSE,alter.names=TRUE)
	
	OUTPUT <- data.frame(matrix(nrow=1,ncol=length(c("COUNTS.INITIAL",	"COUNTS.GATES",	"COUNTS.SINGLES",	"COUNTS.FINAL", "FSC.KURTOSIS",	"WIDTH",	"FSC.MEDIAN.INITIAL",	"FSC.MAD.INITIAL",	"FL1.MEDIAN.INITIAL",	"FL1.MAD.INITIAL",	"YFP.MEDIAN.INITIAL",	"YFP.MAD.INITIAL", "YFP.SD.INITIAL",	"INTERCEPT.INITIAL",	"SLOPE.INITIAL", "THETA", "YFP.MEDIAN.ROT","YFP.MAD.ROT","YFP.SD.ROT",	"FSC.MEDIAN.FINAL",	"FSC.MAD.FINAL",	"YFP.MEDIAN.FINAL",	"YFP.MAD.FINAL", "YFP.SD.FINAL","log.YFP.MEDIAN","log.YFP.MAD","log.YFP.SD"))))
	colnames(OUTPUT) <- c("COUNTS.INITIAL",	"COUNTS.GATES",	"COUNTS.SINGLES",	"COUNTS.FINAL", "FSC.KURTOSIS",	"WIDTH",	"FSC.MEDIAN.INITIAL",	"FSC.MAD.INITIAL",	"FL1.MEDIAN.INITIAL",	"FL1.MAD.INITIAL",	"YFP.MEDIAN.INITIAL",	"YFP.MAD.INITIAL", "YFP.SD.INITIAL",	"INTERCEPT.INITIAL",	"SLOPE.INITIAL", "THETA", "YFP.MEDIAN.ROT","YFP.MAD.ROT","YFP.SD.ROT",	"FSC.MEDIAN.FINAL",	"FSC.MAD.FINAL",	"YFP.MEDIAN.FINAL",	"YFP.MAD.FINAL", "YFP.SD.FINAL","log.YFP.MEDIAN","log.YFP.MAD","log.YFP.SD")
		
	OUTPUT["COUNTS.INITIAL"] <- nrow(exprs(Merge.Frame))
	
	if (OUTPUT["COUNTS.INITIAL"] > 1500 & x["SKIP"] == "NO")	{
		
		##############################	
		##Log transformation of data##
		##############################
		
		Start.exp <- exprs(Merge.Frame)
		Start.exp[,"FL1.A"] <- Start.exp[,"FL1.A"] + 10
		
		Merge.Frame <- new("flowFrame",Start.exp)
		
		Merge.Frame <- transform(Merge.Frame,`logFSC.A`=logTrans(`FSC.A`))
		Merge.Frame <- transform(Merge.Frame,`logFSC.H`=logTrans(`FSC.H`))
		Merge.Frame <- transform(Merge.Frame,`logFL1.A`=logTrans(`FL1.A`))
		Merge.Frame <- transform(Merge.Frame,`logFL1.H`=logTrans(`FL1.H`))
		
		####################################
		##Calculate phenotypes of interest##
		####################################
		
		Data.Fluo <- as.data.frame(exprs(Merge.Frame))
		Data.Fluo[Data.Fluo == 0] <- 1
		
		Phenotype3 <- Data.Fluo[,"logFL1.A"]/Data.Fluo[,"logFSC.A"]
		Phenotype3 <- as.matrix(Phenotype3)
		colnames(Phenotype3) <- "YFP.INITIAL"
		Merge.Frame <- cbind2(Merge.Frame, Phenotype3)
		
		Phenotype4 <- (Data.Fluo[,"logFSC.A"])/(Data.Fluo[,"logFSC.H"])
		Phenotype4 <- as.matrix(Phenotype4)
		colnames(Phenotype4) <- "FSC.A/FSC.H"
		Merge.Frame <- cbind2(Merge.Frame, Phenotype4)		
		
		PlotAll <- as.data.frame(exprs(Merge.Frame))	
	
		############
		#Hard Gates# 
		############
		
		rectGate <- rectangleGate(filterId="Noise Removal","logFSC.A"=c(GATES["logFSC.A.MIN"],GATES["logFSC.A.MAX"]), "logFSC.H"=c(GATES["logFSC.H.MIN"],GATES["logFSC.H.MAX"]), "FSC.A/FSC.H"=c(GATES["FSC.A_FSC.H.MIN"],GATES["FSC.A_FSC.H.MAX"]),"Width"=c(GATES["Width.MIN"],GATES["Width.MAX"]),"YFP.INITIAL"=c(min(PlotAll[,"YFP.INITIAL"]),max(PlotAll[,"YFP.INITIAL"])),"FL1.A"=c(11,max(PlotAll[,"FL1.A"])))
		
		Hard.Gates <- Subset(Merge.Frame, rectGate)
		Hard.Gates.exp <- exprs(Hard.Gates)

		OUTPUT["COUNTS.GATES"] <- nrow(Hard.Gates.exp)
	
		####################
		#Doublet Hard Gates# 
		####################
		
		Doublet.Model <- PCAgrid(cbind(Hard.Gates.exp[,"logFSC.A"],Hard.Gates.exp[,"FSC.A/FSC.H"]),k=2,method="sd",scores=TRUE,center="median")
		
		Scores <- Doublet.Model$scores
		
		Distri <- normalmixEM2comp(Scores[,2],sigsqrd=c(0.0022,0.0068)^2,mu=c(-0.0013,0.0086),lambda=c(0.56,0.44))
		
		Lambda <- Distri$lambda
		Mu <- Distri$mu
		Sigma <- Distri$sigma 
		
		Order <- c(which(Mu == min(Mu)),which(Mu == max(Mu)))
		
		Lambda <- Lambda[Order]
		Mu <- Mu[Order]
		Sigma <- Sigma[Order] 
		
		#Good cluster
		f <- function(x) dnorm(x,m=Mu[1],sd=Sigma[1])*Lambda[1]-dnorm(x,m=Mu[2],sd=Sigma[2])*Lambda[2]
		Threshold <- try(uniroot(f,interval=c(Mu[1],Mu[2]+Sigma[2]))$root)
		
		# if (Threshold > 0.1)
		# {
			# Threshold <- 0.075
		# }
			
		#Remove big cells based on FSC.A/FSC.H
		Position <- which(Scores[,2] < Threshold)
		Doublet.Gates.exp <- Hard.Gates.exp[Position,]
		Doublet.Gates <- new("flowFrame",Doublet.Gates.exp)
		
		#Remove cells with extreme FSC.A 
		DOUBLETS <- Doublet.Gates.exp[,"logFSC.A"]
		
		MEDIAN <- median(DOUBLETS)
		MAD <- mad(DOUBLETS)
		LOW <- MEDIAN - 2*MAD
		HIGH <- MEDIAN + 2*MAD
	
		NEW.MEDIAN <- MEDIAN
		OLD.MEDIAN <- median(DOUBLETS[which(DOUBLETS > LOW & DOUBLETS < HIGH)])
		
		while (abs(NEW.MEDIAN-OLD.MEDIAN) > 0.001)
		{
			NEW.DOUBLETS <- DOUBLETS[which(DOUBLETS > LOW & DOUBLETS < HIGH)]
			OLD.MEDIAN <- NEW.MEDIAN
			NEW.MEDIAN <- median(NEW.DOUBLETS)
			NEW.MAD <- mad(NEW.DOUBLETS)
			
			LOW <- NEW.MEDIAN - 2*NEW.MAD
			HIGH <- NEW.MEDIAN + 2*NEW.MAD
		}

		rectGate <- rectangleGate(filterId="Outliers logFSC.A","logFSC.A"=c(LOW,HIGH))
		
		Final.Doublets <- Subset(Doublet.Gates, rectGate)
		
		OUTPUT["FSC.KURTOSIS"] <- kurtosis(DOUBLETS)

		#################
		#Singles Cluster#
		#################
		
		Doublet.filter <- flowClust(Final.Doublets,varNames=c("logFSC.H","logFSC.A"),K=1,B=50,min.count=1000,nu.est=2,trans=0,seed=10,z.cutoff=0,level=0.9,tol=1e-4)
		Well.pop <- split(Final.Doublets,Doublet.filter,population=list(sc1=1))
		Well.Doublet <- Well.pop$sc1
		
		Doublets.exp <- as.data.frame(exprs(Well.Doublet))
						
		OUTPUT["COUNTS.SINGLES"] <- nrow(Doublets.exp)
		
		# plot(Hard.Gates.exp[,"logFSC.A"],Hard.Gates.exp[,"logFSC.H"],pch=20,col="#00000066")
		# points(Doublets.exp[,"logFSC.A"],Doublets.exp[,"logFSC.H"],pch=20,col="#FF000099")
		
		#############
		#Fluo filter#
		#############
	
		#REMOVE OUTLIERS FL1/FSC
		FLUO <- Doublets.exp[,"YFP.INITIAL"]
		
		MEDIAN <- median(FLUO)
		MAD <- mad(FLUO)
		LOW <- MEDIAN - 4*MAD
		HIGH <- MEDIAN + 4*MAD
	
		NEW.MEDIAN <- MEDIAN
		OLD.MEDIAN <- median(FLUO[which(FLUO > LOW & FLUO < HIGH)])
		
		while (abs(NEW.MEDIAN-OLD.MEDIAN) > 0.001)
		{
			NEW.FLUO <- FLUO[which(FLUO > LOW & FLUO < HIGH)]
			OLD.MEDIAN <- NEW.MEDIAN
			NEW.MEDIAN <- median(NEW.FLUO)
			NEW.MAD <- mad(NEW.FLUO)
			
			LOW <- NEW.MEDIAN - 4*NEW.MAD
			HIGH <- NEW.MEDIAN + 4*NEW.MAD
		}
	
		rectGate <- rectangleGate(filterId="Outliers FL1/FSC Removal","YFP.INITIAL"=c(LOW,HIGH))
		
		Hard.Fluo <- Subset(Well.Doublet, rectGate)
	
		Gate.Fluo <- flowClust(Hard.Fluo, varNames=c("logFSC.A","logFL1.A"),K=1,B=50,min.count=1000,nu.est=1,trans=0,z.cutoff=0.5,seed=10,tol=1e-5,nu=1.5,level=0.98)
		Well.pop <- split(Hard.Fluo,Gate.Fluo,population=list(sc1=1))
		Well.Fluo <- Well.pop$sc1
		Fluo.exp <- as.data.frame(exprs(Well.Fluo))

		#SAVE DATA
		OUTPUT["COUNTS.FINAL"] <- nrow(Fluo.exp)
		OUTPUT["WIDTH"] <- median(Fluo.exp[,"Width"])
		OUTPUT["FSC.MEDIAN.INITIAL"] <- median(Fluo.exp[,"logFSC.A"])
		OUTPUT["FSC.MAD.INITIAL"] <- mad(Fluo.exp[,"logFSC.A"])
		OUTPUT["FL1.MEDIAN.INITIAL"] <- median(Fluo.exp[,"logFL1.A"])
		OUTPUT["FL1.MAD.INITIAL"] <- mad(Fluo.exp[,"logFL1.A"])	
		OUTPUT["YFP.MEDIAN.INITIAL"] <- median(Fluo.exp[,"YFP.INITIAL"])
		OUTPUT["YFP.MAD.INITIAL"] <- mad(Fluo.exp[,"YFP.INITIAL"])
		OUTPUT["YFP.SD.INITIAL"] <- sd(Fluo.exp[,"YFP.INITIAL"])
		
		# plot(Doublets.exp$logFSC.A,Doublets.exp$logFL1.A,pch=20,col="#00000066")
		# points(Fluo.exp$logFSC.A,Fluo.exp$logFL1.A,pch=20,col="#FF000099")		
		
		###############################################################
		#####Remove correlation between logFSC.A and FL1.A/FSC.A#######
		###############################################################
		
		#1-Define orthogonal regression
		Intercept <- c()
		Slope <- c()
		Theta <- c()
		
		Fluo.Model <- PCAgrid(cbind(Fluo.exp[,"logFSC.A"],Fluo.exp[,"YFP.INITIAL"]),k=2,method="sd",scores=FALSE,center="median")
		
		#2-Center of rotation
		x.center <- Fluo.Model$center[1]
		y.center <- Fluo.Model$center[2]
				
		#3-Initial Intercept and Slope
		Slope[1] <- Fluo.Model$loadings[2,1] / Fluo.Model$loadings[1,1]
		Intercept[1] <- Fluo.Model$center[2] - Slope[1]*Fluo.Model$center[1]
				
		#4-Calculate angle of rotation
		a <- c(x.center-0,y.center-Intercept[1]) #Vector from Intercept to Centroid
		b <- c(x.center-0,y.center-y.center) #Vector with slope 0 through Centroid
				
		Theta[1] <- acos(sum(a*b)/(sqrt(sum(a*a))*sqrt(sum(b*b)))) #Angle between 2 vectors
				
		if (Slope[1] < 0)
		{
			Theta[1] <- -Theta[1]
		}		
				
		#5-Define rotation matrix
		Rotation <- matrix(c(cos(Theta[1]),-sin(Theta[1]),sin(Theta[1]),cos(Theta[1])),ncol=2,nrow=2)
	
		#6-Transform Data
		Coord <- t(as.matrix(Fluo.exp[,c("logFSC.A","YFP.INITIAL")]))
		
		Coord[1,] <- Coord[1,] - x.center
		Coord[2,] <- Coord[2,] - y.center
		
		Result <- ROT(x=Coord,Rotation=Rotation)
		
		Result[1,] <- Result[1,] + x.center
		Result[2,] <- Result[2,] + y.center
		
		#7-Keep record of rotated values
		Fluo.exp[,"FSC.FINAL"] <- Result[1,]
		Fluo.exp[,"YFP.ROT"] <- Result[2,]
		
		
		###########################################################################
		#Apply correction for linear relation between fluorescence and mRNA levels#
		###########################################################################
		
		for (j in 1:nrow(Fluo.exp))	
		{
		  Fluo.exp[j,"YFP.FINAL"]  <-  exp(7.820027*Fluo.exp[j,"YFP.ROT"])
		}
		
		OUTPUT["INTERCEPT.INITIAL"] <- Intercept[1]
		OUTPUT["SLOPE.INITIAL"] <- Slope[1]
		OUTPUT["THETA"] <- Theta[1]
		OUTPUT["YFP.MEDIAN.ROT"] <- median(Fluo.exp[,"YFP.ROT"])
		OUTPUT["YFP.MAD.ROT"] <- mad(Fluo.exp[,"YFP.ROT"])
		OUTPUT["YFP.SD.ROT"] <- sd(Fluo.exp[,"YFP.ROT"])
		OUTPUT["FSC.MEDIAN.FINAL"] <- median(Fluo.exp[,"FSC.FINAL"])
		OUTPUT["FSC.MAD.FINAL"] <- mad(Fluo.exp[,"FSC.FINAL"])	
		OUTPUT["YFP.MEDIAN.FINAL"] <- median(Fluo.exp[,"YFP.FINAL"])
		OUTPUT["YFP.MAD.FINAL"] <- mad(Fluo.exp[,"YFP.FINAL"])	
		OUTPUT["YFP.SD.FINAL"] <- sd(Fluo.exp[,"YFP.FINAL"])	
		OUTPUT["log.YFP.MEDIAN"] <- median(log(Fluo.exp[,"YFP.FINAL"]))
		OUTPUT["log.YFP.MAD"] <- mad(log(Fluo.exp[,"YFP.FINAL"]))	
		OUTPUT["log.YFP.SD"] <- sd(log(Fluo.exp[,"YFP.FINAL"]))	
		
		x["COUNTER"]
		print(x["COUNTER"])
		
		#Save clean data with rotation correction
		Data <- Fluo.exp[,c("Width","Time","logFSC.A","logFSC.H","logFL1.A","logFL1.H","YFP.INITIAL","YFP.ROT","FSC.FINAL","YFP.FINAL")]
		write.table(Data,file=paste("CLEAN.DATA/","Day",x["DAY"],"_Rep",x["REP.UNIQUE"],"_Plate",x["PLATE"],"_Well",x["POSITION"],".txt",sep=""),row.names=FALSE,col.names=TRUE,sep="\t",quote=FALSE)

		return(OUTPUT)	
		
	} else {
		OUTPUT[] <- NA
		OUTPUT["COUNTS.INITIAL"] <- nrow(exprs(Merge.Frame))
		
		Data <- as.data.frame(matrix(data=NA,ncol=10))
		colnames(Data) <- c("Width","Time","logFSC.A","logFSC.H","logFL1.A","logFL1.H","YFP.INITIAL","YFP.ROT","FSC.FINAL","YFP.FINAL")
		write.table(Data,file=paste("CLEAN.DATA/","Day",x["DAY"],"_Rep",x["REP.UNIQUE"],"_Plate",x["PLATE"],"_Well",x["POSITION"],".txt",sep=""),row.names=FALSE,col.names=TRUE,sep="\t",quote=FALSE)
		
		return(OUTPUT)
	}
	
}


#--------------------------------------------------------------------------------------------------------------------------------#

  ###########################
### 4-QUALITY CONTROL PLOTS ###
  ###########################


FCS.PLOT <- function(x) {
	
	INITIAL.DATA <- read.FCS(x["INITIAL"],transformation=FALSE,alter.names=TRUE)
	CLEAN.DATA <-  read.table(x["CLEAN"],header=TRUE,as.is=TRUE)

	##############################	
	##Log transformation of data##
	##############################
		
	Merge.Frame <- transform(INITIAL.DATA,`logFSC.A`=logTrans(`FSC.A`))
	Merge.Frame <- transform(Merge.Frame,`logFSC.H`=logTrans(`FSC.H`))
	Merge.Frame <- transform(Merge.Frame,`logFL1.A`=logTrans(`FL1.A`))
	Merge.Frame <- transform(Merge.Frame,`logFL1.H`=logTrans(`FL1.H`))
		
	####################################
	##Calculate phenotypes of interest##
	####################################
		
	Data.Fluo <- as.data.frame(exprs(Merge.Frame))
	Data.Fluo[Data.Fluo == 0] <- 1
		
	Phenotype3 <- Data.Fluo[,"logFL1.A"]/Data.Fluo[,"logFSC.A"]
	Phenotype3 <- as.matrix(Phenotype3)
	colnames(Phenotype3) <- "YFP.INITIAL"
	Merge.Frame <- cbind2(Merge.Frame, Phenotype3)
		
	Phenotype4 <- (Data.Fluo[,"logFSC.A"])/(Data.Fluo[,"logFSC.H"])
	Phenotype4 <- as.matrix(Phenotype4)
	colnames(Phenotype4) <- "FSC.A/FSC.H"
	Merge.Frame <- cbind2(Merge.Frame, Phenotype4)		
		
	PLOT.ALL <- as.data.frame(exprs(Merge.Frame))

	NAME <- paste("P",x["PLATE"],"R",x["REP"],"_",x["POSITION"],"_",x["STRAIN"],"_",x["CLASS"],sep="")
	

	pdf(paste("CLEANING.PLOTS/",NAME,".pdf",sep=""))

	par(mfrow=c(2,2))

	plot(PLOT.ALL$logFSC.A,PLOT.ALL$logFSC.H,pch=20,cex=0.5,col="#00000033",xlim=c(4.3,6.5),ylim=c(5,7),xlab="logFSC.A",ylab="logFSC.H",main=NAME)
	points(CLEAN.DATA$logFSC.A,CLEAN.DATA$logFSC.H,pch=20,cex=0.5,col="#FF000066",main=NAME)

	plot(PLOT.ALL$logFSC.A,PLOT.ALL$logFL1.A,pch=20,cex=0.5,col="#00000033",xlim=c(4.3,6.5),ylim=c(1.5,6.5),xlab="logFSC.A",ylab="logFL1.A",main=NAME)
	points(CLEAN.DATA$logFSC.A,CLEAN.DATA$logFL1.A,pch=20,cex=0.5,col="#FF000066",main=NAME)

	plot(CLEAN.DATA$logFSC.A,CLEAN.DATA$YFP.INITIAL,pch=20,cex=0.5,col="#FF0000AA",xlab="logFSC.A",ylab="FL1/FSC",main=NAME)
	abline(lm(CLEAN.DATA$YFP.INITIAL~CLEAN.DATA$logFSC.A),lty=2,col="red",lwd=2.5)
	points(CLEAN.DATA$FSC.FINAL,CLEAN.DATA$YFP.FINAL,pch=20,cex=0.5,col="#00FF00AA",main=NAME)
	abline(lm(CLEAN.DATA$YFP.FINAL~CLEAN.DATA$FSC.FINAL),lty=2,col="green",lwd=2.5)
	
	hist(CLEAN.DATA$YFP.FINAL,breaks=50,main=NAME,xlab="FL1/FSC")

	dev.off()	

}	
	
#--------------------------------------------------------------------------------------------------------------------------------#

  #################################
### 5 - PLATE CORRECTION FUNCTION ###
  #################################

PLATE.CORRECT <- function(DATA.NAME, CTRL.STRAIN, PLOT) {
  #Set read-in data types and constants
  DATA.TYPE <- c(rep("factor",19),rep("integer",5),rep("numeric",23))
  REF <- 1
  NEG <- 0
  
  #Read data
  DATA <- read.table(paste(DATA.NAME,".CLEAN.txt",sep=""),header=TRUE,colClasses=DATA.TYPE)
 
  #Set data types and transformations
  DATA[,"log.RNA.MEDIAN"] <- log(((DATA[,"YFP.MEDIAN.FINAL"] - NEG) / (REF - NEG)) + 0.05)
  DATA[,"log.RNA.SD"] <- DATA[,"YFP.SD.FINAL"] / ((DATA[,"YFP.MEDIAN.FINAL"] - NEG) + (REF - NEG)*0.05)
  DATA[,"FLOW.RUN"] <- as.factor(DATA[,"FLOW.RUN"])
  DATA[,"ROW"] <- as.factor(DATA[,"ROW"])
  
  #Subset complete cases and remove known bad samples
  WORKING.DATA <- subset(DATA, DATA$COUNTS.FINAL > 1000)
  
  if(DATA.NAME == "CIS") {
    WORKING.DATA <- droplevels(subset(WORKING.DATA,WORKING.DATA$DAY != 1))
    WORKING.DATA <- droplevels(subset(WORKING.DATA,!(WORKING.DATA$FLOW.RUN == "146" & WORKING.DATA$ROW == "C")))
    WORKING.DATA <- droplevels(subset(WORKING.DATA,!(WORKING.DATA$FLOW.RUN == "109" & WORKING.DATA$ROW == "B")))
  } else if (DATA.NAME == "TRANS.1") {
    WORKING.DATA <- droplevels(subset(WORKING.DATA,WORKING.DATA$FLOW.RUN != "042"))
  }
  WORKING.DATA <- droplevels(WORKING.DATA)
  WORKING.DATA <- WORKING.DATA[complete.cases(WORKING.DATA),]

  ##CORRECT FOR FSC MEDIAN
  #Flow Run
  WORKING.CONTROL <- subset(WORKING.DATA,WORKING.DATA$STRAIN == CTRL.STRAIN)
  FSC.MEDIAN.CORRECT <- rlm(FSC.MEDIAN.FINAL ~ FLOW.RUN, data = WORKING.CONTROL)
  COEF.FLOW.RUN <- c(0,coefficients(FSC.MEDIAN.CORRECT)[grep("FLOW.RUN",names(coefficients(FSC.MEDIAN.CORRECT)))])
  I <- 1:nrow(WORKING.DATA)
  OUT <- numeric(length(I))
  for(i in I) {
    FLOW.RUN <- which(levels(WORKING.DATA$FLOW.RUN) == WORKING.DATA$FLOW.RUN[i])
    OUT[i] <- COEF.FLOW.RUN[FLOW.RUN]
  }
  WORKING.DATA[,"FSC.MEDIAN.CORRECT"] <- WORKING.DATA$FSC.MEDIAN.FINAL - OUT
  WORKING.CONTROL <- subset(WORKING.DATA,WORKING.DATA$STRAIN == CTRL.STRAIN)
  
  #Row
  WORKING.CONTROL <- subset(WORKING.DATA,WORKING.DATA$STRAIN == CTRL.STRAIN)
  FSC.MEDIAN.CORRECT <- rlm(FSC.MEDIAN.CORRECT ~ ROW, data = WORKING.CONTROL)
  COEF.ROW   <- c(0,coefficients(FSC.MEDIAN.CORRECT)[grep("ROW",names(coefficients(FSC.MEDIAN.CORRECT)))])
  I <- 1:nrow(WORKING.DATA)
  OUT <- numeric(length(I))
  for(i in I) {
    ROW <- which(levels(WORKING.DATA$ROW) == WORKING.DATA$ROW[i])
    OUT[i] <- COEF.ROW[ROW]
  }
  WORKING.DATA[,"FSC.MEDIAN.CORRECT"] <- WORKING.DATA$FSC.MEDIAN.CORRECT - OUT
  WORKING.CONTROL <- subset(WORKING.DATA,WORKING.DATA$STRAIN == CTRL.STRAIN)
  
  #Identify outliers in cell size for controls
  LOW <- median(WORKING.CONTROL$FSC.MEDIAN.CORRECT) - 4*mad(WORKING.CONTROL$FSC.MEDIAN.CORRECT)
  HIGH <- median(WORKING.CONTROL$FSC.MEDIAN.CORRECT) + 4*mad(WORKING.CONTROL$FSC.MEDIAN.CORRECT)
  WORKING.CONTROL <- subset(WORKING.CONTROL, FSC.MEDIAN.CORRECT >= LOW & FSC.MEDIAN.CORRECT <= HIGH)
  
  ##CORRECT FOR YFP MEDIAN
  #Flow Run
  WORKING.CONTROL <- subset(WORKING.DATA,WORKING.DATA$STRAIN == CTRL.STRAIN)
  WORKING.CONTROL <- subset(WORKING.CONTROL, FSC.MEDIAN.CORRECT >= LOW & FSC.MEDIAN.CORRECT <= HIGH)
  YFP.MEDIAN.CORRECT <- rlm(log.RNA.MEDIAN ~ 0 + FLOW.RUN, data = WORKING.CONTROL)
  COEF.FLOW.RUN   <- c(coefficients(YFP.MEDIAN.CORRECT)[grep("FLOW.RUN",names(coefficients(YFP.MEDIAN.CORRECT)))])
  I <- 1:nrow(WORKING.DATA)
  OUT <- numeric(length(I))
  for(i in I) {
    FLOW.RUN <- which(levels(WORKING.DATA$FLOW.RUN) == WORKING.DATA$FLOW.RUN[i])
    OUT[i] <- COEF.FLOW.RUN[FLOW.RUN]
  }
  WORKING.DATA[,"log.RNA.MEDIAN.CORRECT"] <- WORKING.DATA$log.RNA.MEDIAN - OUT + mean(WORKING.CONTROL$log.RNA.MEDIAN)
  WORKING.CONTROL <- subset(WORKING.DATA,WORKING.DATA$STRAIN == CTRL.STRAIN)
  WORKING.CONTROL <- subset(WORKING.CONTROL, FSC.MEDIAN.CORRECT >= LOW & FSC.MEDIAN.CORRECT <= HIGH)
  
  #Row
  WORKING.CONTROL <- subset(WORKING.DATA,WORKING.DATA$STRAIN == CTRL.STRAIN)
  WORKING.CONTROL <- subset(WORKING.CONTROL, FSC.MEDIAN.CORRECT >= LOW & FSC.MEDIAN.CORRECT <= HIGH)
  YFP.MEDIAN.CORRECT <- rlm(log.RNA.MEDIAN.CORRECT ~ 0 + ROW, data = WORKING.CONTROL)
  COEF.ROW   <- c(coefficients(YFP.MEDIAN.CORRECT)[grep("ROW",names(coefficients(YFP.MEDIAN.CORRECT)))])
  I <- 1:nrow(WORKING.DATA)
  OUT <- numeric(length(I))
  for(i in I) {
    ROW <- which(levels(WORKING.DATA$ROW) == WORKING.DATA$ROW[i])
    OUT[i] <- COEF.ROW[ROW]
  }
  WORKING.DATA[,"log.RNA.MEDIAN.CORRECT"] <- WORKING.DATA$log.RNA.MEDIAN.CORRECT - OUT + mean(WORKING.CONTROL$log.RNA.MEDIAN.CORRECT)
  WORKING.DATA[,"YFP.MEDIAN.CORRECT"] <- (exp(WORKING.DATA[,"log.RNA.MEDIAN.CORRECT"]) - 0.05) * (REF - NEG) + NEG
  WORKING.CONTROL <- subset(WORKING.DATA,WORKING.DATA$STRAIN == CTRL.STRAIN)
  WORKING.CONTROL <- subset(WORKING.CONTROL, FSC.MEDIAN.CORRECT >= LOW & FSC.MEDIAN.CORRECT <= HIGH)
  
  ##CORRECT FOR YFP MAD ON LOG SCALE
  #Flow Run
  WORKING.CONTROL <- subset(WORKING.DATA,WORKING.DATA$STRAIN == CTRL.STRAIN)
  WORKING.CONTROL <- subset(WORKING.CONTROL, FSC.MEDIAN.CORRECT >= LOW & FSC.MEDIAN.CORRECT <= HIGH)
  YFP.MAD.CORRECT <- rlm(log.RNA.SD ~ 0 + FLOW.RUN, data = WORKING.CONTROL)
  COEF.FLOW.RUN   <- c(coefficients(YFP.MAD.CORRECT)[grep("FLOW.RUN",names(coefficients(YFP.MAD.CORRECT)))])
  I <- 1:nrow(WORKING.DATA)
  OUT <- numeric(length(I))
  for(i in I) {
    FLOW.RUN <- which(levels(WORKING.DATA$FLOW.RUN) == WORKING.DATA$FLOW.RUN[i])
    OUT[i] <- COEF.FLOW.RUN[FLOW.RUN]
  }
  WORKING.DATA[,"log.RNA.SD.CORRECT"] <- WORKING.DATA$log.RNA.SD - OUT + mean(WORKING.CONTROL$log.RNA.SD) 
  WORKING.CONTROL <- subset(WORKING.DATA,WORKING.DATA$STRAIN == CTRL.STRAIN)
  WORKING.CONTROL <- subset(WORKING.CONTROL, FSC.MEDIAN.CORRECT >= LOW & FSC.MEDIAN.CORRECT <= HIGH)
  
  #Row
  WORKING.CONTROL <- subset(WORKING.DATA,WORKING.DATA$STRAIN == CTRL.STRAIN)
  WORKING.CONTROL <- subset(WORKING.CONTROL, FSC.MEDIAN.CORRECT >= LOW & FSC.MEDIAN.CORRECT <= HIGH)
  YFP.MAD.CORRECT <- rlm(log.RNA.SD.CORRECT ~ 0 + ROW, data = WORKING.CONTROL)
  COEF.ROW   <- c(coefficients(YFP.MAD.CORRECT)[grep("ROW",names(coefficients(YFP.MAD.CORRECT)))])
  I <- 1:nrow(WORKING.DATA)
  OUT <- numeric(length(I))
  for(i in I) {
    ROW <- which(levels(WORKING.DATA$ROW) == WORKING.DATA$ROW[i])
    OUT[i] <- COEF.ROW[ROW]
  }
  WORKING.DATA[,"log.RNA.SD.CORRECT"] <- WORKING.DATA$log.RNA.SD.CORRECT - OUT + mean(WORKING.CONTROL$log.RNA.SD.CORRECT)
  WORKING.DATA[,"YFP.SD.CORRECT"] <- WORKING.DATA[,"log.RNA.SD.CORRECT"] * ((WORKING.DATA[,"YFP.MEDIAN.CORRECT"] - NEG) + (REF - NEG)*0.05)
  WORKING.CONTROL <- subset(WORKING.DATA,WORKING.DATA$STRAIN == CTRL.STRAIN)
  WORKING.CONTROL <- subset(WORKING.CONTROL, FSC.MEDIAN.CORRECT >= LOW & FSC.MEDIAN.CORRECT <= HIGH)
  
  if(PLOT == TRUE) {
    pdf(paste(DATA.NAME,".CORRECTION.PLOTS.pdf",sep=""))  
    
    plot(WORKING.CONTROL$FSC.MEDIAN.FINAL   ~ WORKING.CONTROL$FLOW.RUN,pch=19,cex=0.6,xlab="Flow Run",ylab="FSC Median",main="Original")
    plot(WORKING.CONTROL$FSC.MEDIAN.CORRECT ~ WORKING.CONTROL$FLOW.RUN,pch=19,cex=0.6,xlab="Flow Run",ylab="FSC Median",main="Corrected")
    plot(WORKING.CONTROL$FSC.MEDIAN.CORRECT ~ WORKING.CONTROL$ROW,pch=19,cex=0.6,xlab="Row",ylab="FSC Median",main="Corrected")
    plot(WORKING.CONTROL$FSC.MEDIAN.CORRECT ~ WORKING.CONTROL$COL,pch=19,cex=0.6,xlab="Col",ylab="FSC Median",main="Corrected")
    plot(WORKING.CONTROL$FSC.MEDIAN.CORRECT ~ WORKING.CONTROL$POSITION,pch=19,cex=0.6,xlab="Position",ylab="FSC Median",main="Corrected")
    plot(WORKING.CONTROL$FSC.MEDIAN.CORRECT ~ WORKING.CONTROL$PLATE,pch=19,cex=0.6,xlab="Plate",ylab="FSC Median",main="Corrected")
    plot(WORKING.CONTROL$FSC.MEDIAN.CORRECT ~ WORKING.CONTROL$REP,pch=19,cex=0.6,xlab="Rep",ylab="FSC Median",main="Corrected")
    plot(WORKING.CONTROL$FSC.MEDIAN.CORRECT ~ WORKING.CONTROL$DAY,pch=19,cex=0.6,xlab="Day",ylab="FSC Median",main="Corrected")
    plot(WORKING.CONTROL$FSC.MEDIAN.CORRECT ~ WORKING.CONTROL$BLOCK,pch=19,cex=0.6,xlab="Block",ylab="FSC Median",main="Corrected")
    
    plot(WORKING.CONTROL$log.RNA.MEDIAN         ~ WORKING.CONTROL$FLOW.RUN,pch=19,cex=0.6,xlab="Flow Run",ylab="log(RNA)",main="Original")
    plot(WORKING.CONTROL$log.RNA.MEDIAN.CORRECT ~ WORKING.CONTROL$FLOW.RUN,pch=19,cex=0.6,xlab="Flow Run",ylab="log(RNA)",main="Corrected")
    plot(WORKING.CONTROL$log.RNA.MEDIAN.CORRECT ~ WORKING.CONTROL$ROW,pch=19,cex=0.6,xlab="Row",ylab="log(RNA)",main="Corrected")
    plot(WORKING.CONTROL$log.RNA.MEDIAN.CORRECT ~ WORKING.CONTROL$COL,pch=19,cex=0.6,xlab="Col",ylab="log(RNA)",main="Corrected")
    plot(WORKING.CONTROL$log.RNA.MEDIAN.CORRECT ~ WORKING.CONTROL$POSITION,pch=19,cex=0.6,xlab="Position",ylab="log(RNA)",main="Corrected")
    plot(WORKING.CONTROL$log.RNA.MEDIAN.CORRECT ~ WORKING.CONTROL$PLATE,pch=19,cex=0.6,xlab="Plate",ylab="log(RNA)",main="Corrected")
    plot(WORKING.CONTROL$log.RNA.MEDIAN.CORRECT ~ WORKING.CONTROL$REP,pch=19,cex=0.6,xlab="Rep",ylab="log(RNA)",main="Corrected")
    plot(WORKING.CONTROL$log.RNA.MEDIAN.CORRECT ~ WORKING.CONTROL$DAY,pch=19,cex=0.6,xlab="Day",ylab="log(RNA)",main="Corrected")
    plot(WORKING.CONTROL$log.RNA.MEDIAN.CORRECT ~ WORKING.CONTROL$BLOCK,pch=19,cex=0.6,xlab="Block",ylab="log(RNA)",main="Corrected")
    
    plot(WORKING.CONTROL$YFP.MEDIAN.FINAL   ~ WORKING.CONTROL$FLOW.RUN,pch=19,cex=0.6,xlab="Flow Run",ylab="YFP Median",main="Original")
    plot(WORKING.CONTROL$YFP.MEDIAN.CORRECT ~ WORKING.CONTROL$FLOW.RUN,pch=19,cex=0.6,xlab="Flow Run",ylab="YFP Median",main="Corrected")
    plot(WORKING.CONTROL$YFP.MEDIAN.CORRECT ~ WORKING.CONTROL$ROW,pch=19,cex=0.6,xlab="Row",ylab="YFP Median",main="Corrected")
    plot(WORKING.CONTROL$YFP.MEDIAN.CORRECT ~ WORKING.CONTROL$COL,pch=19,cex=0.6,xlab="Col",ylab="YFP Median",main="Corrected")
    plot(WORKING.CONTROL$YFP.MEDIAN.CORRECT ~ WORKING.CONTROL$POSITION,pch=19,cex=0.6,xlab="Position",ylab="YFP Median",main="Corrected")
    plot(WORKING.CONTROL$YFP.MEDIAN.CORRECT ~ WORKING.CONTROL$PLATE,pch=19,cex=0.6,xlab="Plate",ylab="YFP Median",main="Corrected")
    plot(WORKING.CONTROL$YFP.MEDIAN.CORRECT ~ WORKING.CONTROL$REP,pch=19,cex=0.6,xlab="Rep",ylab="YFP Median",main="Corrected")
    plot(WORKING.CONTROL$YFP.MEDIAN.CORRECT ~ WORKING.CONTROL$DAY,pch=19,cex=0.6,xlab="Day",ylab="YFP Median",main="Corrected")
    plot(WORKING.CONTROL$YFP.MEDIAN.CORRECT ~ WORKING.CONTROL$BLOCK,pch=19,cex=0.6,xlab="Block",ylab="YFP Median",main="Corrected")
    
    plot(WORKING.CONTROL$log.RNA.SD         ~ WORKING.CONTROL$FLOW.RUN,pch=19,cex=0.6,xlab="Flow Run",ylab="log(RNA) SD",main="Original")
    plot(WORKING.CONTROL$log.RNA.SD.CORRECT ~ WORKING.CONTROL$FLOW.RUN,pch=19,cex=0.6,xlab="Flow Run",ylab="log(RNA) SD",main="Corrected")
    plot(WORKING.CONTROL$log.RNA.SD.CORRECT ~ WORKING.CONTROL$ROW,pch=19,cex=0.6,xlab="Row",ylab="log(RNA) SD",main="Corrected")
    plot(WORKING.CONTROL$log.RNA.SD.CORRECT ~ WORKING.CONTROL$COL,pch=19,cex=0.6,xlab="Col",ylab="log(RNA) SD",main="Corrected")
    plot(WORKING.CONTROL$log.RNA.SD.CORRECT ~ WORKING.CONTROL$POSITION,pch=19,cex=0.6,xlab="Position",ylab="log(RNA) SD",main="Corrected")
    plot(WORKING.CONTROL$log.RNA.SD.CORRECT ~ WORKING.CONTROL$PLATE,pch=19,cex=0.6,xlab="Plate",ylab="log(RNA) SD",main="Corrected")
    plot(WORKING.CONTROL$log.RNA.SD.CORRECT ~ WORKING.CONTROL$REP,pch=19,cex=0.6,xlab="Rep",ylab="log(RNA) SD",main="Corrected")
    plot(WORKING.CONTROL$log.RNA.SD.CORRECT ~ WORKING.CONTROL$DAY,pch=19,cex=0.6,xlab="Day",ylab="log(RNA) SD",main="Corrected")
    plot(WORKING.CONTROL$log.RNA.SD.CORRECT ~ WORKING.CONTROL$BLOCK,pch=19,cex=0.6,xlab="Block",ylab="log(RNA) SD",main="Corrected")
    
    plot(WORKING.CONTROL$YFP.SD.FINAL   ~ WORKING.CONTROL$FLOW.RUN,pch=19,cex=0.6,xlab="Flow Run",ylab="YFP SD",main="Original")
    plot(WORKING.CONTROL$YFP.SD.CORRECT ~ WORKING.CONTROL$FLOW.RUN,pch=19,cex=0.6,xlab="Flow Run",ylab="YFP SD",main="Corrected")
    plot(WORKING.CONTROL$YFP.SD.CORRECT ~ WORKING.CONTROL$ROW,pch=19,cex=0.6,xlab="Row",ylab="YFP SD",main="Corrected")
    plot(WORKING.CONTROL$YFP.SD.CORRECT ~ WORKING.CONTROL$COL,pch=19,cex=0.6,xlab="Col",ylab="YFP SD",main="Corrected")
    plot(WORKING.CONTROL$YFP.SD.CORRECT ~ WORKING.CONTROL$POSITION,pch=19,cex=0.6,xlab="Position",ylab="YFP SD",main="Corrected")
    plot(WORKING.CONTROL$YFP.SD.CORRECT ~ WORKING.CONTROL$PLATE,pch=19,cex=0.6,xlab="Plate",ylab="YFP SD",main="Corrected")
    plot(WORKING.CONTROL$YFP.SD.CORRECT ~ WORKING.CONTROL$REP,pch=19,cex=0.6,xlab="Rep",ylab="YFP SD",main="Corrected")
    plot(WORKING.CONTROL$YFP.SD.CORRECT ~ WORKING.CONTROL$DAY,pch=19,cex=0.6,xlab="Day",ylab="YFP SD",main="Corrected")
    plot(WORKING.CONTROL$YFP.SD.CORRECT ~ WORKING.CONTROL$BLOCK,pch=19,cex=0.6,xlab="Block",ylab="YFP SD",main="Corrected")
    dev.off()
  }
  
  write.table(WORKING.DATA,paste(DATA.NAME,".CORRECT.txt",sep=""),row.names=FALSE,sep="\t",quote=FALSE)
  
}

#--------------------------------------------------------------------------------------------------------------------------------#

  ################################
### 6 - STRAIN ESTIMATES FUNCTION###
  ################################

STRAIN.ESTIMATES <- function(DATA.NAME, WT.STRAIN, NEG.STRAIN, N.PERM) {
  
  WORKING.DATA <- read.table(paste(DATA.NAME,".CORRECT.txt",sep=""),header=TRUE)
  WORKING.DATA <- WORKING.DATA[complete.cases(WORKING.DATA),]
  I <- 1:nrow(WORKING.DATA)
  
  if(DATA.NAME %in% c("TRANS.1","TRANS.2")) {
    WT.STRAINS <- which(WORKING.DATA$TREATMENT == "SHAM" & WORKING.DATA$TAIL == "ALL")
  } else {
    WT.STRAINS <- which(WORKING.DATA$STRAIN == WT.STRAIN)
  }
  
  #YFP Median
  WT.1 <- WORKING.DATA[WT.STRAINS,]
  NEG.1 <- subset(WORKING.DATA, STRAIN == NEG.STRAIN)
  for (i in I) {
    WORKING.DATA[i,"YFP.MEDIAN.ADJUST"] <- WORKING.DATA[i,"YFP.MEDIAN.CORRECT"] - mean(NEG.1[,"YFP.MEDIAN.CORRECT"])
    WORKING.DATA[i,"YFP.SD.ADJUST"] <- WORKING.DATA[i,"YFP.SD.CORRECT"] - mean(NEG.1[,"YFP.SD.CORRECT"])
  }
  
  #Relative Values
  WT.1 <- WORKING.DATA[WT.STRAINS,]
  NEG.1 <- subset(WORKING.DATA, STRAIN == NEG.STRAIN)
  for (i in 1:nrow(WORKING.DATA)) {
    WORKING.DATA[i,"FSC.MEDIAN.RELATIVE"] <- WORKING.DATA[i,"FSC.MEDIAN.CORRECT"]/mean(WT.1[,"FSC.MEDIAN.CORRECT"])
    WORKING.DATA[i,"YFP.MEDIAN.RELATIVE"] <- WORKING.DATA[i,"YFP.MEDIAN.ADJUST"]/mean(WT.1[,"YFP.MEDIAN.ADJUST"])
    WORKING.DATA[i,"YFP.SD.SCALED"] <- WORKING.DATA[i,"YFP.SD.ADJUST"]/mean(WT.1[,"YFP.MEDIAN.ADJUST"])
  }
  
  #Alternative Noise Metrics
  WT.1 <- WORKING.DATA[WT.STRAINS,]
  NEG.1 <- subset(WORKING.DATA, STRAIN == NEG.STRAIN)
  for (i in 1:nrow(WORKING.DATA)) {
    WORKING.DATA[i,"YFP.CV"] <- WORKING.DATA[i,"YFP.SD.SCALED"]/WORKING.DATA[i,"YFP.MEDIAN.RELATIVE"]
    WORKING.DATA[i,"YFP.FANO"] <- WORKING.DATA[i,"YFP.SD.SCALED"]^2/WORKING.DATA[i,"YFP.MEDIAN.RELATIVE"]
  }
  
  #Relative Alternative Noise Metrics
  WT.1 <- WORKING.DATA[WT.STRAINS,]
  NEG.1 <- subset(WORKING.DATA, STRAIN == NEG.STRAIN)
  for (i in 1:nrow(WORKING.DATA)) {
    WORKING.DATA[i,"YFP.SD.RELATIVE"] <- WORKING.DATA[i,"YFP.SD.SCALED"]/mean(WT.1[,"YFP.SD.SCALED"])
    WORKING.DATA[i,"YFP.CV.RELATIVE"] <- WORKING.DATA[i,"YFP.SD.RELATIVE"]/WORKING.DATA[i,"YFP.MEDIAN.RELATIVE"]
    WORKING.DATA[i,"YFP.FANO.RELATIVE"] <- WORKING.DATA[i,"YFP.SD.RELATIVE"]^2/WORKING.DATA[i,"YFP.MEDIAN.RELATIVE"]
  }
  
  #Add NA for samples where CV or FANO are undefined.
  WT.1 <- WORKING.DATA[WT.STRAINS,]
  NEG.1 <- subset(WORKING.DATA, STRAIN == NEG.STRAIN)
  for (i in 1:nrow(WORKING.DATA)) {
    if (WORKING.DATA[i,"YFP.MEDIAN.ADJUST"] < 0 | WORKING.DATA[i,"YFP.SD.ADJUST"] < 0) {
      WORKING.DATA[i,"YFP.CV"] <- NA
      WORKING.DATA[i,"YFP.CV.RELATIVE"] <- NA
      WORKING.DATA[i,"YFP.FANO"] <- NA
      WORKING.DATA[i,"YFP.FANO.RELATIVE"] <- NA		
    }
  }
  
  #Write processed data to file
  write.table(WORKING.DATA,paste("ALL.",DATA.NAME,".DATA.txt",sep=""),sep="\t",quote=FALSE,row.names=FALSE)

  #Complete Cases
  WORKING.DATA <- WORKING.DATA[complete.cases(WORKING.DATA),]
  
  #Remove FSC Outliers
  for (i in 1:nrow(WORKING.DATA)){
    CUR <- subset(WORKING.DATA, PLATE == WORKING.DATA[i,"PLATE"] & POSITION == WORKING.DATA[i,"POSITION"])
    LOW <- median(CUR$FSC.MEDIAN.RELATIVE) - 6*mad(CUR$FSC.MEDIAN.RELATIVE)
    HIGH <- median(CUR$FSC.MEDIAN.RELATIVE) + 6*mad(CUR$FSC.MEDIAN.RELATIVE)
    if (WORKING.DATA[i,"FSC.MEDIAN.RELATIVE"] > LOW & WORKING.DATA[i,"FSC.MEDIAN.RELATIVE"] < HIGH) {
      WORKING.DATA[i,"FSC.OUTLIER"] <- "NO"
    } else {
      WORKING.DATA[i,"FSC.OUTLIER"] <- "YES"
    }
  }
  
  #Remove YFP Outliers
  for (i in 1:nrow(WORKING.DATA)) {
    CUR <- subset(WORKING.DATA, PLATE == WORKING.DATA[i,"PLATE"] & POSITION == WORKING.DATA[i,"POSITION"])
    LOW <- median(CUR$YFP.MEDIAN.RELATIVE) - 6*mad(CUR$YFP.MEDIAN.RELATIVE)
    HIGH <- median(CUR$YFP.MEDIAN.RELATIVE) + 6*mad(CUR$YFP.MEDIAN.RELATIVE)
    if (WORKING.DATA[i,"YFP.MEDIAN.RELATIVE"] > LOW & WORKING.DATA[i,"YFP.MEDIAN.RELATIVE"] < HIGH) {
      WORKING.DATA[i,"YFP.OUTLIER"] <- "NO"
    } else {
      WORKING.DATA[i,"YFP.OUTLIER"] <- "YES"
    }
  }

  FILTER.WORKING.DATA <- subset(WORKING.DATA, FSC.OUTLIER == "NO" & YFP.OUTLIER == "NO")
  
  write.table(FILTER.WORKING.DATA,paste("FILTER.",DATA.NAME,".DATA.txt",sep=""),sep="\t",quote=FALSE,row.names=FALSE)

  #Summary Stats
  WORKING.MEDIAN <- aggregate(cbind(FSC.MEDIAN.RELATIVE, YFP.MEDIAN.RELATIVE, YFP.SD.SCALED, YFP.CV, YFP.FANO, YFP.SD.RELATIVE, YFP.CV.RELATIVE, YFP.FANO.RELATIVE) ~ STRAIN + PLATE + POSITION, data=FILTER.WORKING.DATA, FUN = median)
  WORKING.SD <- aggregate(cbind(FSC.MEDIAN.RELATIVE, YFP.MEDIAN.RELATIVE, YFP.SD.SCALED, YFP.CV, YFP.FANO, YFP.SD.RELATIVE, YFP.CV.RELATIVE, YFP.FANO.RELATIVE) ~ STRAIN + PLATE + POSITION, data=FILTER.WORKING.DATA, FUN = sd)
  WORKING <- aggregate(cbind(FSC.MEDIAN.RELATIVE, YFP.MEDIAN.RELATIVE, YFP.SD.SCALED, YFP.CV, YFP.FANO, YFP.SD.RELATIVE, YFP.CV.RELATIVE, YFP.FANO.RELATIVE) ~ STRAIN + PLATE + POSITION, data=FILTER.WORKING.DATA, FUN = length)
  
  WORKING.MEDIAN <- WORKING.MEDIAN[which(WORKING[,"YFP.MEDIAN.RELATIVE"] >= 2),]
  WORKING.SD <- WORKING.SD[which(WORKING[,"YFP.MEDIAN.RELATIVE"] >= 2),]
  WORKING <- WORKING[which(WORKING[,"YFP.MEDIAN.RELATIVE"] >= 2),]
  
  WORKING.SUMMARY <- cbind.data.frame(WORKING.MEDIAN,WORKING.SD[,4:11],WORKING[,11])
  colnames(WORKING.SUMMARY) <- c("STRAIN","PLATE","POSITION","FSC.MEDIAN.RELATIVE.MEAN","YFP.MEDIAN.RELATIVE.MEAN","YFP.SD.SCALED.MEAN","YFP.CV.MEAN","YFP.FANO.MEAN","YFP.SD.RELATIVE.MEAN","YFP.CV.RELATIVE.MEAN","YFP.FANO.RELATIVE.MEAN","FSC.MEDIAN.RELATIVE.SD","YFP.MEDIAN.RELATIVE.SD","YFP.SD.SCALED.SD","YFP.CV.SD","YFP.FANO.SD","YFP.SD.RELATIVE.SD","YFP.CV.RELATIVE.SD","YFP.FANO.RELATIVE.SD","N")
  
  write.table(WORKING.SUMMARY,paste("SUMMARY.",DATA.NAME,".DATA.txt",sep=""),sep="\t",quote=FALSE,row.names=FALSE)
  
  #PERMUTATION TESTs
  ALL.REP <- FILTER.WORKING.DATA
  SUMMARY <- WORKING.SUMMARY
  
  WT <- ALL.REP[WT.STRAINS,]
  
  #FSC
  #Create list of WT phenotype grouped by strain
  N.WT <- length(WT)
  WT.LIST <- vector('list',N.WT)
  for (i in 1:N.WT)
  {
    CUR.WT <- subset(WT, PLATE == SUMMARY[WT.STRAINS[i],"PLATE"] & POSITION == SUMMARY[WT.STRAINS[i],"POSITION"])
    WT.LIST[[i]] <- CUR.WT$FSC.MEDIAN.RELATIVE
  }
  
  #Create list of all strain phenotypes grouped by strain
  SUMMARY <- SUMMARY[order(SUMMARY$STRAIN),]
  SUMMARY <- droplevels(SUMMARY)
  ALL.REP <- droplevels(subset(ALL.REP,STRAIN %in% as.character(SUMMARY$STRAIN)))
  ALL.REP <- ALL.REP[order(ALL.REP$STRAIN),]
  N.STRAIN <- nrow(SUMMARY)
  STRAIN.LIST <- vector('list',N.STRAIN)
  for (i in 1:N.STRAIN)
  {
    CUR.STRAIN <- subset(ALL.REP, PLATE == SUMMARY[i,"PLATE"] & POSITION == SUMMARY[i,"POSITION"])
    STRAIN.LIST[[i]] <- CUR.STRAIN$FSC.MEDIAN.RELATIVE
  }
  
  #Permutation function
  SHUFFLE <- function(STRAIN.LIST = STRAIN.LIST[[i]],WT.LIST = WT.LIST) {
    SAMPLE <- c(STRAIN.LIST,WT.LIST[[sample(1:length(WT.LIST),1)]])
    OUTPUT <- sample(SAMPLE,length(SAMPLE))
    DIST.SIM <- abs(mean(OUTPUT[1:length(STRAIN.LIST)]) - mean(OUTPUT[(length(STRAIN.LIST)+1):length(SAMPLE)]))
    DIST.OBS <- abs(mean(SAMPLE[1:length(STRAIN.LIST)]) - mean(SAMPLE[(length(STRAIN.LIST)+1):length(SAMPLE)]))
    DIFF <- DIST.OBS - DIST.SIM
    return(DIFF)
  }
  
  for (i in 1:length(STRAIN.LIST))
  {
    DIFF <- sapply(1:N.PERM,FUN=function(x)	SHUFFLE(STRAIN.LIST = STRAIN.LIST[[i]],WT.LIST = WT.LIST))
    P.VAL <- (length(which(DIFF < 0)) + 1) / (length(which(DIFF < 0)) + length(which(DIFF > 0)) + 1) 	
    SUMMARY[i,"P.VAL.FSC"] <- P.VAL 
    print(round(i/length(STRAIN.LIST)*100,1))
  }
  
  #MEDIAN YFP
  ALL.REP <- FILTER.WORKING.DATA
  WT <- ALL.REP[WT.STRAINS,]
  
  #Create list of WT phenotype grouped by strain
  N.WT <- length(WT.STRAINS)
  WT.LIST <- vector('list',N.WT)
  for (i in 1:N.WT)
  {
    CUR.WT <- subset(WT, PLATE == SUMMARY[WT.STRAINS[i],"PLATE"] & POSITION == SUMMARY[WT.STRAINS[i],"POSITION"])
    WT.LIST[[i]] <- CUR.WT$YFP.MEDIAN.RELATIVE
  }
  
  #Create list of all strain phenotypes grouped by strain
  SUMMARY <- SUMMARY[order(SUMMARY$STRAIN),]
  SUMMARY <- droplevels(SUMMARY)
  ALL.REP <- droplevels(subset(ALL.REP,STRAIN %in% as.character(SUMMARY$STRAIN)))
  ALL.REP <- ALL.REP[order(ALL.REP$STRAIN),]
  N.STRAIN <- nrow(SUMMARY)
  STRAIN.LIST <- vector('list',N.STRAIN)
  for (i in 1:N.STRAIN)
  {
    CUR.STRAIN <- subset(ALL.REP, PLATE == SUMMARY[i,"PLATE"] & POSITION == SUMMARY[i,"POSITION"])
    STRAIN.LIST[[i]] <- CUR.STRAIN$YFP.MEDIAN.RELATIVE
  }
  
  for (i in 1:length(STRAIN.LIST))
  {
    DIFF <- sapply(1:N.PERM,FUN=function(x)	SHUFFLE(STRAIN.LIST = STRAIN.LIST[[i]],WT.LIST = WT.LIST))
    P.VAL <- (length(which(DIFF < 0)) + 1) / (length(which(DIFF < 0)) + length(which(DIFF > 0)) + 1) 	
    SUMMARY[i,"P.VAL.MEDIAN"] <- P.VAL 
    print(round(i/length(STRAIN.LIST)*100,1))
  }
  
  #NOISE YFP
  ALL.REP <- FILTER.WORKING.DATA
  WT <- ALL.REP[WT.STRAINS,]
  
  #Create list of WT phenotype grouped by strain
  N.WT <- length(WT.STRAINS)
  WT.LIST <- vector('list',N.WT)
  for (i in 1:N.WT)
  {
    CUR.WT <- subset(WT, PLATE == SUMMARY[WT.STRAINS[i],"PLATE"] & POSITION == SUMMARY[WT.STRAINS[i],"POSITION"])
    WT.LIST[[i]] <- CUR.WT$YFP.CV.RELATIVE
  }
  
  #Create list of all strain phenotypes grouped by strain
  SUMMARY <- SUMMARY[order(SUMMARY$STRAIN),]
  SUMMARY <- droplevels(SUMMARY)
  ALL.REP <- droplevels(subset(ALL.REP,STRAIN %in% as.character(SUMMARY$STRAIN)))
  ALL.REP <- ALL.REP[order(ALL.REP$STRAIN),]
  N.STRAIN <- nrow(SUMMARY)
  STRAIN.LIST <- vector('list',N.STRAIN)
  for (i in 1:N.STRAIN)
  {
    CUR.STRAIN <- subset(ALL.REP, PLATE == SUMMARY[i,"PLATE"] & POSITION == SUMMARY[i,"POSITION"])
    STRAIN.LIST[[i]] <- CUR.STRAIN$YFP.CV.RELATIVE
  }
  
  for (i in 1:length(STRAIN.LIST))
  {
    DIFF <- sapply(1:N.PERM,FUN=function(x)	SHUFFLE(STRAIN.LIST = STRAIN.LIST[[i]],WT.LIST = WT.LIST))
    P.VAL <- (length(which(DIFF < 0)) + 1) / (length(which(DIFF < 0)) + length(which(DIFF > 0)) + 1) 	
    SUMMARY[i,"P.VAL.NOISE"] <- P.VAL 
    print(round(i/length(STRAIN.LIST)*100,1))
  }
  
  write.table(SUMMARY,paste("SUMMARY.",DATA.NAME,".P.VAL.txt",sep=""),sep="\t",quote=FALSE,row.names=FALSE)
}

#--------------------------------------------------------------------------------------------------------------------------------#

  ########################
### 7 - GEOMETRIC AVERAGE###
  ########################

GEOMEAN <- function(x, na.rm = FALSE, trim = 0, ...){exp(mean(log(x, ...), na.rm = na.rm, trim = trim, ...))}

#--------------------------------------------------------------------------------------------------------------------------------#

  ###################################
### 8 - GEOMETRIC STANDARD DEVIATION###
  ###################################

GEOSD <- function(x, na.rm = FALSE, ...){exp(sd(log(x, ...), na.rm = na.rm, ...))}

#--------------------------------------------------------------------------------------------------------------------------------#

  ########################
### 9 - MUTATION DENSITY ###
  ########################
#Calculate density and probability from arbitrary distribution (base.dist)
#Assumes that density can be modeled as sum of N(mu,sigma) 
#mu = estimate of mean of each mutant (uses median)
#sigma = estimate of sd of each mutant (uses MAD)
#Default base.dist is the trans mutation data

DMUTATION.MEAN <- function(x, base.dist = TRANS.STRAIN.2$MEAN) {
  #Calculate density at each point specified
  I <- 1:length(x)
  #Normalization constant for using MAD over SD
  k <- 1/qnorm(.75)
  #Set up matrix of results
  M <- matrix(0,nrow=length(base.dist),ncol=length(I))
  #Uses median MAD estimated from mutational data
  MEDIAN.MAD <- median(TRANS.STRAIN.2$MEAN.MAD)
  for(i in I){
    M[,i] <- dnorm(x[i],mean=base.dist,sd=k*MEDIAN.MAD)
  }
  #Output is density for each of the specified values (x)
  return(colSums(M)/length(base.dist))
}

#--------------------------------------------------------------------------------------------------------------------------------#

  #####################
### 10 - eQTL Overlap ###
  #####################
overlap <- function(test.data) {
  #Count number of single peaks, peaks shared among two strains, or peaks shared among all three strains
  SINGLES <- c(0,0,0)
  DOUBLES <- c(0,0,0,0,0)
  TRIPLES <- 0
  
  #Collect all overlaps
  #Peaks overlap if their CIs overlap
  I <- 1:nrow(test.data)
  OVERLAP.MATRIX <- matrix(1,nrow=length(I),ncol=length(I))
  for(i in I) {
    OUT <- which(test.data$CHR != test.data$CHR[i] | (test.data$POS.MAX < test.data$POS.MIN[i] | test.data$POS.MIN > test.data$POS.MAX[i])) 
    OVERLAP.MATRIX[i,OUT] <- 0
  }

  #Resolve overlaps into discrete sets
  MERGE <- test.data
  MERGE$SET <- 0
  SITE <- 1
  X <- 1
  SET <- 1
  OVERLAP.SET <- list()
  while(SITE <= nrow(MERGE)) {
    if(SITE < nrow(MERGE)) {
      while(sum(OVERLAP.MATRIX[SITE:(SITE+X),SITE:(SITE+X)]) >= 0.6*(X+1)^2) {
        if(SITE + X + 1 <= nrow(MERGE)) {
          X <- X + 1
        } else {
          X <- X + 1
          break
        }
      }
    }
    X <- X - 1
    if(X == 0){
      MERGE$SET[SITE] <- SET
      OVERLAP.SET[[SET]] <- MERGE[SITE,]
    } else {
      MERGE$SET[SITE:(SITE+X)] <- SET
      OVERLAP.SET[[SET]] <- MERGE[SITE:(SITE+X),]
    }
    SITE <- SITE + X + 1
    SET <- SET + 1 
    X <- 1
  }    
  
  #Divide each discrete set into its overlap components    
  K <- 1:length(OVERLAP.SET)
  for(k in K) {
    if(nrow(OVERLAP.SET[[k]]) == 1) {
      SINGLES[which(levels(test.data$SAMPLE.STRAIN) %in% OVERLAP.SET[[k]]$SAMPLE.STRAIN)] <- SINGLES[which(levels(test.data$SAMPLE.STRAIN) %in% OVERLAP.SET[[k]]$SAMPLE.STRAIN)] + 1
    } else if(nrow(OVERLAP.SET[[k]]) == 2) {
      DOUBLES[sum(which(levels(test.data$SAMPLE.STRAIN) %in% OVERLAP.SET[[k]]$SAMPLE.STRAIN))] <- DOUBLES[sum(which(levels(test.data$SAMPLE.STRAIN) %in% OVERLAP.SET[[k]]$SAMPLE.STRAIN))] + 1
    } else if(nrow(OVERLAP.SET[[k]]) == 3) {
        if(length(which(levels(test.data$SAMPLE.STRAIN) %in% OVERLAP.SET[[k]]$SAMPLE.STRAIN)) == 3) {
          TRIPLES <- TRIPLES + 1
        } else {
          DOUBLES[sum(which(levels(test.data$SAMPLE.STRAIN) %in% OVERLAP.SET[[k]]$SAMPLE.STRAIN))] <- DOUBLES[sum(which(levels(test.data$SAMPLE.STRAIN) %in% OVERLAP.SET[[k]]$SAMPLE.STRAIN))] + 1
          SINGLES[which(table(OVERLAP.SET[[k]]$SAMPLE.STRAIN) == 2)] <- SINGLES[which(table(OVERLAP.SET[[k]]$SAMPLE.STRAIN) == 2)] + 1
        }
    } else if(nrow(OVERLAP.SET[[k]]) == 4) {
      if(length(which(levels(test.data$SAMPLE.STRAIN) %in% OVERLAP.SET[[k]]$SAMPLE.STRAIN)) == 3) {
        TRIPLES <- TRIPLES + 1
        SINGLES[which(table(OVERLAP.SET[[k]]$SAMPLE.STRAIN) == 2)] <- SINGLES[which(table(OVERLAP.SET[[k]]$SAMPLE.STRAIN) == 2)] + 1
      } else {
        DOUBLES[sum(which(table(OVERLAP.SET[[k]]$SAMPLE.STRAIN) == 2))] <- DOUBLES[sum(which(table(OVERLAP.SET[[k]]$SAMPLE.STRAIN) == 2))] + 2
      }
    } else if(nrow(OVERLAP.SET[[k]]) == 5) {
      if(length(which(levels(test.data$SAMPLE.STRAIN) %in% OVERLAP.SET[[k]]$SAMPLE.STRAIN)) == 3) {
        TRIPLES <- TRIPLES + 1
        DOUBLES[sum(which(table(OVERLAP.SET[[k]]$SAMPLE.STRAIN) == 2))] <- DOUBLES[sum(which(table(OVERLAP.SET[[k]]$SAMPLE.STRAIN) == 2))] + 1
      } else {
        DOUBLES[sum(which(table(OVERLAP.SET[[k]]$SAMPLE.STRAIN) == 2))] <- DOUBLES[sum(which(table(OVERLAP.SET[[k]]$SAMPLE.STRAIN) == 2))] + 2
        SINGLES[which(table(OVERLAP.SET[[k]]$SAMPLE.STRAIN) == 3)] <- SINGLES[which(table(OVERLAP.SET[[k]]$SAMPLE.STRAIN) == 3)] + 1
      }
    }
  }
  OUTPUT <- list(MERGE,SINGLES,DOUBLES,TRIPLES)
  names(OUTPUT) <- c("SETS","SINGLES","DOUBLES","TRIPLES")
  return(OUTPUT)
}

#--------------------------------------------------------------------------------------------------------------------------------#

  #########################################
### 11 - eQTL Overlap Significance Testing###
  #########################################
overlap.test <- function(test.data, N) {
  I <- 1:N
  FRAC.OVERLAP <- numeric(length(I))
  for(i in I){
    S1 <- subset(test.data, test.data$SAMPLE.STRAIN == levels(test.data$SAMPLE.STRAIN)[1])
    S2 <- subset(test.data, test.data$SAMPLE.STRAIN == levels(test.data$SAMPLE.STRAIN)[2])
    S3 <- subset(test.data, test.data$SAMPLE.STRAIN == levels(test.data$SAMPLE.STRAIN)[3])
  
    S1.PERMUTE <- S1
    S2.PERMUTE <- S2
    S3.PERMUTE <- S3
  
    #Get distances of intervals
    S1.DISTANCE <- S1$POS.MAX-S1$POS.MIN
    S2.DISTANCE <- S2$POS.MAX-S2$POS.MIN
    S3.DISTANCE <- S3$POS.MAX-S3$POS.MIN
  
    #Get distance of peak from min
    S1.PEAK.DIST <- S1$POS.PEAK - S1$POS.MIN
    S2.PEAK.DIST <- S2$POS.PEAK - S2$POS.MIN
    S3.PEAK.DIST <- S3$POS.PEAK - S3$POS.MIN
  
    #Sample chromosomes according to length
    S1.CHR <- sample(1:16, length(S1.DISTANCE), prob=CHROMOSOMES$LENGTH/sum(CHROMOSOMES$LENGTH),replace=TRUE)
    S2.CHR <- sample(1:16, length(S2.DISTANCE), prob=CHROMOSOMES$LENGTH/sum(CHROMOSOMES$LENGTH),replace=TRUE)
    S3.CHR <- sample(1:16, length(S3.DISTANCE), prob=CHROMOSOMES$LENGTH/sum(CHROMOSOMES$LENGTH),replace=TRUE)

    S1.PERMUTE$CHR <- S1.CHR
    S2.PERMUTE$CHR <- S2.CHR
    S3.PERMUTE$CHR <- S3.CHR
  
    #Sample non-overlapping intervals for each chromosome
    for(j in J) {
      S1.LENGTH <- sum(S1.DISTANCE[S1.CHR == j])
      S1.SAMPLE <- sample(1:(CHROMOSOMES$LENGTH[j] - S1.LENGTH), length(S1.DISTANCE[S1.CHR == j]))
      S1.SAMPLE <- S1.SAMPLE[order(S1.SAMPLE)]
      S1.MAX <- S1.SAMPLE + cumsum(S1.DISTANCE[S1.CHR == j])
      S1.MIN <- S1.MAX - S1.DISTANCE[S1.CHR == j]
      S1.PERMUTE$POS.MIN[S1.CHR==j] <- S1.MIN
      S1.PERMUTE$POS.MAX[S1.CHR==j] <- S1.MAX
      S1.PERMUTE$POS.PEAK[S1.CHR==j] <- S1.MIN + S1.PEAK.DIST[which(S1.CHR==j)]
      
      S2.LENGTH <- sum(S2.DISTANCE[S2.CHR == j])
      S2.SAMPLE <- sample(1:(CHROMOSOMES$LENGTH[j] - S2.LENGTH), length(S2.DISTANCE[S2.CHR == j]))
      S2.SAMPLE <- S2.SAMPLE[order(S2.SAMPLE)]
      S2.MAX <- S2.SAMPLE + cumsum(S2.DISTANCE[S2.CHR == j])
      S2.MIN <- S2.MAX - S2.DISTANCE[S2.CHR == j]
      S2.PERMUTE$POS.MIN[which(S2.CHR==j)] <- S2.MIN
      S2.PERMUTE$POS.MAX[which(S2.CHR==j)] <- S2.MAX
      S2.PERMUTE$POS.PEAK[which(S2.CHR==j)] <- S2.MIN + S2.PEAK.DIST[which(S2.CHR==j)]

      S3.LENGTH <- sum(S3.DISTANCE[S3.CHR == j])
      S3.SAMPLE <- sample(1:(CHROMOSOMES$LENGTH[j] - S3.LENGTH), length(S3.DISTANCE[S3.CHR == j]))
      S3.SAMPLE <- S3.SAMPLE[order(S3.SAMPLE)]
      S3.MAX <- S3.SAMPLE + cumsum(S3.DISTANCE[S3.CHR == j])
      S3.MIN <- S3.MAX - S3.DISTANCE[S3.CHR == j]
      S3.PERMUTE$POS.MIN[which(S3.CHR==j)] <- S3.MIN
      S3.PERMUTE$POS.MAX[which(S3.CHR==j)] <- S3.MAX
      S3.PERMUTE$POS.PEAK[which(S3.CHR==j)] <- S3.MIN + S3.PEAK.DIST[which(S3.CHR==j)]
    }
  
    MERGE <- rbind(S1.PERMUTE,S2.PERMUTE,S3.PERMUTE)
    MERGE <- MERGE[order(MERGE$CHR,MERGE$POS.MIN),]

    OVERLAP <- overlap(MERGE)
  
    FRAC.OVERLAP[i] <- (sum(OVERLAP$DOUBLES) + OVERLAP$TRIPLES)/(sum(OVERLAP$SINGLES) + sum(OVERLAP$DOUBLES) + OVERLAP$TRIPLES)
  }
  return(FRAC.OVERLAP)
}

#--------------------------------------------------------------------------------------------------------------------------------#
```
